# Myelin Supports Cortical Circuit Function Underlying Skilled Movement

**DOI:** 10.64898/2025.12.23.696289

**Authors:** Kimberly Gagnon, Gustavo Della Flora Nunes, Dailey Nettles, Thomas Nguyen, Elise R. Carter, Andreza Lins, Ryan Williamson, Jeremie Lefebvre, Daniel Denman, Ethan G. Hughes, Cristin G. Welle

## Abstract

Primary motor cortex (M1) is among the most heavily myelinated cortical regions and generates tightly coordinated neuronal activity patterns that drive skilled movement. Activity-dependent myelination is required for motor skill acquisition, and myelin loss in demyelinating diseases such as multiple sclerosis leads to motor impairment. Yet how myelination influences neuronal activity underlying skilled behavior remains unclear. By combining *in vivo* imaging of oligodendrocytes with high density Neuropixels recordings during dexterous reaching, we demonstrate that cuprizone-induced demyelination impairs movement efficiency, and alters cell-type–specific neuronal activity and synchrony in a manner that predicts motor output. Using a computational model constrained by these data, we identify inhibitory axonal propagation failures as a mechanistic link between myelin loss and altered circuit function. Partial remyelination normalizes cortical network-level metrics and reach consistency but leaves smooth movement impaired, revealing a selective vulnerability in inhibitory circuits. These findings close a critical gap between cellular models of demyelination and clinical motor impairment by demonstrating how myelin supports cortical circuit dynamics driving skilled behavior.

---

Skilled movement depends on precisely coordinated neural activity within the primary motor cortex (M1), which is among the most heavily myelinated cortical regions1–4. This reproducible pattern of neural activity emerges with learning and supports the execution of stable, consistent motor output4,5. Although myelin is classically viewed as a fixed substrate for rapid conduction, oligodendrocytes and their precursors are now known to undergo activity-dependent remodeling, linking neuronal activity to adaptive changes in myelin6–8. This plasticity supports the consolidation of learned movement9, positioning myelination as an active regulator of motor circuit function. Yet, how myelin shapes the cortical population dynamics that support the execution of skilled movement remains poorly understood. The consequences of demyelination in both clinical disorders and experimental models underscore a critical role for myelin in maintaining motor output. In multiple sclerosis (MS), a demyelinating disorder of the central nervous system, demyelination produces motor impairments ranging from subtle deficits in stable movement to profound disability10,11. Although axons can be remyelinated after injury, repair is often incomplete and heterogenous-raising the possibility that partial remyelination may restore some aspects of circuit function while leaving others impaired8,12,13. However, concurrent autoimmunity, axonal/synaptic degeneration, and chronicity that frequently accompany demyelination in both clinical populations and many experimental models, make isolating acute, myelin-specific effects on motor function difficult to ascertain. Previously we found that acute treatment with cuprizone, a model of toxin-induced demyelination that avoids autoimmunity, axonal loss, and spinal pathology13–15 shows that myelin loss alone induces M1 hyperexcitability and impairs learning of skilled reach, with neuronal activity and behavior improving after remyelination8. These data highlight a myelin-specific contribution to cortical circuit function and aligns with evidence that cortical myelin also supports neuronal excitability, metabolic demand, and action potential propagation fidelity16–18. Among these locally myelinated populations, parvalbumin interneurons appear especially dependent on intact myelin to sustain high-frequency firing17, which in turn is essential for stabilizing the activity of principal excitatory neurons driving behavior19. These findings broaden the classical view of myelin as merely a substrate for rapid long-range conduction to an active regulator of cortical computation. Therefore, understanding how demyelination disrupts cell type-specific dynamics across cortical layers during movement- and how these dynamics recover with remyelination- is critical for understanding the circuit principles through which myelin enables motor output.

To understand how myelin supports cortical circuit dynamics underlying skilled movement, we combine longitudinal in vivo imaging of oligodendrocytes and myelin with Neuropixels recordings of cortical activity during skilled reach in the same animal. We show that myelin loss degrades movement smoothness and consistency, selectively disrupts deep-layer excitatory and inhibitory recruitment, and increases fast-timescale synchrony in a manner that predicts movement variability. We further use a computational model of local circuit dynamics constrained by these data to demonstrate that modest reductions in inhibitory spike propagation fidelity are sufficient to produce the observed hyperexcitability and hypersynchrony. Finally, we show that partial remyelination restores network-level properties and movement consistency but leaves movement smoothness impaired. These findings reveal how myelin shapes cortical dynamics during behavior and identify inhibitory fidelity as a critical substrate of precise myelin-dependent motor execution.

## Results

### Longitudinal Two-Photon Imaging of Oligodendrocytes Reveals Extent of Demyelination in Primary Motor Cortex

To determine how myelin shapes neural activity underlying skilled movement, we combined longitudinal two-photon imaging of oligodendrocytes and myelin in forelimb M1 with high-density Neuropixels recordings and kinematic analysis during a head-fixed dexterous reach-to-grasp task (**Fig. 1A,B**). To monitor oligodendrocytes and myelin in M1 across time, we used *Mobp-EGFP* mice, which label all myelinating oligodendrocytes and sheaths. Mice between 6 and 8 weeks old were implanted with a cranial window centered over the caudal-forelimb area (CFA) of M1 and trained to expert levels in a head-fixed dexterous forelimb reaching task (**Methods**). Following the end of the training, mice underwent baseline imaging, were randomly assigned to demyelination or healthy cohorts and fed 0.2% cuprizone or normal chow for 3 weeks, respectively. Cuprizone treatment selectively ablates myelinating oligodendrocytes without autoimmunity, and this acute three-week treatment protocol preserves axonal and synaptic integrity (**Figure S1A,B**), enabling measurement of circuit changes downstream of myelin loss. The dynamics of the loss and generation of oligodendrocytes and myelin was monitored by *in vivo* two-photon imaging of the same region within M1 in individual mice through time (**Figure 1C**). Four days after removal of cuprizone treatment, mice exhibited a 55.1 ± 4.3% reduction in oligodendrocytes and generated only 9.29 ± 1.87% new cells (**Fig. 1D**). Immunohistochemistry confirmed a reduction in the density of both oligodendrocytes in superficial M1 (**Figures S1D,E**). Importantly, corresponding oligodendrocyte loss in deep layers was correlated to oligodendrocyte loss in superficial layers (**Figures SF-H**), similar to previous work^20^. Myelin content scales linearly with oligodendrocyte density across brain regions^21^, and we confirmed this relationship in M1 by histology, where we found a strong positive correlation between oligodendrocyte number and myelin levels in superficial cortex (**Figure S1I-J**). Since myelin loss precedes oligodendrocyte death in the cuprizone model^8,22^, we developed a computational approach to better estimate myelin levels from counts of oligodendrocytes derived from volumetric two-photon images. We quantified myelin sheath generation and loss in individual oligodendrocytes and extrapolated these dynamics to all cells within the imaged volume (**Fig. S2**; **Methods**). To validate this model, we manually traced all myelin sheaths in a small volume before and after cuprizone and found close agreement with our computational estimates (**Figures 1E–G**). Applying this approach to our dataset, revealed that myelin loss preceded oligodendrocyte loss by ~5 days (**Figure 1H**), similar to previous work^8^. Three days after cuprizone withdrawal, mice had lost 67.7 ± 4.7% of initial myelin with minimal new sheath formation (9.75 ± 2%), resulting in an overall 58 ± 3.5% reduction relative to baseline (**Figure 1I,J**). Thus, acute cuprizone treatment produces profound M1 demyelination, with myelin loss closely tracking-but preceding-oligodendrocyte loss.

**Figure 1.**
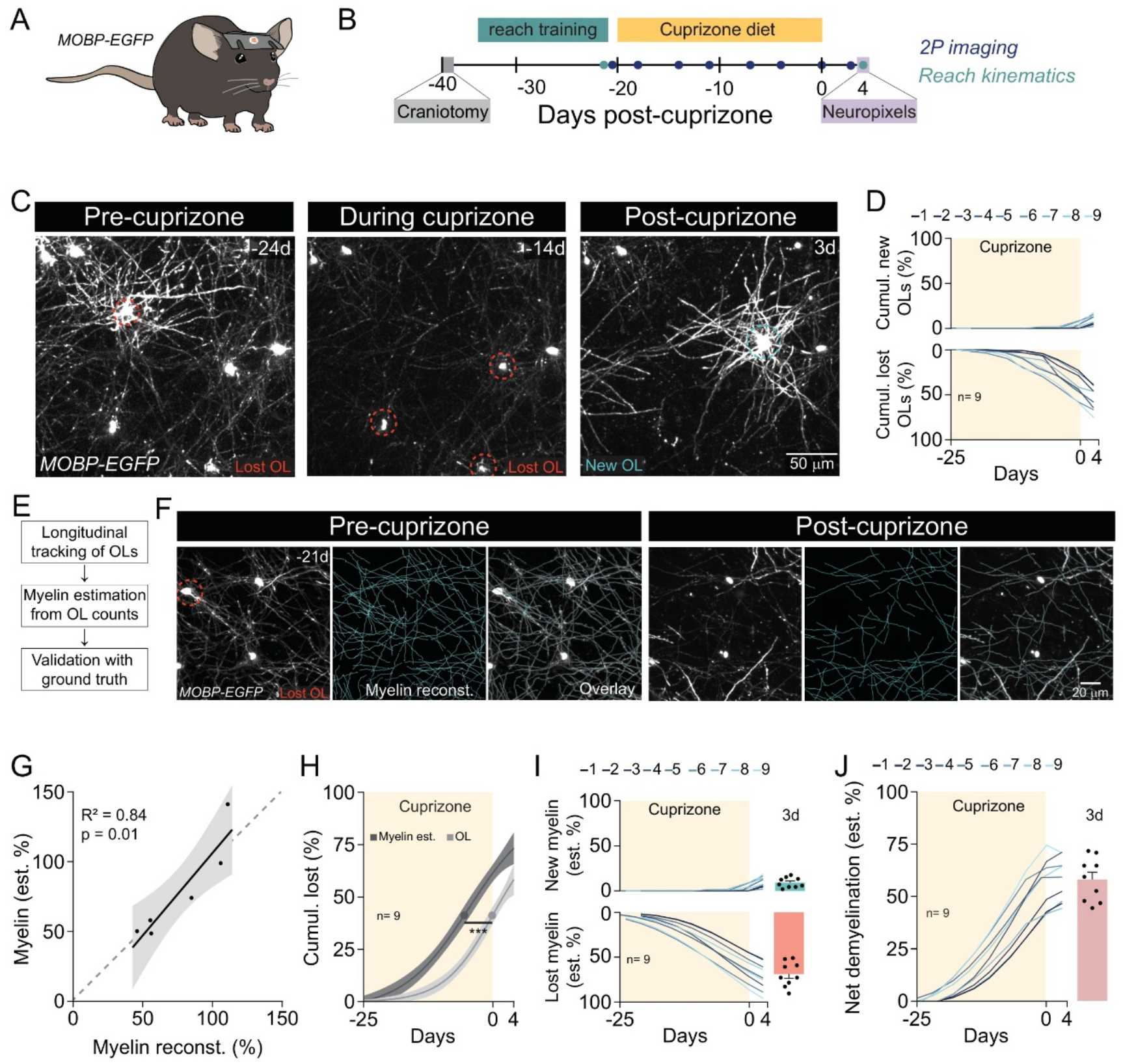
Longitudinal Two-Photon Imaging of Oligodendrocytes Reveals Extent of Demyelination in Primary Motor Cortex. **(A)** MOBP-EGFP mice were used for *in vivo* visualization of oligodendrocytes and myelin. **(B)** Experimental timeline: mice received a CFA-centered cranial window, were trained to expert performance on the head-fixed reach task, imaged at baseline, fed 0.2% cuprizone, and recorded with Neuropixels 4 days after cuprizone. **(C)** Representative two-photon images showing oligodendrocyte gain and loss. **(D)** Quantification of oligodendrocyte loss and replacement across time. **(E)** Validation of the myelin-estimation approach. **(F)** Manual myelin tracing from the same M1 region before and after cuprizone. **(G)** Estimated myelin levels strongly correlated with ground-truth reconstructions (n=3 mice; two post-cuprizone time points; myelin expressed as % of pre-cuprizone sheaths). **(H)** Modeled dynamics showed myelin loss preceded oligodendrocyte loss (paired t-test, t(8)=7.11, ***p<0.001, n=9). **(I)** Estimated myelin loss and gain for individual mice. **(J)** Overall demyelination dynamics combining sheath loss and regeneration.

### Demyelination severity in primary motor cortex impacts the execution of skilled movement

To evaluate the role of myelin in shaping motor output, we examined high-resolution kinematic features during expert rehearsal of the dexterous reach four days (4D) after cuprizone withdrawal in the same mice. Forelimb reach trajectories were tracked using custom code derived from DeepLabCut^23,24^, decomposed into X and Y components, and analyzed over the reach duration: reach initiation to maximum extension (reach max) (**Figure 2A**). Reach attempts and success rates following cuprizone treatments were comparable between expert healthy and demyelinated mice and did not vary with demyelination severity, indicating preserved task engagement (**Figures 2B; S3A,B**). Gross kinematic features-including reach duration, velocity, and pathlength- were also maintained (**Figure S3C–H**). Despite this preserved performance, demyelinated mice showed disorganized reach trajectories (**Figure 2C**). The consistency of reach events, measured as within-session pairwise reach correlations, were significantly reduced in cuprizone treated animals compared to healthy mice (**Figure 2D**; demyelinated: 0.89 ± 0.02; healthy: 0.94 ± 0.01), reflecting greater reach-to-reach variability. Importantly, reach consistency did not correlate with net demyelination, suggesting that movement consistency is not directly influenced by demyelination severity (**Figure S2I**; Pearson, R^2^ = 0.07, *p* = 0.57).

**Figure 2.**
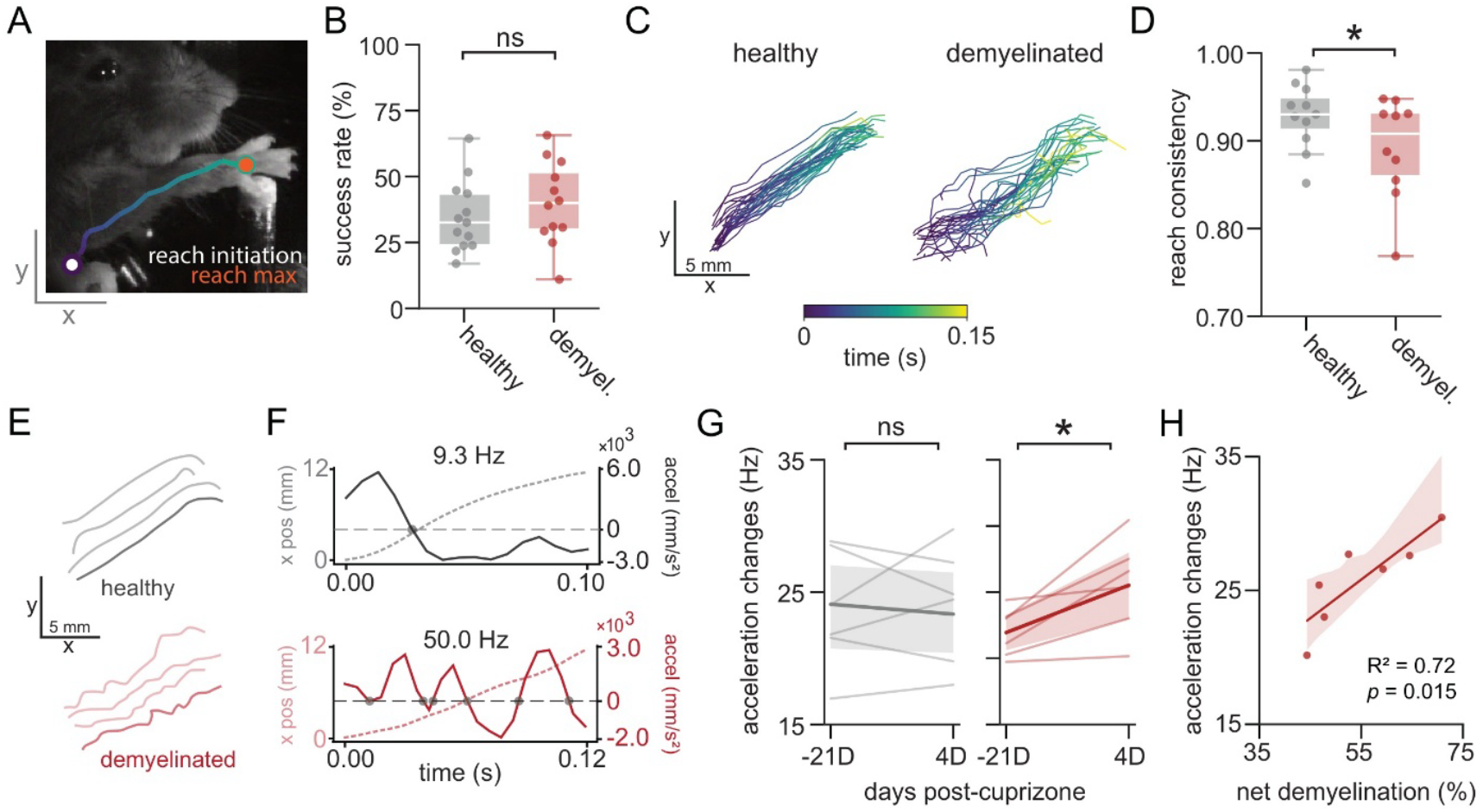
Demyelination severity in M1 impacts the execution of skilled movement. **(A)** Example video frame of a mouse performing the reach-to-grasp task, with trajectory from reach initiation (purple) to reach maximum (orange). **(B)** Post-cuprizone success rates did not differ between healthy and demyelinated mice (n=13 healthy; n=12 demyelinated; t=-1.2, p=0.241, t-test). **(C)** Representative reach trajectories from a healthy and a demyelinated mouse (first 30 reaches). **(D)** Reach consistency, measured as mean pairwise trajectory correlation, was reduced after demyelination (n=11 healthy; n=10 demyelinated; t=2.42, p=0.02, t-test; box plots represent median, interquartile range). **(E)** Representative reach trajectories from healthy (gray) and demyelinated (red) mice. **(F)** Example position (dashed) and acceleration (solid) traces from **(E)**, with zero crossings marked (gray dots). Healthy reaches showed smooth motion; demyelinated reaches exhibited high-frequency acceleration changes. **(G)** Acceleration changes increased after cuprizone in demyelinated but not in healthy mice (n=6 healthy mice, t=0.66, n=6 demyelinated mice, t=0.02, p=−3.24, paired t-test). Solid lines/shaded regions indicate mean±SEM, respectively. **(H)** Acceleration changes correlated with demyelination severity (n=7; R^2^=0.72, p=0.015), indicating reduced smooth movement with greater myelin loss. Boxplots represent median and interquartile range. Shaded regions denote 95% confidence intervals around linear regression fits. ns, not significant. *p < 0.05

Smooth ongoing movement is a hallmark of efficient motor control and can be quantified by measuring the rate of acceleration changes, measured as zero crossings in acceleration, during reach extension^25^. Higher rates of acceleration change indicate more frequent deviations from a stable trajectory^26^. We found that individual reaches made by demyelinated mice appeared less smooth during ongoing movement (**Figure 2E,F**). Before cuprizone treatment (−21D), both groups showed low rates of acceleration changes, which was consistent with smooth, efficient reaching (**Figure 2G**). Healthy mice showed no change in rates of acceleration (Δpost–pre = –0.75 Hz, p = 0.664), whereas demyelinated mice exhibited an increase in acceleration changes (Δpost–pre = +3.58 Hz, p = 0.023), reflecting impaired smooth movement relative to individual baselines (**Figure 2G**). Notably, rate of acceleration changes positively correlated with demyelination severity across mice (**Figure 2H**), indicating that greater myelin loss is associated with reduced smooth movement and degradation in skilled motor output.

### Neuronal circuit dysfunction scales with demyelination severity in primary motor cortex

Given that skilled movement depends on precisely coordinated neural activity^4^, we next asked whether myelin loss alters M1 cortical function underlying reach execution. To assess the role of myelin in shaping movement-related neural activity, we performed high-density Neuropixels recordings in caudal forelimb area (CFA) during reach execution in the same mice (**Figure 3A; Figure 1A**). A similar number of units were recorded in healthy and demyelinated mice (**Figure S4A**). Mean session-wide firing rates were modestly higher in demyelinated mice (healthy: 1.85 ± 0.11 Hz, demyelinated: 2.18 ± 0.17 Hz) but not significantly different (**Figure S4B,C**, REML, p=0.086). However, we found that mouse-averaged firing rates correlated with both demyelination severity (R^2^=0.81, p=0.002, Pearson) and reach acceleration changes (R^2^=0.72, p=0.006, Pearson), linking myelin-dependent M1 activity levels to movement quality (**Figure S4D,E**).

**Figure 3.**
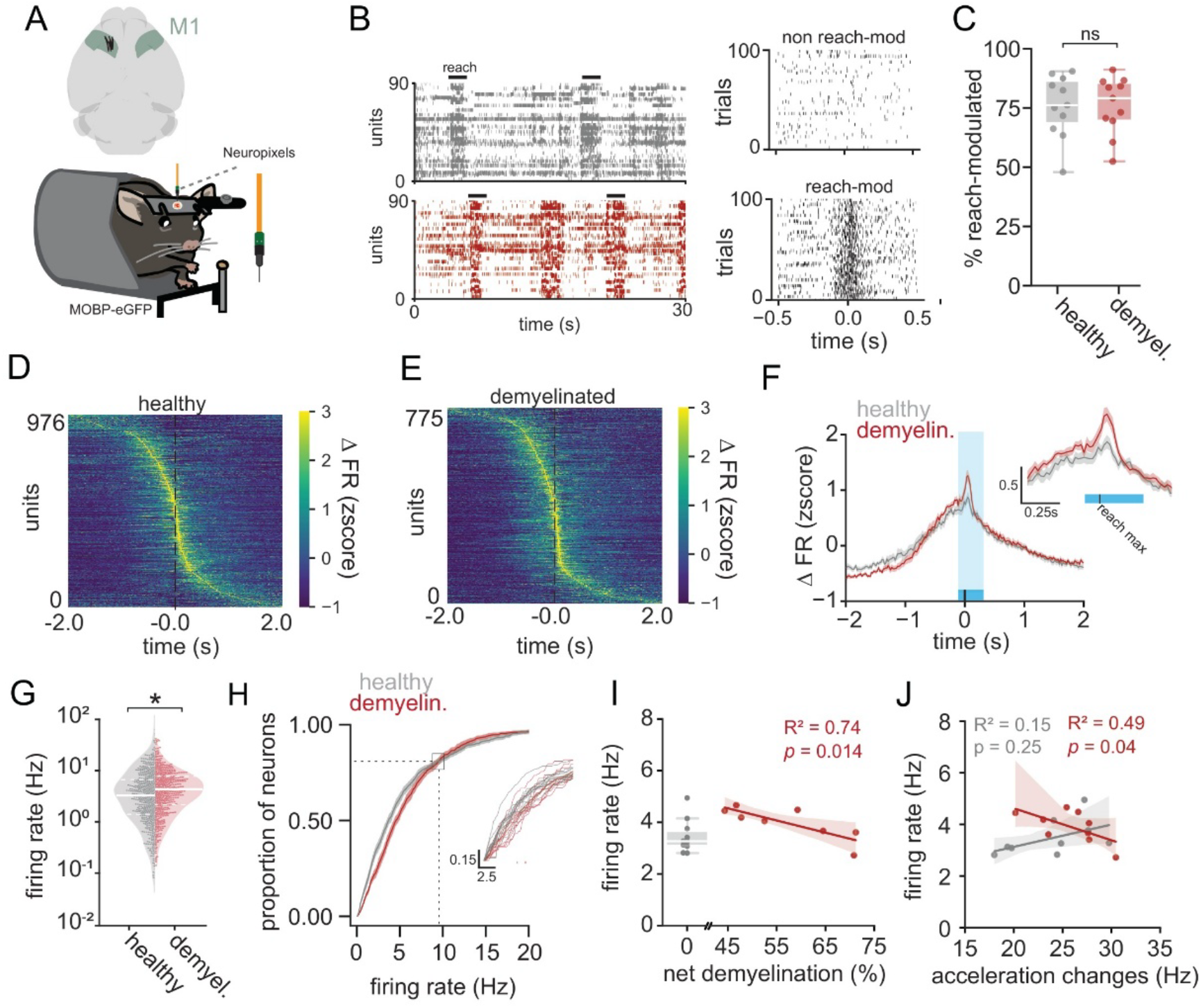
Motor cortical firing rates scale with demyelination severity. **(A)** Schematic of acute Neuropixels recordings from M1 in *Mobp*-eGFP mice performing the skilled reach task. Probe tracks (black) aligned to histology confirm placement in left M1. **(B)** Example rasters from a healthy (gray) and demyelinated (red) mouse during 30 seconds of reaching; reach periods indicated by black bar. **(C)** Example non–reach-modulated (top) and reach-modulated (bottom) units aligned to reach max. Right: the proportion of reach-modulated units did not differ between groups (n=11 per group, t=0.036, p=0.971; box plots represent median, interquartile range). **(D–E)** Population firing-rate heatmaps aligned to reach max for healthy (**D**) and demyelinated (**E**) mice, sorted by peak response time. **(F)** Average PSTH of z-scored firing for all reach-modulated units in healthy and demyelinated mice. Mean ± s.e.m.; reach window highlighted in blue (inset: magnified view). **(G)** Distribution of log-normalized firing rates for reach-modulated units. Demyelinated mice had elevated reach-modulated firing rates (REML, p=0.010; n=11 healthy, 976 units; n=11 demyelinated, 775 units; violin plots indicate median, interquartile range). **(H)** Cumulative firing-rate distribution showing a rightward shift in demyelinated mice across the 0–10 Hz range. Inset shows individual mice shaded by group (n=11 per group). **(I)** Mean reach-modulated firing rate per animal negatively correlated with cortical demyelination (n=11 healthy mice, n=8 demyelinated mice, R^2^=0.74, p=0.014, Pearson). **(J)** Mean reach-modulated firing rate also negatively correlated with reach acceleration changes in demyelinated mice (n=9, Pearson), but not healthy (n=10, Pearson) linking motor cortical activity to irregular limb kinematics. Boxplots represent median and interquartile range. Shaded regions denote 95% confidence intervals around linear regression fits. ns, not significant. *p < 0.05

To further investigate the relationship between firing rates and movement output, we isolated units specifically modulated during reach execution (**Figure 3B, see Methods**). The proportion of reach-modulated units was similar in healthy (75.91 ± 3.93%) and demyelinated (76.11 ± 3.63%) mice (**Figure 3C**), indicating preserved encoding capacity^4^. We aligned this spiking activity to reach max, a consistent kinematic landmark across trials, and examined z-scored firing within a four second window chosen to capture reach preparation, execution, and return to baseline. This approach revealed temporally organized neural population activity in both groups (**Figure 3D,E**), consistent with patterns underlying skilled movement execution^4^. Despite this preserved temporal structure following demyelination, peri-stimulus time histograms (PSTHs) revealed a global increase in movement-related firing (**Figure 3F, Figure S4F**), which is consistent with previous work showing heightened cortical excitability after myelin loss^8,16^. Quantifying firing rates across the reach window confirmed elevated movement-related activity in demyelinated mice (healthy: 3.3 ± 0.14 Hz, n = 11; demyelinated: 4.0 ± 0.20 Hz), apparent in both mouse-averaged firing rates (**Figure S4G**; p = 0.0082, Mann–Whitney U) and mixed-effects modeling accounting for inter-animal variability (**Figure 3G**; p = 0.010, REML). This enhancement was concentrated to neurons below 9.4 Hz, reflected by a right-shifted cumulative distribution that intersected with the healthy distribution just below 10 Hz (**Figure 3H**). Notably, animal-averaged reach-modulated firing rates correlated with demyelination severity, where moderate myelin loss increased firing rates, which scale down with more demyelination (**Figure 3I**; R^2^ = 0.74, p = 0.014). These data directly implicate myelin loss in altered reach-related M1 neuronal activity.

Importantly, myelin loss did not correlate with basic reach parameters (velocity, pathlength) or reach consistency, indicating that elevated firing did not simply reflect rudimentary kinematic differences (**Figure S4H-J**). In contrast, we found that firing rates were correlated with the observed myelin-dependent impairment in smooth movement (R^2^ = 0.49, p = 0.04), a relationship not present in healthy controls (R^2^ = 0.15, p = 0.25; **Figure 3J**). Together, these results indicate that increased severity of demyelination is associated with altered neuronal responsiveness and more irregular ongoing movement, linking myelin-dependent changes in M1 activity directly to impaired skilled motor output.

### Demyelination alters cell type–specific neuronal activity in M1 during movement

Deep layer corticospinal neurons of motor cortex drive motor output and their activity is dependent on fast timescale regulation by PV interneurons^19,27^. Given the growing evidence that inhibitory interneurons rely on the function of myelin^17,28^, we next examined whether myelin loss selectively affects these circuit dynamics driving motor output. Leveraging the dense laminar sampling of Neuropixels probes, we determined the cortical depth of each unit relative to a physiologically defined surface channel (**Figure S5A; Methods**). Units were categorized as superficial (putative layers 1–3) or deep (putative layers 4–6) based on established depth boundaries^29,30^ (**Figure 4A**), with comparable unit distributions across depth between groups (**Figure 4B**). Units were further classified as regular spiking (RS; putative excitatory) or fast spiking (FS; putative inhibitory) using conventional waveform criteria^31,32^ (**Figure 4C; Figure S5B**), enabling us to examine cell-type–specific activity across cortical depth during motor output.

**Figure 4.**
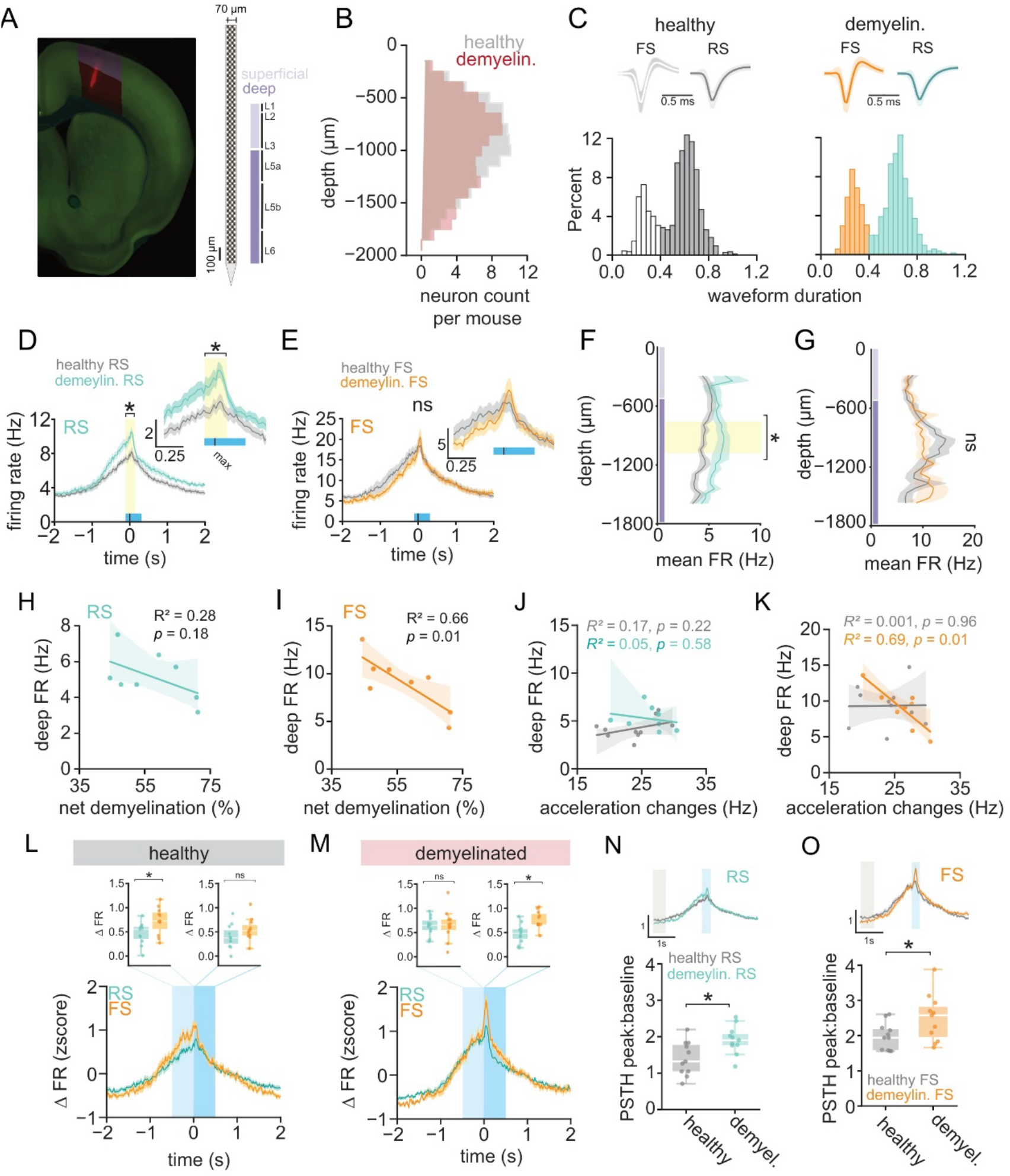
Demyelination alters cell type–specific neuronal activity in M1 during movement. **(A)** Coronal section showing Neuropixels probe placement in M1 of an MOBP-eGFP mouse, with schematic of probe depth marking superficial (L1–3) and deep (L5–6) layers. **(B)** Distribution of recorded unit depths did not differ between healthy (gray) and demyelinated (red) mice (U = 35, p = 0.10, Mann Whitney). **(C)** Classification of regular-spiking (RS) and fast-spiking (FS) units based on waveform duration, with representative waveforms and duration distributions. **(D)** RS neurons showed increased reach-evoked firing for 220 ms during reach (yellow window, n=11 healthy mice, n=11 demyelinated mice, cluster-based permutation test, p < 0.05). **(E)** FS neurons did not show significant demyelination-induced firing changes during the reach (n=11 healthy mice, n=11 demyelinated mice). **(F)** RS firing increased selectively in deep cortex following demyelination (p < 0.05). **(G)** FS firing did not differ across cortical depth. **(H)** Deep-layer RS firing was not related to demyelination severity (R^2^ = 0.28, p = 0.06, Pearson, n=8 demyelinated animals). **(I)** Deep-layer FS firing was negatively correlated with demyelination severity (R^2^ = 0.66, p = 0.02, Pearson, n=8 demyelinated mice). **(J)** Deep-layer RS firing rates were not correlated acceleration changes (n=10 healthy mice, n=8 demyelinated mice, Pearson). **(K)** Deep-layer FS firing was negatively correlated with acceleration changes in demyelinated mice but not healthy mice, linking reduced inhibition to irregular kinematics (n=10 healthy mice, n=8 demyelinated mice, Pearson). **(L–M)** Reach-aligned, z-scored firing rates for RS (teal) and FS (orange) neurons in healthy **(L)** and demyelinated **(M)** mice. Insets show average rates across the early reach window (light blue) and late reach window (darker blue), demonstrating stronger early FS recruitment in healthy mice and stronger late FS recruitment in demyelinated mice. Solid lines/shaded regions indicate mean±SEM, respectively. **(N)** Peak reach-evoked modulation (PSTH peak:baseline) of RS units increased following demyelination. Top: average RS PSTHs between healthy (gray, n=11 mice) and demyelinated (teal, n=11 mice) with baseline window indicated in gray; bottom: quantified peak modulation for healthy and demyelinated mice. **(O)** Peak reach-evoked modulation of FS units was also elevated after demyelination. Boxplots represent median and interquartile range. Shaded regions denote 95% confidence intervals around linear regression fits. ns, not significant. *p < 0.05

To determine how demyelination differentially affects putative excitatory and inhibitory populations, we used a cluster-based permutation test to identify phases where demyelinated firing exceeded healthy levels during reach. RS units showed a sustained increase beginning at reach initiation (**Figure 4D**), whereas FS units never exceeded healthy activity levels (**Figure 4E**). This increase localized to deep-layer RS units (760–1100 µm; **Figure 4F**), while FS units showed a slight, nonsignificant decrease at the same depth (**Figure 4G**). This suggests that the observed increase in reach-related firing rates is driven primarily by deep layer excitatory neurons. Within individual mice, the activity of both deep-layer RS and FS neurons decreases with demyelination severity, but only deep-layer FS activity showed a clear relationship with myelin loss (**Figure 4H,I**). Consistent with this selective vulnerability, deep-layer FS firing-but not RS firing-correlated with impaired smooth movement (**Figure 4J,K**). Thus, demyelination disproportionately affects deep cortex, increasing RS firing rates and linking FS output to deficits in motor execution.

To determine how demyelination might influence the timing of excitatory and inhibitory neuron response during reach, we compared z-scored activity across reach phases, allowing temporal modulation to be evaluated independent of baseline firing rate. In healthy mice, FS units showed earlier and stronger modulation than RS neurons in the initial reach epoch, 1 second before reach max (**Figure 4L**). Demyelination reversed this pattern, in which FS activity initially matched RS activity, but became more strongly engaged after reach max (**Figure 4M**). Both RS and FS populations also showed sharper reach-evoked peak responses in demyelinated animals than in healthy controls (**Figure 4N,O**), indicating more temporally concentrated rate modulation during movement execution. Together, these results reveal a demyelination-induced shift in the timing of FS relative to RS recruitment, accompanied by more temporally concentrated rate modulation during movement execution.

### Demyelination increases synchrony in primary motor cortex

Since fast-spiking PV interneurons play a central role in coordinating cortical activity^27^, their reduced recruitment with demyelination may have consequences on neuronal synchrony^17^. Task execution specifically engages PV interneurons to decorrelate neuronal activity^33^, so we next asked whether myelin loss increases neuronal synchrony in primary motor cortex. We assessed synchrony by measuring spike cross-correlograms (CCGs) between simultaneously recorded units (**Figure 5A**). Synchrony was identified by sharp, short-latency (±20 ms) peaks in jitter-corrected CCGs, which eliminate movement-driven covariation to isolate true fast spike-timing interactions^34^ (**Methods**). While inhibitory synchrony can be inferred from troughs in CCGs, too few events were observed to support meaningful conclusions (**Figure S6A**).

**Figure 5.**
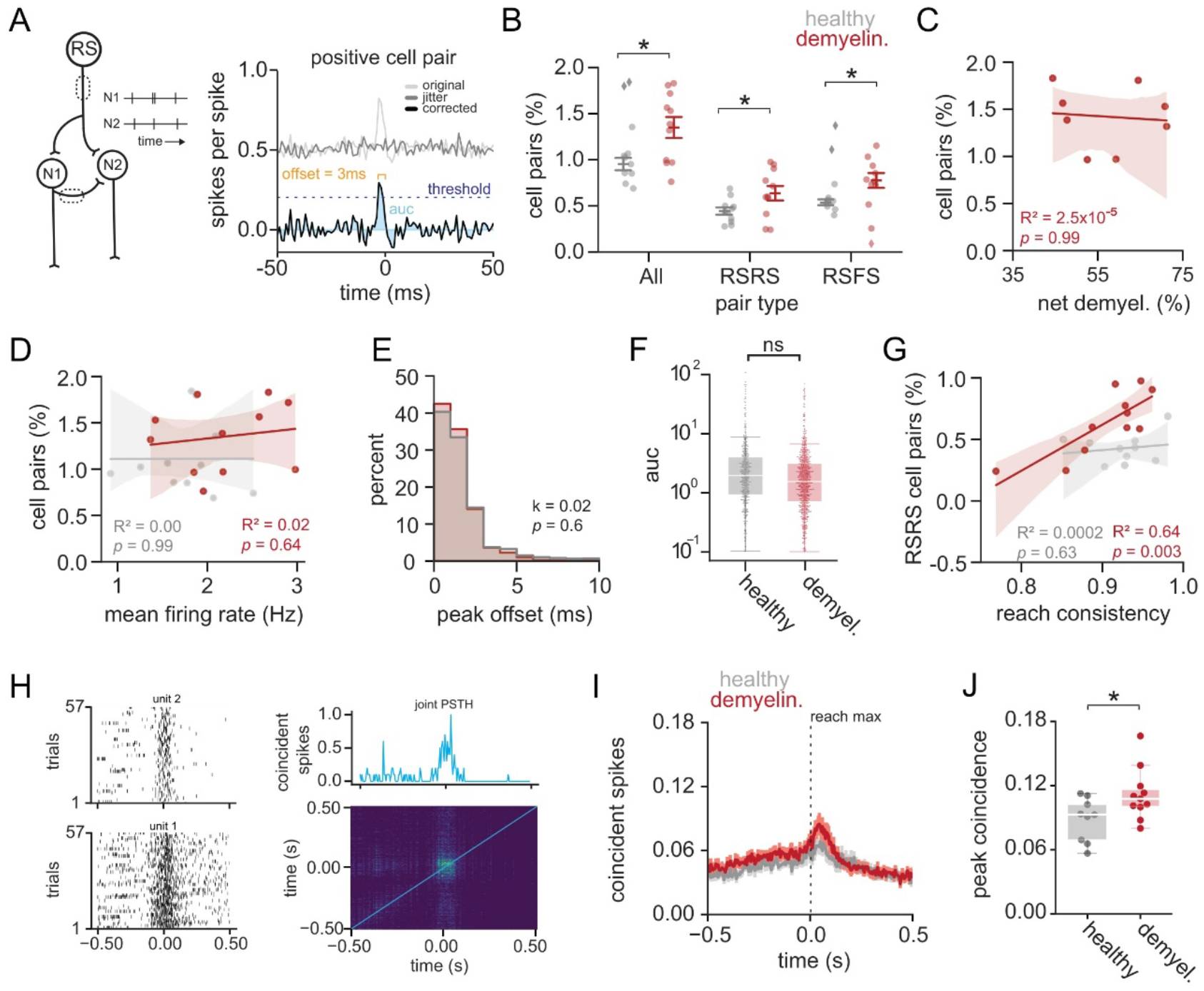
Demyelination increases synchrony in primary motor cortex. **(A)** Synchrony was assessed using jitter-corrected spike cross-correlograms (CCGs). Positive pairs were defined as short-latency peaks. **(B)** Demyelinated mice showed a higher proportion of positive excitatory cell pairs across all pair types (n=11 healthy, n=11 demyelinated; Bonferroni– Holm corrected p<0.05). **(C)** The proportion of positive pairs did not correlate with demyelination severity (R^2^ =0.0, p=0.991). **(D)** Firing rate did not influence proportion of pairwise synchrony in either group (Pearson). **(E)** Peak CCG offsets were unchanged by demyelination (KS test, p=0.93). **(F)** Jitter-corrected CCG peak area was also unchanged (REML, p=0.47; box plots represent median, interquartile range). **(G)** RS–RS synchrony predicted increased reach consistency in demyelinated (p=0.003, R^2^=0.64) but not healthy mice. **(H)** Movement-related correlated activity was quantified using joint PSTHs (jPSTHs). Left: example rasters; right: coincident spikes along the line of coincidence. **(I)** Mean coincident activity during reaching (mean ± SEM; n=9 healthy mice, n=11 demyelinated mice). **(J)** Demyelination increased peak coincident activity between movement-modulated pairs (n=9 healthy mice, n=11 demyelinated mice, Mann–Whitney U=21.0, p=0.018; box plots represent median, interquartile range). Boxplots represent median and interquartile range. Shaded regions denote 95% confidence intervals around linear regression fits. ns, not significant. *p < 0.05

To quantify overall synchrony, we calculated the proportion of synchronous unit pairs. Demyelinated mice exhibited a higher proportion of synchronous pairs overall (healthy: 0.95% ± 0.06%; demyelinated: 1.3% ± 0.11%) and across excitatory pair types (RS→RS, RS→FS; **Figure 5B**). This increase did not scale with demyelination severity (**Figure 5C**) and was not explained by elevated firing rates (**Figure 5D; Figure S6B,C**), indicating a generalized enhancement of fast-timescale synchrony. Measures of spike-timing precision-including CCG peak offset and peak area (auc, area under curve)- were unchanged (**Figure 5E,F**), and the spatial distances between paired units were similar (**Figure S6D**), suggesting preserved timing fidelity and layer-to-layer communication despite myelin loss. Notably, stronger RS→RS synchrony predicted better reach consistency in demyelinated mice, whereas no synchrony metric related to smooth movement in healthy animals (**Figure 5G; Figure S6E,F**), indicating that excitatory synchrony may stabilize reach trajectories.

We next asked whether increases in synchrony were specifically engaged during movement. Because cross-correlograms can be noisy when data are limited to reach epochs, we used cumulative joint PSTHs (jPSTHs) to quantify synchrony across reaches between all movement-modulated unit pairs (**Figure 5H**). Demyelinated mice showed higher peak coincident firing (0.11 ± 0.01 normalized spikes/bin) than healthy controls (0.08 ± 0.01; **Figure 5I,J**). This jPSTH coactivity did not vary with demyelination severity or firing rate (**Figure S6G-H**), indicating that demyelination induces increased fast-timescale synchrony in M1 that is enhanced specifically during movement.

### Action potential propagation failures, but not conduction delays, increase firing rates and correlated activity in a computational model of primary motor cortex demyelination

We next sought to define the mechanisms by which demyelination alters firing rates and neuronal synchrony in primary motor cortex. Although myelin is traditionally thought to accelerate action potential (AP) conduction, delays are expected to be minimal along short intracortical axons^35–37^. Recent work instead suggests that myelin supports reliable AP fidelity, particularly in fast-spiking interneurons, where demyelination produces propagation failures and disrupts inhibition^17,38^.

Given that extracellular recordings cannot easily detect AP propagation failures, we used computational modeling to examine whether our experimental effects could arise from either intracortical conduction delays or AP propagation failures across cell types. We developed a biologically constrained recurrent excitatory(E) - inhibitory(I) local circuit network fit to reproduce observed firing-rate distributions (**Methods**). We simulated demyelination by introducing cell-type–specific conduction delays or AP failure at levels consistent with putative myelin loss (**Figure 6A**), enabling us to identify the relative contribution these modes of failure may have to the experimental observations.

**Figure 6.**
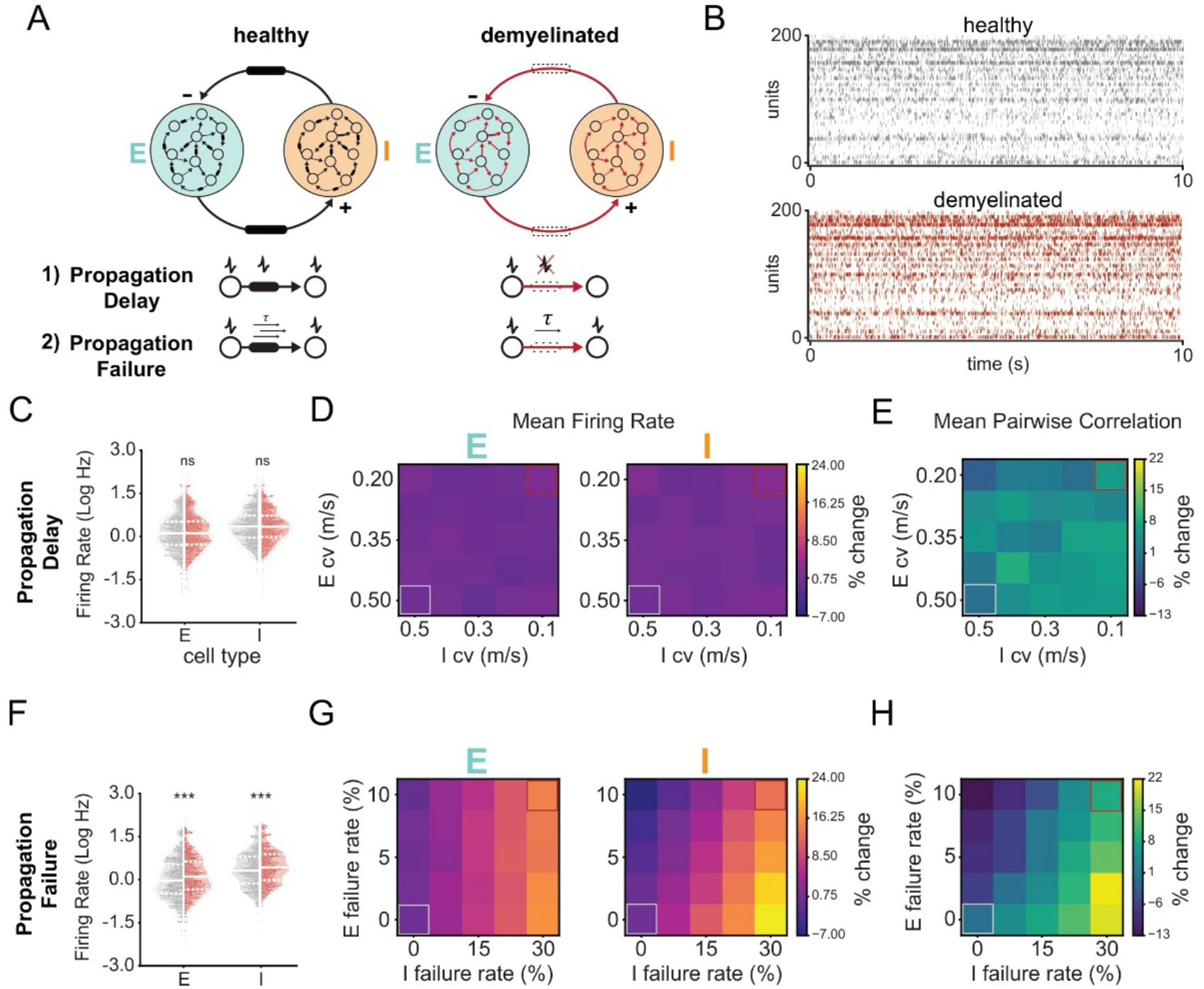
Action potential propagation failures, but not conduction delays, increase firing rates and correlated activity in a computational model of primary motor cortex demyelination. **(A)** Schematic of the E–I network, with intact myelination (black, healthy) or impaired myelination (red, demyelinated) implemented as either reduced conduction velocity or propagation failure. **(B)** Example raster plots for healthy and demyelinated networks under propagation failure (10 s window; neurons 0–99 = E, 100–200 = I). **(C)** Conduction velocity deficits did not alter firing rates for E or I cells. Each point shows mean firing rate of one neuron across ten 60-s trials; violins show population distributions. **(D)** Percent change in mean firing rate across conduction velocity levels for E and I populations; healthy and demyelinated conditions highlighted. **(E)** Percent change in mean pairwise spike-train correlation across conduction velocity levels, averaged across all neuron pairs. **(F–H)** Under propagation failure, both E and I populations showed increased firing (F) and increases in firing-rate and correlation heatmaps (G–H) relative to healthy networks (0% failure), demonstrating that propagation failure, not slowed conduction, produces network hyperexcitability and hypersynchrony. Violin plots represent median and interquartile range. ns, not significant. ***p < 0.0001

Example spike trains from the model produced physiologically relevant dynamics consistent with experimental observations (**Figure 6B**). Reducing conduction velocity across excitatory or inhibitory neurons had little effect on firing rates (**Figure 6C,D**) or pairwise spike-train correlations (**Figure 6E**), indicating that slowed intracortical propagation alone cannot account for the experimental observations. In contrast, introducing propagation failure produced robust increases in firing rates (**Figure 6F**, p < 0.001, KS test) and a monotonic rise in mean activity as failure probability increased (**Figure 6G**). Network correlations also increased (**Figure 6H**), recapitulating the enhanced synchrony observed *in vivo*. Importantly, these differences appear most sensitive to increasing inhibitory propagation failure (**Figure 6G,H**, x axis). Together, these simulations indicate that demyelination-like propagation failure weakens inhibitory control, driving disinhibition and network-wide increases in excitability and correlated activity.

### Partial remyelination recovers neural activity in motor cortex but leaves smooth movement impaired

In demyelinating diseases such as multiple sclerosis, myelin loss is typically followed by a period of new myelin generation.^12^ However, remyelination is often incomplete, contributing to persistent disability^12,39^. Although remyelination can restore neural activity and prevent neurodegeneration^8,40^, the degree of myelin repair needed to recover normal circuit dynamics and movement remains unclear. To assess the effect of remyelination on behavioral and circuit recovery, we conducted longitudinal two-photon imaging and Neuropixels recordings during reach in a separate cohort of *Mobp-EGFP* mice 24 days after cuprizone removal (**Figure 7A,B**). Although overall myelin repair remained incomplete (**Figures 7C,D, S7A**), we found that remyelination increased myelin content from day 3 to day 24 in individual mice (49.3 ± 3.6% → 57 ± 4.6%, p = 0.036; **Figure 7E**) and that across cohorts remyelinated mice had higher myelin levels than the demyelinated cohort (57 ± 4.6% vs. 42.0 ± 3.5%, p = 0.02; **Figure 7F**). Remyelinated mice maintained similar performance in the reach task to healthy mice (**Figure S7B**, healthy: 41.94 ± 4.68%, remyelinated: 40.15 ± 3.22%), similar to the demyelination timepoint. Reach kinematics of remyelinated mice appeared to be more similar to healthy skilled movements and reach-to-reach consistency recovered to healthy levels (**Figures 7G,H; S7C-G**). However, acceleration changes-our measure of smooth movement-remained elevated (**Figure 7I**), indicating a persistent deficit despite improved trajectory stability.

**Figure 7.**
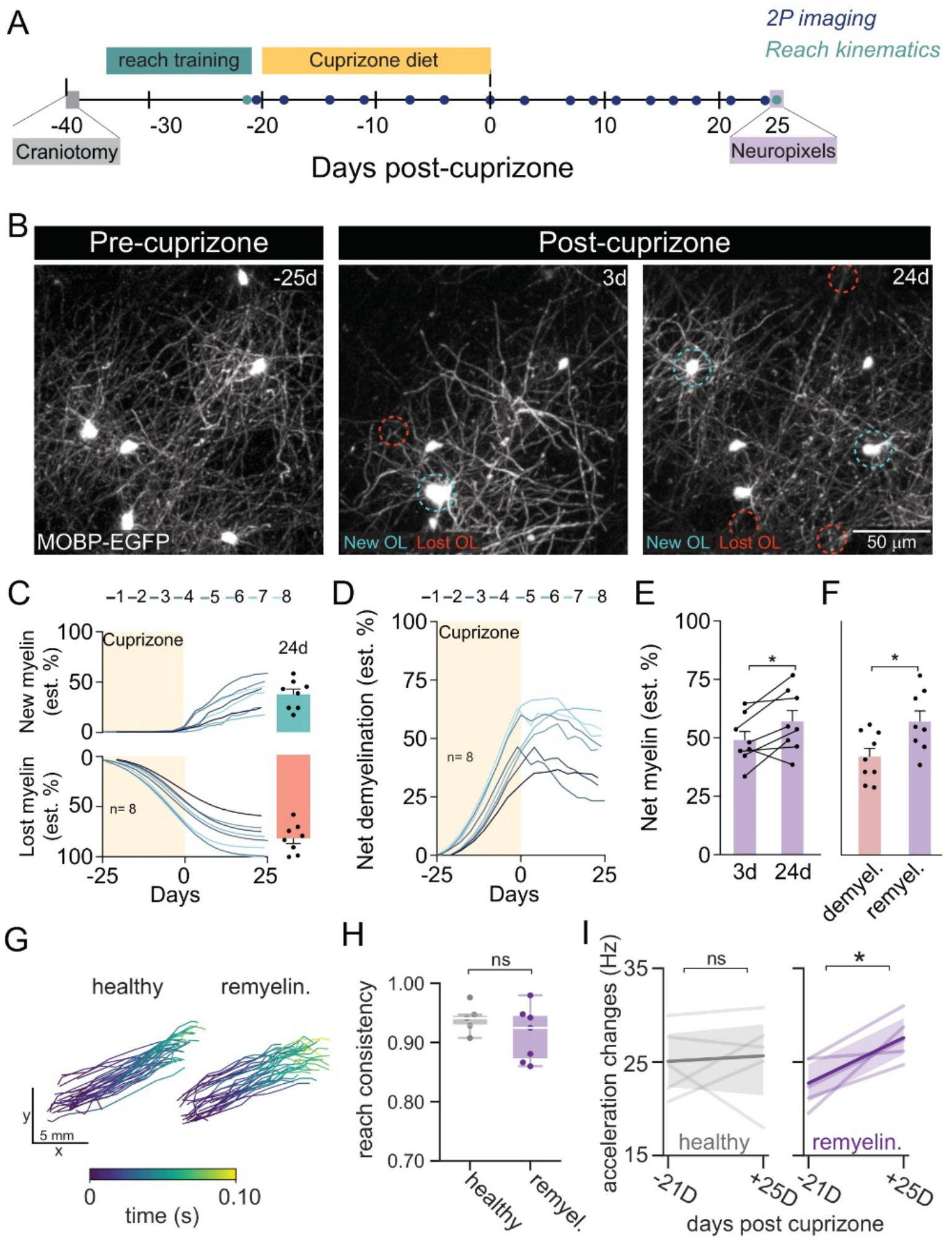
Reach consistency remains impaired with partial remyelination. **(A)** Experimental timeline. **(B)** Representative images from *in vivo* two-photon microscopy illustrating oligodendrocyte gain and loss. **(C)** Estimated dynamics of myelin loss and gain in individual mice. **(D)** Temporal dynamics of overall demyelination considering both myelin loss and gain. **(E)** Mice in the remyelinated cohort have more myelin at the 24D time point than at their 3D time point (t(7)=2.6, p=0.036, n = 8 remyelinated mice). **(F)** Mice in the remyelinated cohort (24d) have overall more myelin than mice in the demyelinated cohort (4d): t-test (t(15) =2.6, p = 0.02, n = 9 demyelinated and 8 remyelinated mice). **(G)** Representative reach trajectories from a healthy (left) and remyelinated (right) mouse performing the skilled reach task, color-coded by time from reach initiation to maximum. **(H)** Reach consistency, quantified as mean pairwise trajectory correlation, did not differ between healthy and remyelinated mice (t=1.0, p=0.33, t-test, n=6 mice per group; box plots represent median, interquartile range). **(I)** Rate of acceleration changes remained elevated following partial remyelination (n=5 healthy mice, t = −0.27, p = 0.80, n=5 remyelinated mice, paired t-test, t=-3.5, p=0.024, paired t-test). Boxplots represent median and interquartile range. Shaded regions denote 95% confidence intervals around linear regression fits. ns, not significant. *p < 0.05

Similarly, neural activity broadly recovered to healthy control levels. Both healthy and remyelinated mice exhibited preserved movement-related neural activation patterns during reach execution (**Figure S8A-D**), indicating stable neural encoding of the reach even weeks after training. Reach-modulated firing rates returned to healthy levels (**Figure 8A,B**) and no longer correlated with demyelination severity (**Figure 8C**). Synchronous pairs likewise normalized (**Figure 8D**) and no longer predicted reach consistency (**Figure 8E**), with greater myelin repair associated with reduced synchrony (**Figure 8F**). Cell type specific activity also recovered, where RS and FS units showed no firing-rate or modulation differences across the reach epoch (**Figure 8G; Figure S8E–F**) or across cortical depth (**Figure S8G**), and firing rates were not related to myelin loss (**Figure S8H**). Nonetheless, deep-layer FS firing rates remained correlated with reach acceleration changes (**Figure 8I**), whereas RS firing did not (**Figure 8H**), suggesting persistent sensitivity of motor behavior to residual inhibitory dysfunction. Together, these results show that partial remyelination restores major features of cortical network organization and reach consistency, but full recovery of movement may require more complete myelin repair or circuit adaptation.

**Figure 8.**
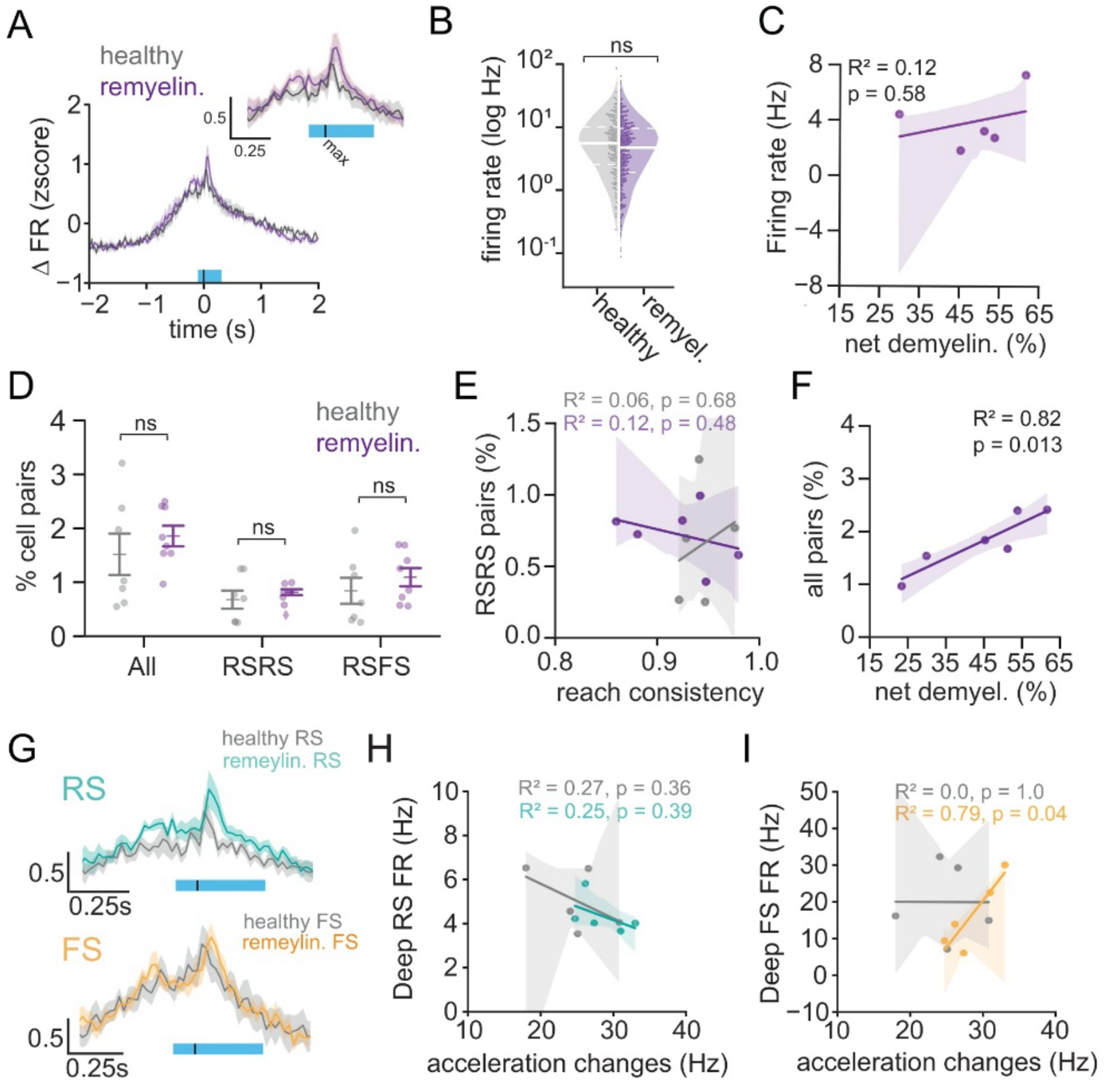
Partial remyelination restores motor cortical activity. **(A)** Average z-scored firing rates (ΔFR) of all reach-modulated neurons aligned to reach onset and maximum, showing comparable reach-evoked activity between healthy and remyelinated mice. **(B)** Mean firing rate did not differ significantly between groups (*p*=0.279, REML; violin plots indicate median, interquartile range). **(C)** Mean firing rate was not correlated with demyelination extent (R^2^ =0.12, p=0.58, Pearson, n=5 mice). **(D)** Proportion of cell pairs (all, RS–RS, RS–FS) did not differ significantly between healthy and remyelinated mice (n=7 mice each group) **(E)** The proportion of RS–RS cell pairs was not correlated with reach consistency in either group (Pearson, n=5 healthy mice, n=6 remyelinated mice). **(F)** The overall proportion of correlated cell pairs positively correlated with residual cortical demyelination (R^2^ = 0.82, p = 0.013, Pearson, n=6 mice). **(G)** Average z-scored neural activity PSTH for reach-modulated RS and FS neurons, aligned to reach maximum, showing preserved reach-evoked activity (n=7 mice each group). **(H)** Deep-layer RS neuron firing rates did not correlate with changes in acceleration, or smooth movement (R^2^ = 0.25, p = 0.036, Pearson, n=5 mice each group). **(I)** Deep-layer FS neuron firing rates positively correlated with acceleration changes (R^2^ =0.79, p = 0.04, Pearson, n=5 mice each group), indicating abnormal inhibitory activity with persistent movement irregularity. Boxplots represent median and interquartile range. Shaded regions denote 95% confidence intervals around linear regression fits. ns, not significant. *p < 0.05

## DISCUSSION

Here, we provide a behaviorally anchored, circuit-level framework for how myelin shapes motor function. By directly linking longitudinal measurements of myelin loss with *in vivo* population recordings and high-resolution kinematic analyses during skilled reaching, we demonstrate that cuprizone-induced demyelination impairs movement smoothness and reach consistency, alters movement-modulated excitatory and inhibitory activity, and increases fast-timescale synchrony in a manner that predicts motor output. Guided by experimental observations, we used a computational model to show that failures in inhibitory axonal spike propagation provide a key mechanism connecting myelin loss to circuit function. Although partial remyelination normalizes cortical network-level metrics and reach consistency, movement smoothness remained impaired, revealing a potential vulnerability in inhibitory networks. Together, these results integrate cellular models of demyelination with motor impairments observed in disease by demonstrating how myelin supports the cortical circuit dynamics underlying skilled behavior.

### Behavioral impact of myelin loss

Motor deficits in multiple sclerosis frequently manifest as inconsistent or poorly coordinated movements, including increased movement jerk and reduced smoothness across a broad range of upper-limb disability. In fact, subtle deficits in stable movement are detectable even in individuals without clinically diagnosed motor impairment^41–43^. Our findings parallel these clinical observations, where demyelination impaired both ongoing reach smoothness and across-trial reach consistency. Importantly, the three-week cuprizone treatment paradigm used here induces substantial myelin and oligodendrocyte loss without neuronal degeneration or synaptic changes, allowing us to isolate myelin-specific contributions to behavior^13^. We find a graded relationship between myelin loss and smooth movement, which suggests that the characteristic irregularity of movements in MS may arise, in part, from the loss of myelin itself rather than from autoimmunity or neurodegeneration. This finding highlights a critical role for myelin in supporting smooth and reliable motor execution.

Although cuprizone-induced demyelination does not cause uniform oligodendrocyte loss across brain regions^44^, recording from the motor cortex captures the integrated consequences of this widespread myelin loss. Since M1 serves as the major cortical output to spinal circuits and the convergence point for long-range motor pathways relayed through thalamus, M1 activity reflects the combined impact of myelin disruption across distributed motor networks on voluntary motor execution. However, studies utilizing multi-region recordings would be required to delineate the contributions of myelin loss across motor regions to movement output.

### Inhibitory circuit consequences of myelin loss

Motor output depends on tightly coordinated interactions between excitatory neurons and locally connected parvalbumin (PV) interneurons^1,27,45^. We found that reductions in FS firing rates, thought to reflect the high frequency firing of PV interneurons, during reach was driven by demyelination severity and robustly predicted impaired movement smoothness in demyelinated animals. This finding extends current views of cortical myelination by highlighting a role for myelin in supporting the physiological demands of fast-spiking PV interneurons during movement^17,46^. In line with this view, recent studies report robust evidence of AP propagation failures following cuprizone-mediated demyelination^16,17,35^. Incorporating these experimentally constrained levels of action potential propagation failure selectively in inhibitory neurons in a recurrent network model recapitulates the observed hyperexcitability and hypersynchrony in excitatory networks^17^.

The experimental observations in RS and FS activity were most pronounced in deep-layer motor cortex, where layer 5 corticospinal neurons rely on strong recurrent networks that operate in an inhibition-stabilized regime, requiring rapid, precise PV inhibition to maintain stable activity^47,48^. Under this regime, even modest disruptions to myelin-dependent inhibitory fidelity can produce increases in excitatory firing, providing insight into the altered RS:FS dynamics engaged during skilled reach in our data. Supporting this framework, genetic manipulation of PV interneurons, as well as myelin-targeted perturbations consistently increase kinematic variability and impair learning of skilled movement, respectively^1,8,9,49^. This framework supports a circuit-driven role for myelin, where early demyelination disproportionately disrupts inhibitory function during behavior, shifting cortical activity toward enhanced excitatory drive. However, with more extensive myelin loss, elevated RS and FS reach modulated activity globally scaled down. Since extracellular recordings cannot detect axonal propagation failures, such failures are not reflected in firing rates measures. Instead, early disinhibition due to heightened vulnerability of PV interneurons likely transitions to broader conduction and metabolic failure also affecting excitatory neurons and their long-range inputs. Severe demyelination is known to increase propagation block^38^, reduce feedforward thalamocortical drive^49^, and impose substantial energetic burden on axons^50^, ultimately limiting the ability of all cell types to sustain excessive spiking^17,38^. This pattern aligns with ex vivo findings that prolonged cuprizone exposure diminishes EPSPs and ectopic AP generation, and increases conduction failure^16,17,51^. Our study contributes to emerging views that demyelination does not affect all cortical circuits equally, and that the behavioral consequences of myelin loss depend on which axons are affected (e.g., PV interneurons versus RS pyramidal neurons; L5 corticospinal versus intracortical projections). Further studies of circuit-specific demyelination-resolving how myelin loss differentially impacts distinct cortical cell types and projections-will be crucial for understanding how progressive demyelination drives worsening motor impairment.

### Compensatory synchrony in the demyelinated cortex

Reproducible neural activity patterns in motor cortex underly skilled movement^4,5^. The increased fast timescale synchrony between M1 units in our data like likely reflects this pattern of neural activity known to be tightly coupled to consistent movement execution^4^. Despite clear myelin-dependent changes in firing rate, hypersynchrony persisted across all levels of demyelination. This indicates that elevated fast-timescale coordination can arise from deficits in local inhibitory fidelity- as supported by our computational model-but is not directly proportional to the severity of myelin loss. Instead, enhanced excitatory synchrony predicted better reach consistency, suggesting that coordinated activity may serve as a compensatory mechanism that stabilizes motor output when myelin is compromised. In this view, enhanced synchrony is recruited to produce consistent motor output in an otherwise impaired circuit. This interpretation aligns with work showing that synchronous thalamocortical input can rescue motor learning under demyelinating conditions, underscoring the importance of precise temporal coordination for effective motor output^49^.

In contrast to suppressing adult oligodendrogenesis during motor learning-which disrupts the acquisition of new motor commands^8^-our study focuses on the execution of a well-learned skill. Here, increased synchrony may reflect the cortex leveraging established recurrent architectures to counteract variability introduced by myelin loss. The absence of synapse loss and axonal degeneration in our three-week cuprizone treatment likely preserves the structural pathways required for this compensatory drive. Future work is needed to determine whether severe or chronic demyelination precludes this circuit adaptation by inducing structural degradation and neuronal atrophy^13,52^. Defining when and how this synchrony fails could be key to understanding the limits of functional recovery in demyelinating disease.

### Remyelination and recovery

Remyelination partially restores conduction and neuronal excitability but is often incomplete and heterogeneous across cortical circuits^13^. In our dataset, partial remyelination normalized firing rates, functional connectivity, and reach-to-reach consistency, yet smooth ongoing movement remained impaired. This pattern is consistent with clinical observations in multiple sclerosis, where movement abnormalities often persist despite remyelination or disease-modifying therapies^41–43,53^. A likely explanation is that incomplete remyelination of inhibitory interneurons limits the recovery of cortical inhibition which supports smooth movement. Although remyelination appears broadly distributed across neuronal subtypes, inhibitory circuits may require more complete or spatially patterned myelin restoration to reestablish the metabolic and biophysical demands of fast-spiking propagation^46^. The persistence of impaired smooth movement alongside elevated FS firing suggests that inhibitory network recovery lags behind broader circuit stabilization, although this observation requires further investigation. This discrete dissociation in circuit elements implies that consistent reach execution is reflected primarily in restored excitatory intracortical synchrony, whereas smooth ongoing movement requires precise inhibitory dynamics that may not fully recover with partial remyelination. Our data likely captures an early phase of remyelination; whether later stages could restore full inhibitory fidelity and smooth movement remains an open question.

Finally, our findings suggest that circuit-level signatures-such as firing modulation, synchrony profiles, and inhibitory fidelity-could serve as functional biomarkers of remyelination efficacy. Combining remyelination strategies with approaches that actively engage motor cortical circuits, including activity-dependent rehabilitation or neuromodulation, may further enhance recovery^8,54^. Successful restoration of motor functions will likely require both structural myelin repair and targeted modulation of cortical circuits to recalibrate the dynamics that enable precise motor output. These results establish a mechanistic foundation for understanding the role of myelin in regulating cortical circuits and provide a roadmap for therapies that jointly promote structural repair and circuit recalibration following demyelination.

## Supporting information

TableS1_KeyModelParameters

TableS2_StatsTable

## Resource Availability

### Lead contact

Information and requests should be directed to and will be fulfilled by the lead contacts, Daniel Denman Ethan Hughes Cristin Welle.

### Materials availability

This study did not generate new, unique reagents.

### Data and code availability

- Code is available at DOI: 10.6084/m9.figshare.30924077
- Any original data reported in this paper or additional information required to reanalyze the data are available from the lead contact upon request.

## Acknowledgements

We appreciate members of the Welle, Hughes, and Denman labs for comments and discussions. We would like to thank Rongchen Huang for cranial window implants, Anthony Chavez for brain sectioning, and Yessenia Mancha for staining and data acquisition. work was funded by the following sources: National Institutes of Health grants NS137560 (C.G.W.), NS115975 (C.G.W., E.G.H.), NS125230, NS132859, NS134829 (E.G.H.) and NS120850, EY028612 (D.J.D.). National Multiple Sclerosis Society Postdoctoral Fellowships FG-2107-38324 (G.D.F.N.). National Science Foundation Graduation Research Fellowship Program DGE-1938058 (K.G.G).

## Author Contributions

C.G.W, E.G.H., D.L.N., conceived the project. K.G.G. and G.D.F.N. wrote the original manuscript, administered the project execution, and jointly performed experiments (surgeries, drug treatment, *in vivo* imaging, in vivo electrophysiology). G.D.F.N. carried out *in vivo* imaging and analysis, myelin model, and all statistical analysis for oligodendrocytes and myelin data, and created Figure 1, Supplementary Figures 1-2, and contributed to Figure 7. K.G.G performed electrophysiology experiments, data analysis, and statistical analysis for Figures 2-5; 7-8, as well as Supplementary Figures 3-8. Kinematic data acquisition was established by W.R.W and kinematic data analysis was performed by E.C. and K.G.G. J.L. and T.N developed the computational model, created Figure 6, and wrote the relevant results section. D.N. performed pilot experiments and analysis. G.D.F.N. performed immunohistochemistry, confocal imaging, and analysis that are reported in Supplementary Figs. 1&2. K.G.G and G.D.F.N sectioned the brain tissue. A.L. sectioned, performed immunohistochemistry and analysis of synaptic colocalization. The project was supervised by D.J.D, E.G.H, and C.G.W. The manuscript was edited and revised by J.L., T.N., G.D.F.N., K.G.G., D.J.D., E.G.H., C.G.W.

## Supplemental Information

Supplementary Figures S1-8

Table S1. Key model parameters related to Figure 6

Table S2. Detailed statistical information related to Figures 1-8, S1-8

## Declaration of generative AI and AI-assisted technologies in the writing process

During the preparation of this work the authors used ChatGPT in order to check grammar and improve clarity. No generative AI tools have been used to produce any new content. After using this tool, the authors reviewed and edited the content as needed and take full responsibility for the content of the published article.

## STAR★METHODS - KEY RESOURCES TABLE

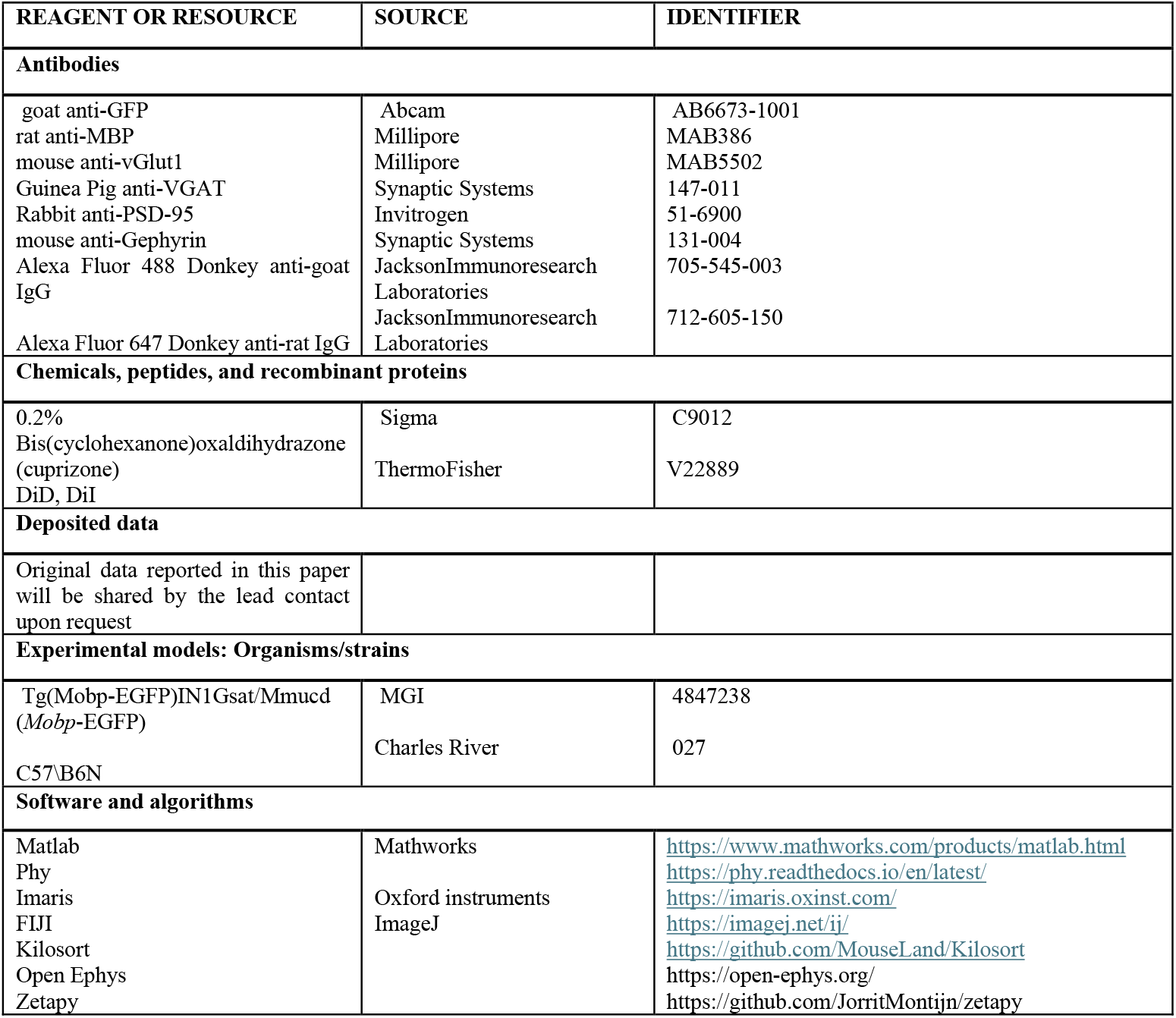

### Experimental Model and Study Participant Details Animals

All animal experiments were conducted in accordance with protocols approved by the Animal Care and Use Committee at the University of Colorado Anschutz Medical Campus. Male and female mice used in these experiments were kept on a 14-h light, 10-h dark schedule with ad lib access to food and water, aside from training-related food restriction (see “forelimb reach training” section). All mice were randomly assigned to conditions and were age-matched across experimental groups. Congenic C57BL/6N MOBP-EGFP (Tg(Mobp-EGFP)IN1Gsat/Mmucd, MGI:4847238) mice, which have been previously described^55,56^, were used for two-photon imaging+electrophysiology experiments. Wild-type C57\B6N Charles River wild-type mice were used in electrophysiology-only experiments. Genotyping for the *Mobp-EGFP* transgene was carried out by Transnetyx Inc. using the primers F: 5’ CGTCGTCCTTGAAGAAGATGGT 3’ and R: 5’ CACATGAAGCAGCACGACTT 3’ and Probe: 5’ CATGCCCGAAGGCTAC 3’.

Multimodal data was collected for each group of replicated experiments. Pilot experiments were conducted in wild-type mice in which no imaging was performed, but endpoint electrophysiology and kinematics were collected (n=3 4D-post cup healthy mice, n=3 4D post-cup demyelinated mice, n=2 24D post-cup healthy mice, n=2 24D post-cup demyelinated mice). Three proceeding groups of replicated experiments included multimodal data collection, pre-cuprizone kinematics (n=6 4D post-cuprizone healthy mice and n=6 4D post-cuprizone demyelinated mice, and n=5 24D post-cuprizone healthy mice and n=5 24D post-cuprizone demyelinated mice), post-cuprizone kinematics (n=10 healthy 4D post-cuprizone mice, n=10 4D post-cuprizone demyelinated mice), two photon imaging of oligodendrocytes (n=8 4D demyelinated mice, n=8 24D demyelinated mice). Electrophysiology recordings n=11 4D post-cuprizone healthy mice, n=11 4D post-cuprizone demyelinated mice, n=7 24D post-cuprizone healthy mice, n=7 24D post-cuprizone demyelinated mice). Not all modalities could be collected for each replicated experiment due to technical limitations.

## Method Details

### Cranial window implantation surgery

6–8-week-old mice were anesthetized with isoflurane inhalation (induction, 5%; maintenance, 1.5–2.0%, mixed with 0.5 L/ min O2) and kept at 37 °C body temperature with a thermostat-controlled heating plate. After removal of the skin over the left cerebral hemisphere, the skull was cleaned and a 2 × 2 mm region of skull centered over the primary motor cortex (0-2.0 mm anterior to bregma and 0.5–2.5 mm lateral to bregma) was removed using a high-speed dental drill. A piece of cover glass (VWR, No. 1) was then placed in the craniotomy and sealed with Vetbond (3 M) and then dental cement (C&B Metabond). A 5 mg/kg dose of carprofen was subcutaneously administered before awakening. For head stabilization during *in vivo* imaging, behavior and Neuropixels recording, a custom metal plate with a central hole was attached to the skull with dental cement.

### Cuprizone-mediated demyelination

Demyelination was induced in 9–10-week-old mice using 0.2% Bis(cyclohexanone)oxaldihydrazone (cuprizone; Sigma C9012), stored in a glass desiccator at 4 °C. Cuprizone was added to powdered chow (Envigo T.2918 M.15), mixed for ~10 min to ensure homogeneity, and provided to mice in custom feeders (designed to minimize exposure to moisture) for 21–25 days on an ad libitum basis. Feeders were refilled every 2–3 days, and fresh cuprizone chow was prepared weekly. Healthy mice received normal powdered chow in identical feeders. Cages were changed weekly to avoid build-up of cuprizone chow in bedding, and to minimize reuptake of cuprizone chow following cessation of diet via coprophagia.

### Two-photon microscopy

*In vivo* imaging sessions began three weeks post-surgery and took place 1–3 times per week (Figures 1B and 7A). During imaging sessions, mice were anesthetized with isoflurane and immobilized by attaching the head plate to a custom stage. Images were collected using a Zeiss LSM 7MP microscope equipped with a BiG GaAsP detector using a mode-locked Ti:sapphire laser (Coherent Ultra) tuned to 920 nm. The average power at the sample during imaging was 5–30 mW. Vascular and cellular landmarks were used to identify the same cortical region over longitudinal imaging sessions. *Mobp-EGFP* image stacks were acquired using a Zeiss W plan-apochromat ×20/1.0 NA water immersion objective giving a volume of 425 μm × 425 μm × 336 μm (1024 × 1024 pixels; corresponding to layers I-III, 0–336 μm from the meninges) from the cortical surface.

### Image processing and analysis

During data analysis, experimenters were blinded to the experimental groups. For presentation in figures, image brightness and contrast levels were adjusted for clarity. Images were analyzed using ImageJ (v1.54 f) ^57^ unless otherwise specified. For analyses of Mobp-EGFP cell density and MBP coverage from immunohistochemistry, images were quantified in the entire region of interest using 3D hysteresis thresholding from the 3D Suite plugin (ImageJ) (Ollion, J. et al., Bioinformatics 2013) for segmentation.

To evaluate the effects of demyelination on synaptic density, the SynBot^58^ ImageJ macro was used to quantify synapses by the colocalization between pre- and postsynaptic markers of excitatory (VGlut-1 and PSD-95) or inhibitory (VGAT and Gephyrin) synapses within the primary motor cortex. Four fields of view of size 54.15 by 34.67 µm were analyzed per animal, consisting of two randomly selected ROIs within the primary motor cortex per tissue section across two sections per animal.

Images of 3D volumes acquired through time using *in vivo* two-photon microscopy, were concatenated into a 4D hyperstack and then registered in ImageJ v1.46r, by sequentially using the Poorman3Dreg plugin and the Correct3D drift plugin (EGFP channel, rigid body registration)^59^.

Oligodendrocyte cell tracking was performed manually. A custom ImageJ script^55^ enabled recording of oligodendrocyte state (new, lost, or stable EGFP+ soma) across timepoints. To account for variability in the number of oligodendrocytes in the imaged area, all oligodendrocyte metrics are reported as a percentage of baseline (pre-cuprizone) values:

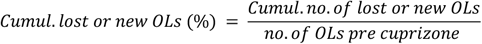

Myelin sheath tracing for Figure 1F was performed using Simple Neurite Tracer (ImageJ) at three different timepoints for each mouse: before cuprizone, around peak demyelination and during remyelination. The volume analyzed was 149.45 *μ*m x 149.45 *μ*m x 31.5 *μ*m (length x width x depth). The three animals examined showed a range of demyelination severity and of myelin repair. The number of sheaths in the timepoints post-cuprizone was normalized to the number pre-cuprizone in each animal to calculate the myelin reconstruction (%).

### Estimation of myelin sheath numbers

For estimation of myelin content, sheaths of individual oligodendrocytes were first traced and counted using ImageJ Fiji. For this, we used a dataset of longitudinal *in vivo* two-photon imaging of primary motor cortex including three animals from a different cohort. For the analysis of lost myelin sheaths, two animals included in this study were added to this dataset. For all the analyses, at least 9 oligodendrocytes from 3 mice were evaluated. Due to characteristics of the *MOBP-EGFP* reporter, new oligodendrocytes and their myelin sheaths are brighter at the time of their formation. Therefore, we counted all bright myelin sheaths that appeared together with the new oligodendrocyte (oligodendrocyte and myelin appear simultaneously with this reporter). To more easily distinguish myelin sheaths associated with the new oligodendrocyte, we selected new oligodendrocytes not neighbored by other cells that were new at the same or preceding timepoint. Considering our data and that from other publications, we estimate that each new oligodendrocyte makes 42.8775 sheaths (see Figure S1B), therefore:

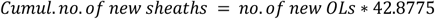

For the analysis of lost oligodendrocytes, we counted all sheaths that could be directly connected to the cell of origin (average±SEM: 23.42±1.77 sheaths per OL). The cumulative percentage of lost sheaths in relation to time of OL loss follows a pattern that can be modeled using the three parameter (3P) logistic regression formula (Figure S1E).

Therefore, the cumulative number of sheaths lost at a given day post cuprizone (dpc) can be estimated by summing the myelin sheaths lost by each individual oligodendrocyte as:

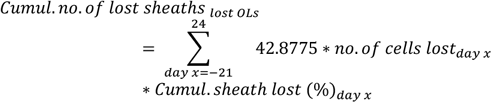

and:

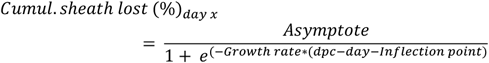

where:

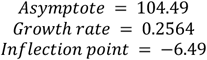

Since oligodendrocyte loss has not been finalized in demyelinated mice 3 dpc, it is important to account for future predicted OL loss when estimating current myelin loss. For that purpose, we took advantage of the direct linear relationship between OL loss at 3dpc and 24dpc (Figure S1F) to estimate the final cumulative OL loss (%):

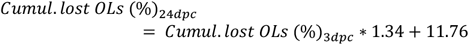

We then used this value as the asymptote on a 3P logistic regression to estimate the dynamics of OL loss (%), using the average values of growth rate and inflection point for our remyelinated (24dpc group, see Figure S1G) (0.2186 and 0.8562, respectively):

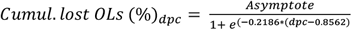

This allowed us to calculate the number and timing of expected OL loss between 3dpc and 24dpc in our demyelinated group. For the analysis of surviving OLs, we also only included sheaths that could be directly connected to the cell of origin (average±SEM: 16.2±1.6 sheaths per OL). On average, these cells lost 18.47±3.09% of their sheaths (Figure S1I). The temporal dynamics of sheath loss followed a Probit 2P function (Figure S1K), and, therefore, could be estimated as:

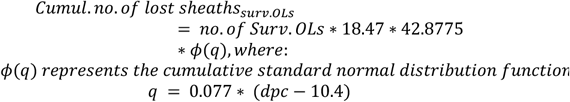

The myelin estimates for all oligodendrocytes were then integrated to calculate the number of myelin sheaths at a given timepoint:

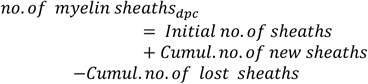

where:

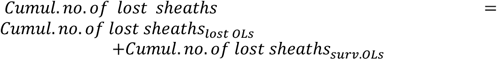

and:

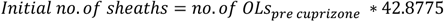

The percentage of lost/new sheaths was calculated as:

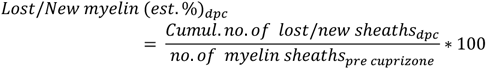

Importantly, the mathematical formula often estimates some small myelin sheath loss pre cuprizone. So, the *Cumul. no. of lost sheaths*_*pre cuprizone*_ was set to 0 by subtracting this value from all timepoints.

Net demyelination was calculated in an analogous way:

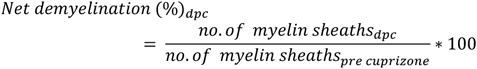

### Immunohistochemistry

Mice were anesthetized with an intraperitoneal injection of sodium pentobarbital (100 mg/kg body weight) and transcardially perfused with 4% paraformaldehyde in Phosphate-buffered saline (pH 7.4). Brains were postfixed in 4% paraformaldehyde overnight at 4 °C, transferred to 30% sucrose solution in PBS (pH 7.4) and stored at 4 °C until processing. Tissue was positioned in cryomolds with Tissue-Tek OCT, frozen, and sections were obtained using a cryostat (40 μm thick). Brain sections were immunostained using a free-floating method. Sections were blocked (5% normal donkey serum, 0.3% Triton X-100 in PBS, pH 7.4) for 1 h at room temperature, then incubated overnight at 4 °C with primary antibody followed by washes and secondary antibody incubation for 1 h at room temperature. Primary and secondary antibodies were diluted in the blocking buffer. Primary antibodies and dilutions were goat anti-GFP 1:500 (Abcam AB6673-1001), rat anti-MBP 1:500 (Millipore MAB386), mouse anti-vGlut1 1:200 (Millipore MAB5502), Guinea Pig anti-VGAT 1:500 (Synaptic Systems 147-011), Rabbit anti-PSD-95 1:1000 (Invitrogen 51-6900) and mouse anti-Gephyrin 1:500 (Synaptic Systems 131-004). Secondary antibodies (diluted 1:500) were Alexa Fluor 488 Donkey anti-goat IgG (Jackson Immunoresearch Laboratories 705-545-003) and Alexa Fluor 647 Donkey anti-rat IgG (Jackson Immunoresearch Laboratories 712-605-150). Sections were mounted on slides with Vectashield antifade reagent (Vector Laboratories). For quantifications of oligodendrocytes and myelin, tile scans of the entire region of interest were acquired using a laser-scanning confocal microscope (Leica SP8). For synapse quantification, images were acquired using a 3i MARIANAS inverted spinning disk confocal microscope at 63X oil immersion magnification using a z-stack with optimal step size (0.27 µm).

### *In Vivo* Electrophysiology

Acute *in vivo* Neuropixel recordings were conducted during 60-minute forelimb reach sessions at 4 days and 24 days following cuprizone-mediated demyelination. To record from neurons across depth in primary motor cortex, a Neuropixels 1.0 probe was inserted into caudal forelimb area of primary motor cortex (0.6 mm anterior and 1.5 lateral to bregma). Mice were first anesthetized (4% and then 1% for maintenance) and the imaging window was removed. Saline was used to temporarily lubricate the eyes and temperature was monitored and maintained using a closed-loop system. After a partial durectomy, the animal was head-fixed on the dexterous reaching behavior rig and DyeD or DyeI (ThermoFisher: V22889) was applied to the probe shank and inserted through the durectomy using a micromanipulator (Sensapex) to span approximately 2mm of primary motor cortex. Recordings were grounded to the animal and the exposed skull was protected with saline throughout the recording to maintain brain health. For all recordings, several hundred channels were left outside of the brain but left in saline to facilitate probe localization by determining a surface channel at the interface between brain surface and saline. Coronal sections containing the DiD- or DiI-labeled probe track were registered to the Allen Mouse Brain Atlas using the HERBS image-registration platform (https://github.com/Whitlock-Group/HERBS), enabling reconstruction of probe trajectories in three-dimensional atlas space and confirmation of recording locations within primary motor cortex (Figure 3A).

### Electrophysiology data acquisition and preprocessing

Neuropixels data was acquired at 30 kHz (spike band) and 2.5 kHz (LFP band) using OpenEphys^60^. Units were extracted with Kilosort^61^ 2.5 and manually refined using phy (github.com/cortexlab/phy). Putative excitatory (regular-spiking) and putative inhibitory (fast-spiking) units display well-documented, fundamental distinctions in waveform characteristics that reflect the underlying electrophysiological properties, such as waveform duration, peak-to-trough time, and end-slope. These two neuron subclasses were subdivided based on a widely-used waveform duration threshold of 0.4 ms confirmed in the experimental data by the time that falls between the two peaks in the bimodal distribution of waveform durations (Fig 4B).

### Cortical depth and layer labels

Depth estimation for units from each mouse was adapted from the ecephys code package https://github.com/AllenInstitute/ecephys_spike_sorting, in which the local field potential (LFP) data was used to identify the surface channel. Typically, up to 2 mm of the 7.5 mm shank were inserted into the brain. A depth profile was created for each mouse in which ten one-minute segments of the recording were filtered for delta(1-4 Hz), theta(4-8 Hz), alpha(8-13Hz), beta(13,30Hz), and gamma(30-90Hz) power and averaged for each channel. These low-frequency LFP bands indicate physiological brain activity and a sharp increase in power in these bands indicates a given channel is in the brain vs. outside the brain. The sharp increase was defined algorithmically as the greatest difference in power between channels, indicating the channel at which the power in these physiological ranges is lost between two neighboring channels demarcating inside and outside of the brain. The surface channel on the Neuropixels electrode array was registered to 0 µm and all channels below the surface channel were assigned a given depth based on the Neuropixels probe geometry. Depth for each unit was then assigned based on the highest amplitude channel and depth normalized to the surface channel. Superficial layer and deep layer assignments for each unit were based on an estimated <500 µm from brain surface (superficial) and >500 µm from brain surface (deep) based on L2/3 and L5 boundaries.

### Head-fixed Forelimb Reach Behavior

Mice were weighed and habituated to head fixation on the electrophysiology rig prior to forelimb reach training. The habituation days consisted of one 10-minute session followed by one 20-minute session on the following days. Once animals were voluntarily eating pellets while head-fixed, mice were weighed and food-deprived and forelimb reach training sessions began daily for 20 minutes. Mice learned to reach for a pellet using their right paw. All mice were positioned such that the pellet was 1 cm anterior and 1 mm lateral from the fixed paw rest bar. Successes were counted when the mouse reached out and successfully grasped the pellet and brought it back to their mouth. Throughout training, mice were kept on a restricted diet in which weight was monitored and maintained at 85-90% of their baseline (habituation) body weight. Mice were trained to the head-fixed reaching task until they achieved expert status, in which their success rate plateaued at >30% for two consecutive days. Mice were excluded from the study if they did not reach a plateau of 30% for 2 days after 14 days of training or did not learn to reach for the pellet after 10 days of training. Once mice reached expert levels, they were removed from food deprivation and placed either on the control diet or the cuprizone diet.

### Retraining

To ensure that mice would reliably reach during the recording session, a 20-minute retraining session was incorporated one day prior to the recording session. Mice were weighed and food deprived at either 2 days post-cuprizone (demyelination timepoint) or 23 days post-cuprizone (remyelination timepoint) and on the following day underwent a 20-minute session of the dexterous reaching task. If mice had less than 10 reaches, another day of food deprivation and retraining was completed to ensure proper motivation during the recording session. Mice were excluded from behavior and kinematic analysis if a success rate of >20% was not achieved at the Neuropixels recording timepoint to ensure comparisons represent maintained skilled motor output (n=1 healthy control animal, n=1 demyelinated animal were excluded).

### Kinematic Tracking and Analysis

Mouse reaching behavior was captured using two high speed (150 Hz) FLIR Blackfly S cameras (model BFS-U3-16S2M-CS, Edmund Optics) positioned in front of and lateral to the pellet. A custom DeepLabCut (DLC)^24^ network was trained to identify the paw and food pellet using manually annotated frames as previously described^23,62^. Custom MATLAB scripts extracted key reaching timepoints: reach initiation, maximum outward extension, and reach end. Manual refinement of these key timepoints, as well as outcome assignment (success or failure), were completed using custom python scripts. Three-dimensional paw position relative to the pellet and corresponding tracking confidence was extracted from the DLC network output from reach initiation to reach end using custom MATLAB scripts. Individual reaches were excluded if more than 50% of the datapoints had less than 90% tracking confidence. Additionally, reaches with a starting position farther than a predefined threshold near the mouse’s body were excluded to remove reaches that did not start from the typical reach initiation position (i.e. reaches that started from the pellet feeder). For included reaches, data points with less than 90% tracking confidence were replaced with interpolated values using a shape-preserving piecewise cubic Hermite interpolated polynomial. To correct tracking jumps that occurred despite high confidence, positional data points with an associated absolute velocity exceeding 1000 mm/sec were replaced with interpolated values. Positional data was smoothed using a gaussian to reduce tracking jitter due to typical high frequency, small scale tracking errors. Positional data from reach initiation to reach maximum were temporally warped as previously described^23^ to allow for the comparison of reach trajectories. Reach consistency within a session was defined as the mean pairwise Pearson coefficient for all reach trajectories in a session.

### Movement-related activity

Recorded single units were classified as movement-modulated using the *zetapy* framework^63^. *Zetapy* provides a parameter-free, timescale-independent statistical test for detecting neuronal responsiveness based on trial-to-trial variability in spike timing, without assumptions about bin width or firing rate distribution. A unit was classified as movement-modulated if its activity during the reach epoch was significantly different from baseline, defined by a *p*-value < 0.05 returned by *zetapy*. The reach epoch was defined as a 4-s window centered on the time of maximum reach extension (2 seconds before to 2 seconds after reach maximum), determined from the peristimulus time histogram (PSTH) aligned to reach peak. To ensure reliable modulation estimates and to exclude potential artifacts arising from unit dropout or drift, only trials in which a unit fired at least one spike during the reach epoch were included in the analysis. Trials with no spiking activity were excluded from that unit’s analysis. The population-averaged PSTH (see Figure 3D–F, supplementary Figure S4F for non–modulated PSTH) confirmed that these units exhibited task-related increases in firing rate, verifying the sensitivity and specificity of the *zetapy* classification in identifying reach-responsive neurons.

### Firing rate analyses across time and cortical depth

To characterize when and where neurons in motor cortex exhibited reach-related modulation, we quantified firing rates both relative to reach onset and across cortical depth. **Peri-Reach Analysis** Spike trains were aligned to reach onset and converted into peri-event time histograms (PSTHs) spanning 2 s before to 2 s after movement initiation, using 20 ms non-overlapping bins. Firing rates were z-scored for each neuron relative to its own baseline activity (–2 s to –1 s prior to reach onset). To identify statistically significant deviations in population activity, we implemented a cluster-based non-parametric permutation test. For each time bin, firing rates between groups were compared using independent-samples t-tests, yielding a t-statistic vector across time. Bins exceeding a critical threshold corresponding to *p* < 0.05 (two-sided) were grouped into contiguous temporal clusters. Each cluster’s test statistic was computed as the sum of its constituent t-values (the “cluster mass”). To determine significance at the cluster level, we constructed a null distribution by randomly permuting condition labels 1,000 times and recomputing the maximum cluster mass for each permutation. The observed cluster masses were then compared against this null distribution; clusters exceeding the 95th percentile of the null were considered significant at *p* < 0.05 (cluster-level corrected). This procedure effectively controls the family-wise error rate across multiple, correlated time bins while maintaining sensitivity to temporally contiguous effects.

### Cortical Depth Analysis

To assess how firing rate modulation varied with cortical depth, neurons were assigned depths based on probe site position and grouped into overlapping sliding windows (400 µm width, 50 µm step size). For each mouse and depth window, firing rates were averaged across neurons and then compared between control and cuprizone-treated cohorts. The same cluster-based permutation framework was applied across depth: independent-samples t-tests were computed for each depth window, significant bins were clustered based on spatial adjacency, and cluster masses were compared to a null distribution derived from 1,000 random label permutations. This approach accounts for spatial correlations between adjacent recording sites, provides a robust correction for multiple comparisons across depth, and identifies contiguous depth ranges exhibiting statistically significant differences in mean firing rate profiles between cohorts.

### Cell pairs cross-correlation analysis

Functional interactions were measured between pairs of units using cross correlograms (CCGs)^34^. The cross correlation between the spike trains over the entire approximately 60-minute recording were calculated for all possible unit pairs per mouse. To correct for firing-rate dependence, each CCG was normalized by the geometric mean of the firing rates of the pair of units, which represents the original CCG. A jitter correction method was used to remove coactivity due to movement and slow temporal correlations from the original CCG. To do this, a jittered CCG was created from the cross correlation in spike times of a resampled version of the original dataset with spike times of one unit in the pair randomly perturbed within a 50 ms jitter window. This jittered CCG was subtracted from the original CCG to obtain the jitter-corrected CCG used for all cell pair analysis^34^. A pair was determined to be a positive cell pair if the maximum of the jitter-corrected CCG amplitude in a +/- 10ms window had a magnitude larger than five-fold the standard deviation of the CCG flanks (between 50-100ms from zero). The proportion of pairs exceeding this threshold divided by the total number of possible cell pairs for each animal was defined as the proportion of cell pairs (% cell pairs).

### Joint Peristimulus Time Histogram (jPSTH)

To assess potential time-resolved coordination between simultaneously recorded neuron pairs during skilled reaching, we computed a joint peristimulus time histogram (jPSTH) for each pair. Spike trains were aligned to reach maximum for each trial and binned (5-ms) over a window spanning relative to reach maximum. For each neuron pair, a two-dimensional joint histogram was constructed by counting coincident spike occurrences across trials as a function of the relative timing of spikes in neuron A and neuron B within the same trial. Because our primary interest was in relative changes in movement-related coordination across conditions, rather than a direct estimate of intrapair interaction strength independent of rate modulation, we analyzed raw jPSTHs without any normalization. Synchrony was quantified by collapsing the jPSTH along the main diagonal. For each neuron pair, coincidence measures were normalized to the mean baseline coincidence computed during a pre-reach epoch, allowing comparisons across pairs with differing baseline firing rates. This normalized vector yields a coincidence time course that captures near-simultaneous spiking at 5- ms precision. Analyses were restricted to neuron pairs that were significantly modulated during the reaching task. jPSTHs were computed separately for each experimental condition, and population-level effects were assessed by averaging diagonal-collapsed coincidence profiles across neuron pairs. Mice were excluded if the peak coincidence value did not occur between 0.5ms from reach max (n=2 healthy control mice).

### Spiking excitatory-inhibitory network

Our cortical neural network consists of recurrently connected excitatory (i.e., *E*) and inhibitory (i.e., *I*) spiking neurons, describing the interplay between the pyramidal and interneuronal cellular populations in the deep layer of the mouse motor cortex, as well as its response to a demyelinating injury. This model captures the variability of neuronal responses, while remaining computationally and mathematically tractable^64,65^. It is further amenable to characterize the effects of changes in axonal conduction^36,37^. A schematic of the model structure and connections is presented in Figure 6A. The spiking activity of each neuron is modeled as a nonhomogeneous Poisson process:

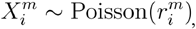

where 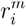 denotes the instantaneous firing rate of neuron *i* in population *m* ∈ {*E,I*}. The spike train is expressed as

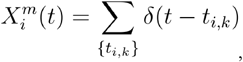

where the summation is performed over the sequence {*t*_*i,k*_}, representing the spike times for neuron *i* (Gerstner & Kistler, 2002).

The output firing rate of individual neurons, 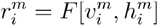, is non-linearly related to the membrane potential 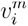 through the firing rate response function, here defined by

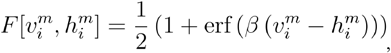

where erf refers to the error function. This sigmoid function dictates the relationship between the membrane potential and output firing rate of individual neurons. The parameter 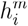 represents the firing threshold, while the gain *β* controls the steepness of the sigmoid. Together, these parameters define the excitability of neurons in the network.

The network consists of *N*^*E*^ excitatory and *N*^*I*^ inhibitory sparsely connected neurons. The dynamics of the excitatory 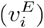 and inhibitory 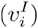 somatic membrane potentials of individual cells in the network evolve according to the set of scalar nonlinear stochastic and delayed differential equations

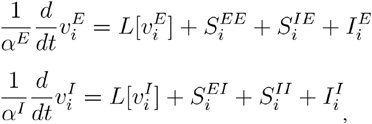

where *α*^*E*^ and *α*^*I*^ correspond to the membrane time constant for each cell type.

The dynamics in Eqs. (4-5) above are governed by the interplay between recurrent presynaptic inputs from both excitatory and inhibitory cells (i.e., 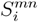), membrane leak (i.e., 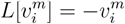) and non-synaptic inputs (i.e., 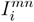), where we recall that *m,n* = *E,I*. Convergent, presynaptic inputs 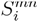 from the population *m* = *E,I* towards the th neuron of the population *n* = *E,I* are respectively given by the sum

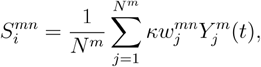

where 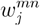 stands for the synaptic weights from neuron *j* towards neuron *i*, and 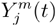 represent presynaptic spikes reaching the synaptic terminal of the postsynaptic cell. Synaptic connectivity within (i.e., 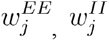) and between (i.e., 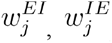) excitatory and inhibitory populations exhibit a sparse profile: synaptic weights are assigned with a value 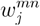 with a probability *p*^*mn*^, and set to zero otherwise. The connection probabilities and synaptic weights values were informed by experimentally measured connectivity rates and postsynaptic potential amplitudes (see below). A global scaling parameter was included to tune the relative magnitude of synaptic inputs (i.e., evenly applied to all cells across both populations), ensuring that the model resides in a dynamical regime consistent with what was observed in our data. The 1/*N*^*m*^ scaling is present so that the total synaptic input received by a cell remains invariant with network size.

The non-synaptic inputs 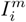 are respectively given by

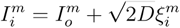

where 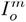 denotes a bias current imparting distinct baseline spiking rates to each cell type whereas 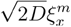 correspond to a Gaussian white noise process, accounting for fluctuations in membrane potential we do not model explicitly. The value intensity *D* was chosen so that the noise-induced fluctuations are commensurate with endogenous dynamics of the network. Such fluctuations are assumed to be independent between each cell, that is 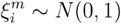, where 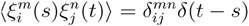 with *δ* being the Delta function. Parameter values used for the simulations are summarized in Table 1.

### Axonal propagation

To take axonal propagation into account, and model the potential effect of cuprizone, each action potential in our network is subjected to i) a conduction delay; and ii) potential propagation failure. To implement this, we relate the presynaptic spike train observed at the soma of neuron *j* (denoted by 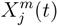, as per Eq. (2) to the corresponding spike train arriving at the synaptic terminal of postsynaptic neuron *i* after its transit along the axon (denoted by 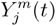). This relationship is defined as

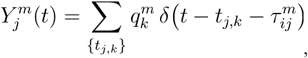

where 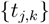 are the spike times of presynaptic neuron *j*, and 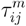 is the conduction delay from neuron *j* to neuron *i*. The 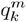 are Bernouilli random variables, that take value 1 (successful propagation) with probability *P*^*m*^, and 0 (propagation failure) with probability 1 − *P*^*m*^, for each spike *k*. As defined, the value of *P*^*m*^ reflects the probability that the *k*th spike successfully propagates along the axon. This framework allows us to rigorously define the propagation failure rate *f*^*m*^ ≡ 1 − *P*^*m*^, which represents the fraction of spikes which experience conduction block before reaching the synaptic terminal.

For instance, a propagation failure probability of *f*^*m*^ = 0 represents the healthy case, indicating that all action potentials generated by a neuron traverse faithfully towards their postsynaptic target, whereas larger values of *f*^*m*^ reflect compromised transmission: some spikes will fail reaching their destination. Note that these probabilities are cell-type specific (i.e., *m* = *E,I*).

The axonal delay 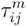 is determined by

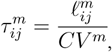

where 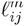 denotes the axonal length and *CV*^*m*^ the conduction velocity of population *m* ∈ {*E,I*}. Distinct conduction velocities were assigned to excitatory and inhibitory neurons, consistent with experimentally reported values (see Table 1). Axonal lengths 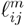 were sampled from a gamma distribution, 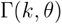, with shape parameter *k* and scale parameter θ, chosen to reproduce experimentally observed statistics (see below).

### Firing rate heterogeneity

Our data reveal a broad distribution of firing rates across neurons, both with and without cuprizone. To incorporate this heterogeneity into our model, we implemented a threshold parameter (i.e., 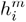), which evolves to normalize each neuron’s spontaneous firing rate. Specifically, each neuron in the network is assigned a target firing rate, toward which it converges by adjusting its excitability. These target rates are independently drawn from a lognormal distribution,

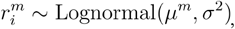

consistent with experimental observations of firing-rate heterogeneity^66^. The choice of *μ*^*m*^ and *σ* for each distribution are informed by our experimentally recorded firing data (Figure 3G). The threshold 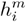, which controls the excitability of neuron (see Eq. (3)), evolves according to

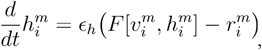

where *ϵ*_*h*_ is a slow learning rate. This mechanism ensures that neurons stabilize around their target firing rates. Note that firing rates were kept constant afterwards: in all simulations, perturbations (e.g., changes in propagation failure and/or conduction velocity; see below) were applied only after an initial transient period, allowing firing rates to first settle into their steady-state values.

### Mechanisms of demyelination

We hypothesized that cuprizone treatment disrupts neural signaling by increasing action potential propagation failure (i.e., *f*^*m*^), impairing axonal transmission of either one or both cell types in the network. We also considered an alternative mechanism in which cuprizone slows conduction velocity, lengthening propagation delays (i.e., 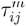).

To model propagation failure, the failure rate *f*^*m*^ in Eq. (8) was chosen to be proportional to those observed in cuprizone-treated mice for each cell type. Specifically, inhibitory failure (*f*^*I*^) is set to 30% and excitatory failure (*f*^*E*^) to 10%. The inhibitory failure rate is based on prior studies measuring action potential propagation failure in cortical PV^+^ interneurons of cuprizone-treated mice following a 6-week 0.2% cuprizone diet (Dubey et al., 2022). At this time point, demyelination is severe, so the 30% failure rate used in the model approximates the maximum biologically relevant failure rate observed experimentally. Since inhibitory neurons are preferentially myelinated and thus preferentially targeted by cuprizone, the excitatory failure rate is set at 10% to reflect the reduced vulnerability of excitatory axons to demyelination^35,67,68^. In the healthy (control) case, the failure rate for both populations is set to 0%, consistent with findings that failure is expected to be near zero at physiological temperatures^69^.

Alternatively, to model changes in conduction velocity, different values for excitatory (*cv*^*E*^) and inhibitory (*cv*^*I*^) conduction velocities are introduced following demyelination. Reference (healthy) conduction velocities of 0.5*m*/*s* for both populations are based on previous work measuring action potential conduction velocity along axons with different degrees of myelination^1,2^. To model cuprizone-induced demyelination, the conduction velocity decreases to 0.2*m*/*s* for excitatory axons and to 0.1*m*/*s* for inhibitory axons, consistent with findings that demyelination is associated with slower propagation. The greater decrease in inhibitory conduction velocity is again based on the idea that PV^+^ neurons are preferentially myelinated, consistent with the difference used when modeling propagation failure rates.

### Parameter Selection

The neuronal connectivity in our model is informed by data from the Allen Institute, which investigated the local connectivity and synaptic dynamics of neurons in the mouse primary visual cortex^70^. Specifically, synaptic physiology recordings from acute brain slices were broadly grouped into excitatory and inhibitory classes to characterize cortical connectivity and synaptic dynamics. Incorporating the results from this dataset improves the biological validity of the model, as the network connectivity reflects what is observed in the real mouse cortex. As such, to establish sparse connectivity between neurons, we randomly generate a connectivity matrix using connection probabilities for each major connection type (E-E, E-I, I-E, I-I). Although connections are randomly assigned, they are constrained by the probabilities from the Allen Institute data to preserve biological realism. The model connection probabilities (*p*^*xy*^) reflect the peak connectivity rates measured between the major cell types in the mouse primary visual cortex. These probabilities used are further adjusted to account for slicing artifacts and detection limits, as described in the original study.

Likewise, our synaptic weight values (*w*^*xy*^) are also informed by synaptic strength measurements from the Allen Institute study of the mouse primary visual cortex. That study measured connection strength as the amplitude of postsynaptic potentials (PSPs) while in a resting state, obtained using electrophysiology recordings from cortical slices^70^. Since the recordings in the original study were classified by cell subclass, the four model weight values were obtained by averaging the measured resting state EPSP/IPSP amplitudes for each major connection type. The averaged PSP values are then scaled using the dimensionless parameter *k* to ensure that the total input scales with both the number of connections and the number of neurons in the network, as stated in the model description above.

The model’s axon length distribution was selected to match the distribution profile and range of axons that observed in the mouse motor cortex. To do this, we sampled model axon lengths from a gamma distribution 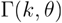 with scale parameter *k* and shape parameter *θ*. Since biological systems have latencies and conduction velocities that often follow a right-skewed distribution, the gamma distribution is often used in modeling as it captures the observed skewness while ensuring that all sampled values are positive. Since model delays are derived from axon lengths, a gamma distribution of lengths would produce a similarly skewed distribution of delays. Accordingly, the shape parameter (*k* = 2) was chosen so that the sampled distribution matches that observed experimentally. Similarly, the scale parameter (*θ* =0.05) was selected so that the mean axon length is approximately 100*μm*, consistent with recordings of intracortical axonal connections in the mouse motor cortex^71,72^.

Different time constants were chosen for E and I cells to reflect the regular spiking behavior observed in pyramidal neurons, and the fast-spiking behavior exhibited by PV^+^ interneurons. Parameter values reflecting the healthy and demyelinated case for both mechanisms of demyelination were chosen as described in the previous subsection. Key model parameters can be found in Table S1.

### Quantification and Statistical Analysis

Statistical details of experiments are reported in figure legends (including statistical tests used, exact value of n, what n represents, and exact value of p). In depth statistical details can be found in the supplemental statistics table. Individual animals are treated as biological replicates unless otherwise noted in figure legends. All data were screened for outliers using interquartile range methods. Data are all presented as mean +/- S.E.M. by animals unless otherwise noted in figure legends. No data was excluded unless specifically specified. Behavioral analysis, neural data analysis, and OL cell counting were performed blinded to the conditions of the experiments. To compare the behavioral tests in different groups, we used unpaired t-tests as appropriate and paired t-tests when comparing within group. Differences in cell pair proportions were determined using Bonferroni’s corrections for family-wise comparisons when appropriate. A two-tailed p-value < 0.05 was considered statistically significant.

## References

1. Guo, J.-Z., Graves, A.R., Guo, W.W., Zheng, J., Lee, A., Rodríguez-González, J., Li, N., Macklin, J.J., Phillips, J.W., Mensh, B.D., et al. (2015). Cortex commands the performance of skilled movement. eLife 4, e10774. 10.7554/eLife.10774.

2. Glasser, M.F., and Van Essen, D.C. (2011). Mapping Human Cortical Areas In Vivo Based on Myelin Content as Revealed by T1- and T2-Weighted MRI. J. Neurosci. 31, 11597–11616. 10.1523/JNEUROSCI.2180-11.2011.

3. Redlich, M.J., and Lim, H. (2019). A Method to Measure Myeloarchitecture of the Murine Cerebral Cortex in vivo and ex vivo by Intrinsic Third-Harmonic Generation. Front. Neuroanat. 13, 65. 10.3389/fnana.2019.00065.

4. Peters, A.J., Chen, S.X., and Komiyama, T. (2014). Emergence of reproducible spatiotemporal activity during motor learning. Nature 510, 263–267. 10.1038/nature13235.

5. Gallego, J.A., Perich, M.G., Miller, L.E., and Solla, S.A. (2017). Neural Manifolds for the Control of Movement. Neuron 94, 978– 984. 10.1016/j.neuron.2017.05.025.

6. Gibson, E.M., Purger, D., Mount, C.W., Goldstein, A.K., Lin, G.L., Wood, L.S., Inema, I., Miller, S.E., Bieri, G., Zuchero, J.B., et al. (2014). Neuronal Activity Promotes Oligodendrogenesis and Adaptive Myelination in the Mammalian Brain. Science 344, 1252304. 10.1126/science.1252304.

7. Bacmeister, C.M., Huang, R., Osso, L.A., Thornton, M.A., Conant, L., Chavez, A.R., Poleg-Polsky, A., and Hughes, E.G. (2022). Motor learning drives dynamic patterns of intermittent myelination on learning-activated axons. Nat Neurosci 25, 1300– 1313. 10.1038/s41593-022-01169-4.

8. Bacmeister, C.M., Barr, H.J., McClain, C.R., Thornton, M.A., Nettles, D., Welle, C.G., and Hughes, E.G. (2020). Motor learning promotes remyelination via new and surviving oligodendrocytes. Nat Neurosci 23, 819–831. 10.1038/s41593-020-0637-3.

9. McKenzie, I.A., Ohayon, D., Li, H., Paes De Faria, J., Emery, B., Tohyama, K., and Richardson, W.D. (2014). Motor skill learning requires active central myelination. Science 346, 318–322. 10.1126/science.1254960.

10. Carpinella, I., Cattaneo, D., and Ferrarin, M. (2014). Quantitative assessment of upper limb motor function in Multiple Sclerosis using an instrumented Action Research Arm Test. J NeuroEngineering Rehabil 11, 67. 10.1186/1743-0003-11-67.

11. Bau, L., Matas, E., Romero-Pinel, L., León, I., Muñoz-Vendrell, A., Arroyo-Pereiro, P., Martínez-Yélamos, A., and Martínez-Yélamos, S. (2025). Assessment of the Multiple Sclerosis Severity Score and the Age-Related Multiple Sclerosis Severity Score as health indicators in a population-based cohort. Neurol Sci 46, 335–342. 10.1007/s10072-024-07767-3.

12. Zuroff, L., Farkhondeh, V., Bove, R., and Green, A.J. (2025). The Road to Remyelination in Multiple Sclerosis: Breakthroughs, Challenges, and Considerations for Future Trial Design. Drugs 85, 1337–1362. 10.1007/s40265-025-02212-x.

13. Della-Flora Nunes, G., Osso, L.A., Haynes, J.A., Conant, L., Thornton, M.A., Stockton, M.E., Brassell, K.A., Morris, A., Mancha Corchado, Y.I., Gaynes, J.A., et al. (2025). Incomplete remyelination via therapeutically enhanced oligodendrogenesis is sufficient to recover visual cortical function. Nat Commun 16, 732. 10.1038/s41467-025-56092-6.

14. Gudi, V., Gingele, S., Skripuletz, T., and Stangel, M. (2014). Glial response during cuprizone-induced de- and remyelination in the CNS: lessons learned. Front. Cell. Neurosci. 8. 10.3389/fncel.2014.00073.

15. Herder, V., Hansmann, F., Stangel, M., Skripuletz, T., Baumgärtner, W., and Beineke, A. (2011). Lack of cuprizone-induced demyelination in the murine spinal cord despite oligodendroglial alterations substantiates the concept of site-specific susceptibilities of the central nervous system: Scientific correspondence. Neuropathology and Applied Neurobiology 37, 676–684. 10.1111/j.1365-2990.2011.01168.x.

16. Hamada, M.S., and Kole, M.H.P. (2015). Myelin Loss and Axonal Ion Channel Adaptations Associated with Gray Matter Neuronal Hyperexcitability. J. Neurosci. 35, 7272–7286. 10.1523/JNEUROSCI.4747-14.2015.

17. Dubey, M., Pascual-Garcia, M., Helmes, K., Wever, D.D., Hamada, M.S., Kushner, S.A., and Kole, M.H. (2022). Myelination synchronizes cortical oscillations by consolidating parvalbumin-mediated phasic inhibition. eLife 11, e73827. 10.7554/eLife.73827.

18. Moore, S., Meschkat, M., Ruhwedel, T., Trevisiol, A., Tzvetanova, I.D., Battefeld, A., Kusch, K., Kole, M.H.P., Strenzke, N., Möbius, W., et al. (2020). A role of oligodendrocytes in information processing. Nat Commun 11, 5497. 10.1038/s41467-020-19152-7.

19. Estebanez, L., Hoffmann, D., Voigt, B.C., and Poulet, J.F.A. (2017). Parvalbumin-Expressing GABAergic Neurons in Primary Motor Cortex Signal Reaching. Cell Reports 20, 308– 318. 10.1016/j.celrep.2017.06.044.

20. Thornton, M.A., Futia, G.L., Stockton, M.E., Budoff, S.A., Ramirez, A.N., Ozbay, B., Tzang, O., Kilborn, K., Poleg-Polsky, A., Restrepo, D., et al. (2024). Long-term in vivo three-photon imaging reveals region-specific differences in healthy and regenerative oligodendrogenesis. Nat Neurosci 27, 846–861. 10.1038/s41593-024-01613-7.

21. Xu, Y.K.T., Bush, A., Musheyev, E., Kim, A.A., Zhang, S., Von Bernhardi, J.E., Sulam, J., and Bergles, D.E. (2024). Brain-wide mapping of oligodendrocyte organization and oligodendrogenesis across the murine lifespan. Preprint at Neuroscience, https://doi.org/10.1101/2024.09.06.611254 10.1101/2024.09.06.611254.

22. Ludwin, S.K. (1978). Central nervous system demyelination and remyelination in the mouse: an ultrastructural study of cuprizone toxicity. Lab Invest 39, 597–612.

23. Bowles, S., Williamson, W.R., Nettles, D., Hickman, J., and Welle, C.G. (2021). Closed-loop automated reaching apparatus (CLARA) for interrogating complex motor behaviors. J. Neural Eng. 18, 045015. 10.1088/1741-2552/ac1ed1.

24. Mathis, A., Mamidanna, P., Cury, K.M., Abe, T., Murthy, V.N., Mathis, M.W., and Bethge, M. (2018). DeepLabCut: markerless pose estimation of user-defined body parts with deep learning. Nat Neurosci 21, 1281–1289. 10.1038/s41593-018-0209-y.

25. Todorov, E., and Jordan, M.I. (2002). Optimal feedback control as a theory of motor coordination. Nat Neurosci 5, 1226–1235. 10.1038/nn963.

26. Bayle, N., Lempereur, M., Hutin, E., Motavasseli, D., Remy-Neris, O., Gracies, J.-M., and Cornec, G. (2023). Comparison of Various Smoothness Metrics for Upper Limb Movements in Middle-Aged Healthy Subjects. Sensors 23, 1158. 10.3390/s23031158.

27. Hu, H., Gan, J., and Jonas, P. (2014). Fast-spiking, parvalbumin+ GABAergic interneurons: From cellular design to microcircuit function. Science 345, 1255263. 10.1126/science.1255263.

28. Stedehouder, J., Couey, J.J., Brizee, D., Hosseini, B., Slotman, J.A., Dirven, C.M.F., Shpak, G., Houtsmuller, A.B., and Kushner, S.A. (2017). Fast-spiking Parvalbumin Interneurons are Frequently Myelinated in the Cerebral Cortex of Mice and Humans. Cerebral Cortex 27, 5001–5013. 10.1093/cercor/bhx203.

29. Hooks, B.M., Mao, T., Gutnisky, D.A., Yamawaki, N., Svoboda, K., and Shepherd, G.M.G. (2013). Organization of Cortical and Thalamic Input to Pyramidal Neurons in Mouse Motor Cortex. J. Neurosci. 33, 748–760. 10.1523/JNEUROSCI.4338-12.2013.

30. Oswald, M.J., Tantirigama, M.L.S., Sonntag, I., Hughes, S.M., and Empson, R.M. (2013). Diversity of layer 5 projection neurons in the mouse motor cortex. Front. Cell. Neurosci. 7. 10.3389/fncel.2013.00174.

31. Connors, B.W., and Gutnick, M.J. (1990). Intrinsic firing patterns of diverse neocortical neurons. Trends in Neurosciences 13, 99– 104. 10.1016/0166-2236(90)90185-D.

32. Barthó, P., Hirase, H., Monconduit, L., Zugaro, M., Harris, K.D., and Buzsáki, G. (2004). Characterization of Neocortical Principal Cells and Interneurons by Network Interactions and Extracellular Features. Journal of Neurophysiology 92, 600–608. 10.1152/jn.01170.2003.

33. Sippy, T., and Yuste, R. (2013). Decorrelating Action of Inhibition in Neocortical Networks. Journal of Neuroscience 33, 9813–9830. 10.1523/JNEUROSCI.4579-12.2013.

34. Smith, M.A., and Kohn, A. (2008). Spatial and Temporal Scales of Neuronal Correlation in Primary Visual Cortex. J. Neurosci. 28, 12591–12603. 10.1523/JNEUROSCI.2929-08.2008.

35. Benamer, N., Vidal, M., Balia, M., and Angulo, M.C. (2020). Myelination of parvalbumin interneurons shapes the function of cortical sensory inhibitory circuits. Nat Commun 11, 5151. 10.1038/s41467-020-18984-7.

36. Talidou, A., Frankland, P.W., Mabbott, D., and Lefebvre, J. (2022). Homeostatic coordination and up-regulation of neural activity by activity-dependent myelination. Nat Comput Sci 2, 665–676. 10.1038/s43588-022-00315-z.

37. Lefebvre, J., Clappison, A., Longtin, A., and Hutt, A. (2025). Myelin-induced gain control in nonlinear neural networks. Commun Phys 8, 145. 10.1038/s42005-025-02055-8.

38. Hamada, M.S., Popovic, M.A., and Kole, M.H.P. (2017). Loss of Saltation and Presynaptic Action Potential Failure in Demyelinated Axons. Front. Cell. Neurosci. 11. 10.3389/fncel.2017.00045.

39. Patrikios, P., Stadelmann, C., Kutzelnigg, A., Rauschka, H., Schmidbauer, M., Laursen, H., Sorensen, P.S., Bruck, W., Lucchinetti, C., and Lassmann, H. (2006). Remyelination is extensive in a subset of multiple sclerosis patients. Brain 129, 3165–3172. 10.1093/brain/awl217.

40. Mei, F., Lehmann-Horn, K., Shen, Y.-A.A., Rankin, K.A., Stebbins, K.J., Lorrain, D.S., Pekarek, K., A Sagan, S., Xiao, L., Teuscher, C., et al. (2016). Accelerated remyelination during inflammatory demyelination prevents axonal loss and improves functional recovery. eLife 5, e18246. 10.7554/eLife.18246.

41. Cameron, M.H., and Wagner, J.M. (2011). Gait Abnormalities in Multiple Sclerosis: Pathogenesis, Evaluation, and Advances in Treatment. Curr Neurol Neurosci Rep 11, 507–515. 10.1007/s11910-011-0214-y.

42. Kalron, A., Achiron, A., Pau, M., and Cocco, E. (2020). The effect of a telerehabilitation virtual reality intervention on functional upper limb activities in people with multiple sclerosis: a study protocol for the TEAMS pilot randomized controlled trial. Trials 21, 713. 10.1186/s13063-020-04650-2.

43. Bethoux, F., and Marrie, R.A. (2016). A Cross-Sectional Study of the Impact of Spasticity on Daily Activities in Multiple Sclerosis. Patient 9, 537–546. 10.1007/s40271-016-0173-0.

44. Xu, Y.K.T., Call, C.L., Sulam, J., and Bergles, D.E. (2021). Automated in vivo Tracking of Cortical Oligodendrocytes. Front. Cell. Neurosci. 15, 667595. 10.3389/fncel.2021.667595.

45. Sauerbrei, B.A., Guo, J.-Z., Cohen, J.D., Mischiati, M., Guo, W., Kabra, M., Verma, N., Mensh, B., Branson, K., and Hantman, A.W. (2020). Cortical pattern generation during dexterous movement is input-driven. Nature 577, 386–391. 10.1038/s41586-019-1869-9.

46. Call, C.L., and Bergles, D.E. (2021). Remyelination restores myelin content on distinct neuronal subtypes in the cerebral cortex (Neuroscience) 10.1101/2021.02.17.431685.

47. Sempere-Ferràndez, A., Martínez, S., and Geijo-Barrientos, E. (2019). Synaptic mechanisms underlying the intense firing of neocortical layer 5B pyramidal neurons in response to cortico-cortical inputs. Brain Struct Funct 224, 1403–1416. 10.1007/s00429-019-01842-8.

48. Naka, A., and Adesnik, H. (2016). Inhibitory Circuits in Cortical Layer 5. Front. Neural Circuits 10. 10.3389/fncir.2016.00035.

49. Kato, D., Wake, H., Lee, P.R., Tachibana, Y., Ono, R., Sugio, S., Tsuji, Y., Tanaka, Y.H., Tanaka, Y.R., Masamizu, Y., et al. (2020). Motor learning requires myelination to reduce asynchrony and spontaneity in neural activity. Glia 68, 193–210. 10.1002/glia.23713.

50. Saab, A.S., Tzvetanova, I.D., and Nave, K.-A. (2013). The role of myelin and oligodendrocytes in axonal energy metabolism. Current Opinion in Neurobiology 23, 1065–1072. 10.1016/j.conb.2013.09.008.

51. Das, A., Bastian, C., Trestan, L., Suh, J., Dey, T., Trapp, B.D., Baltan, S., and Dana, H. (2020). Reversible Loss of Hippocampal Function in a Mouse Model of Demyelination/Remyelination. Front. Cell. Neurosci. 13, 588. 10.3389/fncel.2019.00588.

52. Crawford, D.K., Mangiardi, M., Xia, X., López-Valdés, H.E., and Tiwari-Woodruff, S.K. (2009). Functional recovery of callosal axons following demyelination: a critical window. Neuroscience 164, 1407–1421. 10.1016/j.neuroscience.2009.09.069.

53. Kolasinski, J., Stagg, C.J., Chance, S.A., DeLuca, G.C., Esiri, M.M., Chang, E.-H., Palace, J.A., McNab, J.A., Jenkinson, M., Miller, K.L., et al. (2012). A combined post-mortem magnetic resonance imaging and quantitative histological study of multiple sclerosis pathology. Brain 135, 2938–2951. 10.1093/brain/aws242.

54. Huang, R., Carter, E.R., Hughes, E.G., and Welle, C.G. (2024). Paired vagus nerve stimulation drives precise remyelination and motor recovery after myelin loss. Preprint at Neuroscience, 10.1101/2024.05.10.593609 https://doi.org/10.1101/2024.05.10.593609.

55. Hughes, E.G., Orthmann-Murphy, J.L., Langseth, A.J., and Bergles, D.E. (2018). Myelin remodeling through experience-dependent oligodendrogenesis in the adult somatosensory cortex. Nat Neurosci 21, 696–706. 10.1038/s41593-018-0121-5.

56. Gong, S., Zheng, C., Doughty, M.L., Losos, K., Didkovsky, N., Schambra, U.B., Nowak, N.J., Joyner, A., Leblanc, G., Hatten, M.E., et al. (2003). A gene expression atlas of the central nervous system based on bacterial artificial chromosomes. Nature 425, 917–925. 10.1038/nature02033.

57. Schindelin, J., Arganda-Carreras, I., Frise, E., Kaynig, V., Longair, M., Pietzsch, T., Preibisch, S., Rueden, C., Saalfeld, S., Schmid, B., et al. (2012). Fiji: an open-source platform for biological-image analysis. Nat Methods 9, 676–682. 10.1038/nmeth.2019.

58. Savage, J.T., Ramirez, J., Risher, W.C., Wang, Y., Irala, D., and Eroglu, C. (2023). SynBot: An open-source image analysis software for automated quantification of synapses. Preprint at Neuroscience, 10.1101/2023.06.26.546578 https://doi.org/10.1101/2023.06.26.546578.

59. Parslow, A., Cardona, A., and Bryson-Richardson, R.J. (2014). Sample Drift Correction Following 4D Confocal Time-lapse Imaging. JoVE, 51086. 10.3791/51086.

60. Siegle, J.H., López, A.C., Patel, Y.A., Abramov, K., Ohayon, S., and Voigts, J. (2017). Open Ephys: an open-source, plugin-based platform for multichannel electrophysiology. J. Neural Eng. 14, 045003. 10.1088/1741-2552/aa5eea.

61. Pachitariu, M., Sridhar, S., Pennington, J., and Stringer, C. (2024). Spike sorting with Kilosort4. Nat Methods 21, 914–921. 10.1038/s41592-024-02232-7.

62. Bowles, S., Hickman, J., Peng, X., Williamson, W.R., Huang, R., Washington, K., Donegan, D., and Welle, C.G. (2022). Vagus nerve stimulation drives selective circuit modulation through cholinergic reinforcement. Neuron 110, 2867-2885.e7. 10.1016/j.neuron.2022.06.017.

63. Montijn, J.S., Seignette, K., Howlett, M.H., Cazemier, J.L., Kamermans, M., Levelt, C.N., and Heimel, J.A. (2021). A parameter-free statistical test for neuronal responsiveness. eLife 10, e71969. 10.7554/eLife.71969.

64. Lefebvre, J.L., Sanes, J.R., and Kay, J.N. (2015). Development of Dendritic Form and Function. Annu. Rev. Cell Dev. Biol. 31, 741–777. 10.1146/annurev-cellbio-100913-013020.

65. Rich, S., Valiante, T.A., and Lefebvre, J. (2025). H- and m-channel overexpression promotes seizure-like events by impairing the ability of inhibitory neurons to process correlated inputs. PLoS Comput Biol 21, e1013199. 10.1371/journal.pcbi.1013199.

66. Roxin, A., Brunel, N., Hansel, D., Mongillo, G., and Van Vreeswijk, C. (2011). On the Distribution of Firing Rates in Networks of Cortical Neurons. J. Neurosci. 31, 16217–16226. 10.1523/JNEUROSCI.1677-11.2011.

67. Zonouzi, M., Berger, D., Jokhi, V., Kedaigle, A., Lichtman, J., and Arlotta, P. (2019). Individual Oligodendrocytes Show Bias for Inhibitory Axons in the Neocortex. Cell Reports 27, 2799-2808.e3. 10.1016/j.celrep.2019.05.018.

68. Basu, K., Appukuttan, S., Manchanda, R., and Sik, A. (2023). Difference in axon diameter and myelin thickness between excitatory and inhibitory callosally projecting axons in mice. Cerebral Cortex 33, 4101–4115. 10.1093/cercor/bhac329.

69. Micheva, K.D., Kiraly, M., Perez, M.M., and Madison, D.V. (2021). Conduction Velocity Along the Local Axons of Parvalbumin Interneurons Correlates With the Degree of Axonal Myelination. Cerebral Cortex 31, 3374–3392. 10.1093/cercor/bhab018.

70. Campagnola, L., Seeman, S.C., Chartrand, T., Kim, L., Hoggarth, A., Gamlin, C., Ito, S., Trinh, J., Davoudian, P., Radaelli, C., et al. (2022). Local connectivity and synaptic dynamics in mouse and human neocortex. Science 375, eabj5861. 10.1126/science.abj5861.

71. Muñoz-Castañeda, R., Zingg, B., Matho, K.S., Chen, X., Wang, Q., Foster, N.N., Li, A., Narasimhan, A., Hirokawa, K.E., Huo, B., et al. (2021). Cellular anatomy of the mouse primary motor cortex. Nature 598, 159–166. 10.1038/s41586-021-03970-w.

72. Dura-Bernal, S., Neymotin, S.A., Suter, B.A., Dacre, J., Moreira, J.V.S., Urdapilleta, E., Schiemann, J., Duguid, I., Shepherd, G.M.G., and Lytton, W.W. (2023). Multiscale model of primary motor cortex circuits predicts in vivo cell-type-specific, behavioral state-dependent dynamics. Cell Reports 42, 112574. 10.1016/j.celrep.2023.112574.

