## Supplementary material for "Myelin Supports Cortical Circuit Function Underlying Skilled Movement": TableS1_KeyModelParameters

**Table S1. Key model parameters.** The model parameter values, summarized below, are in accordance with the previous subsections.

| Parameter | Value |
|---|---|
| Number of excitatory neurons, $N^E$ | 100 |
| Number of inhibitory neurons, $N^I$ | 100 |
| Excitatory membrane time constant, $\alpha_E$ | 1 |
| Inhibitory membrane time constant, $\alpha_I$ | 2 |
| Excitatory firing rate distribution mean, $\mu^E$ | 4 Hz |
| Inhibitory firing rate distribution mean, $\mu^I$ | 6 Hz |
| Firing rate distribution standard deviation, $\sigma$ | 1.5 |
| Non-linear gain, $\beta$ | 20 |
| Noise input variance, $D$ | 0.025 |
| Excitatory-excitatory synaptic weight, $w^{EE}$ | 0.1825 mV |
| Excitatory-inhibitory synaptic weight, $w^{EI}$ | 0.3508 mV |
| Inhibitory-excitatory synaptic weight, $w^{IE}$ | -0.2809 mV |
| Inhibitory-inhibitory synaptic weight, $w^{II}$ | -0.2278 mV |
| Synaptic weight scaling, $\kappa$ | 12 |
| Excitatory-excitatory connection probability, $p^{EE}$ | 11% |
| Excitatory-inhibitory connection probability, $p^{EI}$ | 20% |
| Inhibitory-excitatory connection probability, $p^{IE}$ | 29% |
| Inhibitory-inhibitory connection probability, $p^{II}$ | 26% |
| Excitatory propagation failure rate, $f^E$ | Variable |
| Inhibitory propagation failure rate, $f^I$ | Variable |
| Excitatory conduction velocity, $cv^E$ | Variable |
| Inhibitory conduction velocity, $cv^I$ | Variable |
