## Supplementary material for "Myelin Supports Cortical Circuit Function Underlying Skilled Movement": TableS2_StatsTable

| Figure | Measure | Values | N | Statistical test | Significance |
|---|---|---|---|---|---|
| Fig. 1D | Cumul. New OLs (%). Defined as the percentage of new oligodendrocytes (as a percentage of the original population of oligodendrocytes) gained between baseline and the indicated day<br><br>Cumul. Lost OLs (%). Defined as the percentage of the original population of oligodendrocytes that was lost between baseline and the indicated day | | n=9 mice | | |
| Fig. 1G | Myelin (est. %). Defined as the estimated length of myelin sheath (as % of baseline)<br><br>Myelin reconst. (%). Defined as the myelin length (as % of baseline) measure in the myelin rescontruction | | n=3 demyelinated and 3 remyelinated timepoints from 3 mice (2 timepoints per mouse) | Standard least squares regression.<br>Run in JMP as a Fit Y by X analysis<br><br>R2 = 0.84, F(1, 4)= 21.72, p=0.0096 | |
| Fig. 1H | For all:<br>In JMP, 3P Logistic regression curves were fit to the plots for each mouse of cumulative OL loss (% of baseline) vs. day from cuprizone end - plotted in Fig. 1D; and for cumulative myelin loss (% of baseline) vs. day from cuprizone end - plotted in Fig. 1I.<br><br>Inflection point (d) for each 3P Logistic regression taken from JMP. | Inflection point - Cumul. Lost OL (d): -0.2457±1.05 (mean±SEM)<br><br>------------------------------<br><br>Inflection point - Cumul. Lost myelin (d): -5.036±0.414 (mean±SEM) | Demyelinated: 9 mice | Paired t-test:<br>t(8)=17.11, p<0.001 |  |
| Fig. 1I | New myelin (est. %). Defined as the percentage of new myelin (as a percentage of the original population of oligodendrocytes) estimated to be gained between baseline and the indicated day<br>------------------------------<br><br>Lost myelin (est. %). Defined as the percentage of the original quantity of myelin that was estimated to be lost between baseline and the indicated day | New myelin (est. %) 3d: 9.75±2 (mean±SEM)<br><br>------------------------------<br><br>Lost myelin (est. %) 3d: 67.7±4.7 (mean±SEM) | n=9 mice |  |  |
| Fig. 1J | Net demyelination (est. %). Defined as the estimated deficit in myelin (as a percentage of the original quantity) considering both myelin loss and myelin gain until the indicated day | Net demyelination (est. %) 3d: 58±3.5 (mean±SEM) | n=9 mice | | |
| Fig. 2B | success rates defined as the percentage of successful reaches out of all reach attempts for all healthy controls and cuprizone-treated demyelinated animals that reached expert-level status and underwent electrophysiology experiments at 4 days following cuprizone removal. Data points are individual mice, box and whisker plots represet median and IQR | healthy control: mean = 0.36 ± 0.04 (SEM), median = 0.33, IQR = 0.17<br>demyelinated: mean = 0.43 ± 0.04 (SEM), median = 0.41, IQR = 0.22 | n=13 healthy control mice<br>n=12 demyelinated mice<br><br>Data drawn from four batches of replicated experiments. | Shapiro-Wilk p (control) = 0.238<br>Shapiro-Wilk p (treatment) = 0.623<br>Levene's test p (equal variances) = 0.747<br><br>t-test (equal var): t=-1.208, Cohen's d=-0.504 | p=0.241 |
| Fig. 2D | Reach consistency calculated as the mean pairwise pearson correlation coefficient of all time-warped 3-dimensional reaches that passed kinematic tracking quality metrics as outlined in methods. Each data point is the within session (recording session at 4D post-cuprizone) consistency metric for each mouse. | healthy control: mean = 0.94 ± 0.01 (SEM), median = 0.94, IQR = 0.03<br>demyelinated: mean = 0.89 ± 0.02 (SEM), median = 0.89, IQR = 0.08 | n=9 healthy control mice<br>n=10 demyelinated mice<br><br>Data drawn from four batches of replicated experiments. | Shapiro-Wilk p (control) = 0.978<br>Shapiro-Wilk p (treatment) = 0.258<br>Levene's test p (equal variances) = 0.089<br><br>t-test (equal var): t=-2.437, Cohen's d=1.120 | p=0.026 |
| Fig. 2G | acceleration changes. Average number of the measured number of acceleration changes over the duration of each reach. Describes the smooth movement measurement from the post cuprizone kinematics.<br>Pre-cuprizone kinematics represent the expert reaching status on the final day of reach training, 21 days before removal of cuprizone. Post-cuprizone kinematics were collected at the time of neuropixels recording 4 days following cuprizone removal. Only a subset of mice had the pre-cuprizone kinematics collected. Mice without pre-cuprizone kinematics are excluded from this analysis. | Control PRE: 24.09 ± 1.64 (SEM)<br>Control POST: 23.34 ± 1.67 (SEM)<br>Mean Δ (post-pre): -0.75<br><br>demyelinated PRE: 21.95 ± 0.75 (SEM)<br>demyelinated POST: 25.53 ± 1.47 (SEM)<br>Mean Δ (post-pre): 3.58 | n=6 control mice<br><br>n=6 demyelinated mice<br><br>Data drawn from two batches of replicated experiments. | Control difference normality p = 0.992<br>Control: Paired t-test t=0.456,Cohen's dz=-0.172<br><br>Treatment difference normality p = 0.812<br>Treatment: Paired t-test t=-3.250, Cohen's dz=1.327 | healthy control: p=0.664, demyelinated: p=0.023 |
| Fig. 2H | Net demyelination (est. %). Defined as the estimated deficit in myelin (as a percentage of the original quantity) considering both myelin loss and myelin gain until the indicated day.<br>Rate of acceleration changes describes the smooth movement measurement from the post cuprizone kinematics. Only included mice that had both post-cuprizone kinematics and longitudinal imaging | 95% CI = [0.273, 0.978] | n=7 demyelinated mice<br><br>Data drawn from two batches of replicated experiments. | Pearson r = 0.851<br>95% CI = [0.273, 0.978] | p = 0.015 |
| 3C | Reach-modulated units were classified using zetapy python package and represent units that are either positively or negatively modulated during the reach. The %Modulated metric was calculated as the total number of reach modulated units divided by the total number of recorded units | Control: mean = 75.91 ± 3.93 (SEM), median = 76.19, IQR = 17.04<br>Treatment: mean = 76.11 ± 3.63 (SEM), median = 79.12, IQR = 15.02 | n=11 control mice<br><br>n=11 demyelinated mice<br><br>Data drawn from four batches of replicated experiments. | Shapiro-Wilk p (control) = 0.356<br>Shapiro-Wilk p (treatment) = 0.403<br>Levene's test p (equal variances) = 0.894<br><br>t-test (equal var): t=-0.036, Cohen's d=-0.015 | p=0.971 |
| 3G | The reach modulated firing rate for each unit was measured as the average number of spikes for all reach attempts over the four second reach duration. Reaches in which the unit did not spike were excluded. Only reach modulated units were included for this comparison. The firing rates for each reach modulated unit were logged (base 10) to transform the poisson rate distributions into a normal distribution for better visualization of distributions and to perform linear statistical modeling. Outliers removed if outside the interquartile range (IQR). | Control: mean = 0.49 ± 0.02 (SEM), median = 0.54, IQR = 0.67<br>Treatment: mean = 0.63 ± 0.01 (SEM), median = 0.67, IQR = 0.53<br>values in log10 | n=11 control mice, 976 units<br><br>n=11 demyelinated mice, 775 units<br><br>Data drawn from four batches of replicated experiments. | Restricted Maximum Likelihood (REML) with cohort as fixed effect and mouse ID as the random effect | p = 0.010 |
| 3H | The mean ECDF (emperical cumulative distribution function) was calculated across mice for all movement-modulated units between healthy and demyelinated mice. Measurements include the first intersection of healthy and demyelinated ECDFs and the percentile to describe where firing rate values converage along cumulative distributions of healthy and demyelinated reach modulated firing rates. | Firing rate percentiles (per-mouse, summarized by group):<br>CUP:<br>50th percentile = 4.689 ± 0.251<br>95th percentile = 16.418 ± 1.462 (n = 11)<br>cntrl:<br>50th percentile = 3.769 ± 0.269<br>95th percentile = 17.891 ± 1.439 (n = 11)<br><br>Intersection(s) where control avg ECDF equals experimental avg ECDF:<br>rate = 9.4874 Hz, percentile = 0.8088<br>rate = 21.3485 Hz, percentile = 0.9718<br>rate = 23.5771 Hz, percentile = 0.9793<br>rate = 23.7202 Hz, percentile = 0.9793<br>rate = 24.3860 Hz, percentile = 0.9812<br>rate = 26.9104 Hz, percentile = 0.9838<br>rate = 28.2817 Hz, percentile = 0.9853<br>rate = 31.3847 Hz, percentile = 0.9903<br>rate = 34.6670 Hz, percentile = 0.9910<br>rate = 40.4523 Hz, percentile = 0.9966<br>rate = 41.1548 Hz, percentile = 0.9968<br>rate = 66.3961 Hz, percentile = 1.0000 | n=11 control mice, 991 units<br><br>n=11 demyelinated mice, 787 units<br><br>Data drawn from four batches of replicated experiments. | NA | NA |

| Figure | Measure | Values | N | Statistical test | Significance |
|---|---|---|---|---|---|
| 3I | Net demyelination (est. %). Defined as the estimated deficit in myelin (as a percentage of the original quantity) considering both myelin loss and myelin gain until the indicated day.<br>Reach-modulated firing rate. Defined as the average (geometric mean) of all reach-modulated firing rates for each mouse | 95% CI = [-0.964, -0.247] | n=8 demyelinated mice<br><br>Data drawn from two batches of replicated experiments. | pearson correlation, Pearson r = -0.810 | p=0.015 |
| 3J | Reach-modulated firing rate. Defined as the average (geometric mean) of all reach-modulated firing rates for each mouse.<br>acceleration changes. Average number of the measured number of acceleration changes over the duration of each reach. Describes the smooth movement measurement from the post cuprizone kinematics. Only included mice that had both post-cuprizone kinematics and longitudinal imaging | healthy: 95% CI = [-0.313, 0.820]<br>demyelin: 95% CI = [-0.943, -0.010] | n=10 healthy mice<br>n=8 demyelinated mice<br><br>Data drawn from four batches of replicated experiments. | pearson correlation<br>healthy: r=0.39<br>demyel: r = -0.70 | healthy: p=0.25<br>demyel: p=0.049 |
| 4B | Comparison of depth distributions between cohorts while accounting for mouse-level variation. Earth Mover's Distance (EMD) values were determined from the distribution of units across depth collected for each mouse. EMD quantifies the minimal amount of distributional 'mass' that must be shifted to transform one distribution into the other, providing a measure of overall dissimilarity. Group-level differences in EMD were then assessed using a Mann–Whitney U test on per-mouse values. | control: median = -1023.274, IQR = 286.839<br>demyelinated: median = -891.567, IQR = 73.115 | n=11 control mice, 991 units<br><br>n=11 demyelinated mice, 787 units<br><br>Data drawn from four batches of replicated experiments. | Earth movers distance and mann whitney U | Mann–Whitney U test on EMD<br>U-statistic = 35.000, p-value = 0.1007 |
| 4D | Average reach modulated neural activity for regular spiking units. Spiking activity for reach modulated units was binned into 20 ms bins for the 4 second reach period and averaged for each mouse and then averaged across healthy and demyelinated mice. Periods of significant modulation during the reach were established using a cluster based permutation method (see methods) | | n=11 control mice<br><br>n=11 demyelinated mice,<br><br>Data drawn from four batches of replicated experiments. | cluster based permutation test (see methods) | see below for corrected p values for each time bin |
| 4D | p_val | | | | |
| 4D | 0.5522428781506246 | | | | |
| 4D | 0.5261777351887028 | | | | |
| 4D | 0.4219157126233852 | | | | |
| 4D | 0.3964063083160365 | | | | |
| 4D | 0.32288191015248613 | | | | |
| 4D | 0.19665081967530768 | | | | |
| 4D | 0.32288191015248613 | | | | |
| 4D | 0.32288191015248613 | | | | |
| 4D | 0.47382226481129724 | | | | |
| 4D | 0.16231813586520372 | | | | |
| 4D | 0.14671194163710516 | | | | |
| 4D | 0.11860876387689046 | | | | |
| 4D | 0.11860876387689046 | | | | |
| 4D | 0.10608233997812899 | | | | |
| 4D | 0.2557029609250582 | | | | |
| 4D | 0.19665081967530768 | | | | |
| 4D | 0.19665081967530768 | | | | |
| 4D | 0.27726520324471216 | | | | |
| 4D | 0.47382226481129724 | | | | |
| 4D | 0.215354164473551 | | | | |
| 4D | 0.34679418724535405 | | | | |
| 4D | 0.10608233997812899 | | | | |
| 4D | 0.2557029609250582 | | | | |
| 4D | 0.27726520324471216 | | | | |
| 4D | 0.27726520324471216 | | | | |
| 4D | 0.215354164473551 | | | | |
| 4D | 0.27726520324471216 | | | | |
| 4D | 0.11860876387689046 | | | | |
| 4D | 0.44775712184937544 | | | | |
| 4D | 0.5522428781506246 | | | | |
| 4D | 0.5 | | | | |
| 4D | 0.3964063083160365 | | | | |
| 4D | 0.16231813586520372 | | | | |
| 4D | 0.11860876387689046 | | | | |
| 4D | 0.29968034821451794 | | | | |
| 4D | 0.0742808874459343 | | | | |
| 4D | 0.14671194163710516 | | | | |
| 4D | 0.47382226481129724 | | | | |
| 4D | 0.37133295145984113 | | | | |
| 4D | 0.4219157126233852 | | | | |
| 4D | 0.34679418724535405 | | | | |
| 4D | 0.3964063083160365 | | | | |
| 4D | 0.3964063083160365 | | | | |
| 4D | 0.32288191015248613 | | | | |

| Figure | Measure | Values | N | Statistical test | Significance |
|---|---|---|---|---|---|
| 4D | 0.5 | | | | |
| 4D | 0.27726520324471216 | | | | |
| 4D | 0.5780842873766148 | | | | |
| 4D | 0.34679418724535405 | | | | |
| 4D | 0.32288191015248613 | | | | |
| 4D | 0.2350503799370643 | | | | |
| 4D | 0.14671194163710516 | | | | |
| 4D | 0.215354164473551 | | | | |
| 4D | 0.19665081967530768 | | | | |
| 4D | 0.2557029609250582 | | | | |
| 4D | 0.32288191015248613 | | | | |
| 4D | 0.19665081967530768 | | | | |
| 4D | 0.19665081967530768 | | | | |
| 4D | 0.16231813586520372 | | | | |
| 4D | 0.215354164473551 | | | | |
| 4D | 0.215354164473551 | | | | |
| 4D | 0.215354164473551 | | | | |
| 4D | 0.215354164473551 | | | | |
| 4D | 0.0654840518991059 | | | | |
| 4D | 0.14671194163710516 | | | | |
| 4D | 0.19665081967530768 | | | | |
| 4D | 0.038118057807282196 | | | | |
| 4D | 0.043884055915162 | | | | |
| 4D | 0.0654840518991059 | | | | |
| 4D | 0.0654840518991059 | | | | |
| 4D | 0.024422032087137148 | | | | |
| 4D | 0.0945409029822579 | | | | |
| 4D | 0.0742808874459343 | | | | |
| 4D | 0.0503338437493198 | | | | |
| 4D | 0.0654840518991059 | | | | |
| 4D | 0.16231813586520372 | | | | |
| 4D | 0.10608233997812899 | | | | |
| 4D | 0.00629305875016017 | | | | |
| 4D | 0.024422032087137148 | | | | |
| 4D | 0.0945409029822579 | | | | |
| 4D | 0.00755763467824674 | | | | |
| 4D | 0.00521953794999295 | | | | |
| 4D | 0.010773060475692903 | | | | |
| 4D | 0.00521953794999295 | | | | |
| 4D | 0.0127873249504685 | | | | |
| 4D | 0.015119510508074318 | | | | |
| 4D | 0.00521953794999295 | | | | |
| 4D | 0.0575174811473696 | | | | |
| 4D | 0.0178081820499221 | | | | |
| 4D | 0.028436518017397203 | | | | |
| 4D | 0.020894498401492264 | | | | |
| 4D | 0.0178081820499221 | | | | |
| 4D | 0.00755763467824674 | | | | |
| 4D | 0.00755763467824674 | | | | |
| 4D | 0.00521953794999295 | | | | |
| 4D | 0.0654840518991059 | | | | |
| 4D | 0.0178081820499221 | | | | |
| 4D | 0.0178081820499221 | | | | |
| 4D | 0.0178081820499221 | | | | |
| 4D | 0.00629305875016017 | | | | |
| 4D | 0.0127873249504685 | | | | |
| 4D | 0.00755763467824674 | | | | |
| 4D | 0.00290842860794023 | | | | |
| 4D | 0.010773060475692903 | | | | |
| 4D | 0.00755763467824674 | | | | |
| 4D | 0.00755763467824674 | | | | |
| 4D | 0.010773060475692903 | | | | |
| 4D | 0.032985526548328564 | | | | |
| 4D | 0.11860876387689046 | | | | |

| Figure | Measure | Values | N | Statistical test | Significance |
|---|---|---|---|---|---|
| 4D | 0.0839527378263175 | | | | |
| 4D | 0.11860876387689046 | | | | |
| 4D | 0.27726520324471216 | | | | |
| 4D | 0.14671194163710516 | | | | |
| 4D | 0.10608233997812899 | | | | |
| 4D | 0.0839527378263175 | | | | |
| 4D | 0.0503338437493198 | | | | |
| 4D | 0.11860876387689046 | | | | |
| 4D | 0.10608233997812899 | | | | |
| 4D | 0.0654840518991059 | | | | |
| 4D | 0.0503338437493198 | | | | |
| 4D | 0.0654840518991059 | | | | |
| 4D | 0.0575174811473696 | | | | |
| 4D | 0.0945409029822579 | | | | |
| 4D | 0.0742808874459343 | | | | |
| 4D | 0.10608233997812899 | | | | |
| 4D | 0.028436518017397203 | | | | |
| 4D | 0.032985526548328564 | | | | |
| 4D | 0.020894498401492264 | | | | |
| 4D | 0.0575174811473696 | | | | |
| 4D | 0.0575174811473696 | | | | |
| 4D | 0.043884055915162 | | | | |
| 4D | 0.024422032087137148 | | | | |
| 4D | 0.020894498401492264 | | | | |
| 4D | 0.020894498401492264 | | | | |
| 4D | 0.0503338437493198 | | | | |
| 4D | 0.0503338437493198 | | | | |
| 4D | 0.043884055915162 | | | | |
| 4D | 0.0178081820499221 | | | | |
| 4D | 0.0575174811473696 | | | | |
| 4D | 0.028436518017397203 | | | | |
| 4D | 0.032985526548328564 | | | | |
| 4D | 0.024422032087137148 | | | | |
| 4D | 0.038118057807282196 | | | | |
| 4D | 0.0945409029822579 | | | | |
| 4D | 0.038118057807282196 | | | | |
| 4D | 0.038118057807282196 | | | | |
| 4D | 0.028436518017397203 | | | | |
| 4D | 0.028436518017397203 | | | | |
| 4D | 0.032985526548328564 | | | | |
| 4D | 0.0178081820499221 | | | | |
| 4D | 0.0503338437493198 | | | | |
| 4D | 0.0654840518991059 | | | | |
| 4D | 0.010773060475692903 | | | | |
| 4D | 0.024422032087137148 | | | | |
| 4D | 0.028436518017397203 | | | | |
| 4D | 0.0742808874459343 | | | | |
| 4D | 0.0575174811473696 | | | | |
| 4D | 0.020894498401492264 | | | | |
| 4D | 0.024422032087137148 | | | | |
| 4D | 0.032985526548328564 | | | | |
| 4D | 0.0742808874459343 | | | | |
| 4D | 0.020894498401492264 | | | | |
| 4D | 0.11860876387689046 | | | | |
| 4D | 0.11860876387689046 | | | | |
| 4D | 0.0839527378263175 | | | | |
| 4D | 0.043884055915162 | | | | |
| 4D | 0.0178081820499221 | | | | |
| 4D | 0.020894498401492264 | | | | |
| 4D | 0.020894498401492264 | | | | |
| 4D | 0.0742808874459343 | | | | |
| 4D | 0.0178081820499221 | | | | |
| 4D | 0.038118057807282196 | | | | |
| 4D | 0.032985526548328564 | | | | |

| Figure | Measure | Values | N | Statistical test | Significance |
|---|---|---|---|---|---|
| 4D | 0.010773060475692903 | | | | |
| 4D | 0.0178081820499221 | | | | |
| 4D | 0.015119510508074318 | | | | |
| 4D | 0.015119510508074318 | | | | |
| 4D | 0.043884055915162 | | | | |
| 4D | 0.0178081820499221 | | | | |
| 4D | 0.020894498401492264 | | | | |
| 4D | 0.0503338437493198 | | | | |
| 4D | 0.0654840518991059 | | | | |
| 4D | 0.0127873249504685 | | | | |
| 4D | 0.024422032087137148 | | | | |
| 4D | 0.010773060475692903 | | | | |
| 4D | 0.028436518017397203 | | | | |
| 4D | 0.0127873249504685 | | | | |
| 4D | 0.015119510508074318 | | | | |
| 4D | 0.0503338437493198 | | | | |
| 4D | 0.0654840518991059 | | | | |
| 4D | 0.0127873249504685 | | | | |
| 4D | 0.0575174811473696 | | | | |
| 4D | 0.020894498401492264 | | | | |
| 4D | 0.0178081820499221 | | | | |
| 4D | 0.0178081820499221 | | | | |
| 4D | 0.024422032087137148 | | | | |
| 4D | 0.032985526548328564 | | | | |
| 4D | 0.0654840518991059 | | | | |
| 4D | 0.00629305875016017 | | | | |
| 4D | 0.024422032087137148 | | | | |
| 4D | 0.010773060475692903 | | | | |
| 4E | Average reach modulated neural activity for fast spiking units. Spiking activity for reach modulated units was binned into 20 ms bins for the 4 second reach period and averaged for each mouse and then averaged across healthy and demyelinated mice. Periods of significant modulation during the reach were established using a cluster based permutation method (see methods) | values listed below | n=11 control mice<br><br>n=11 demyelinated mice,<br><br>Data drawn from four batches of replicated experiments. | cluster based permutation test (see methods) | |
| 4E | p_val | | | | |
| 4E | 0.9054590970177421 | | | | |
| 4E | 0.9670144734516715 | | | | |
| 4E | 0.9257191125540657 | | | | |
| 4E | 0.934515948100894 | | | | |
| 4E | 0.9821918179500778 | | | | |
| 4E | 0.9791055015985077 | | | | |
| 4E | 0.9160472621736825 | | | | |
| 4E | 0.9561159440848379 | | | | |
| 4E | 0.9791055015985077 | | | | |
| 4E | 0.9715634819826028 | | | | |
| 4E | 0.934515948100894 | | | | |
| 4E | 0.9424825188526303 | | | | |
| 4E | 0.8813912361231095 | | | | |
| 4E | 0.8532880583628948 | | | | |
| 4E | 0.8210333150533 | | | | |
| 4E | 0.9424825188526303 | | | | |
| 4E | 0.784645835526449 | | | | |
| 4E | 0.9257191125540657 | | | | |
| 4E | 0.7442970390749418 | | | | |
| 4E | 0.893917660021871 | | | | |
| 4E | 0.893917660021871 | | | | |
| 4E | 0.9561159440848379 | | | | |
| 4E | 0.9424825188526303 | | | | |
| 4E | 0.9496661562506802 | | | | |
| 4E | 0.8532880583628948 | | | | |
| 4E | 0.9160472621736825 | | | | |
| 4E | 0.9715634819826028 | | | | |
| 4E | 0.9160472621736825 | | | | |
| 4E | 0.7442970390749418 | | | | |
| 4E | 0.934515948100894 | | | | |

| Figure | Measure | Values | N | Statistical test | Significance |
|---|---|---|---|---|---|
| 4E | 0.9715634819826028 | | | | |
| 4E | 0.8678542381330093 | | | | |
| 4E | 0.8210333150533 | | | | |
| 4E | 0.893917660021871 | | | | |
| 4E | 0.9054590970177421 | | | | |
| 4E | 0.8033491803246924 | | | | |
| 4E | 0.6771180898475139 | | | | |
| 4E | 0.9257191125540657 | | | | |
| 4E | 0.7442970390749418 | | | | |
| 4E | 0.784645835526449 | | | | |
| 4E | 0.8678542381330093 | | | | |
| 4E | 0.8813912361231095 | | | | |
| 4E | 0.934515948100894 | | | | |
| 4E | 0.8813912361231095 | | | | |
| 4E | 0.893917660021871 | | | | |
| 4E | 0.8376818641347963 | | | | |
| 4E | 0.784645835526449 | | | | |
| 4E | 0.8033491803246924 | | | | |
| 4E | 0.9257191125540657 | | | | |
| 4E | 0.7227347967552878 | | | | |
| 4E | 0.8210333150533 | | | | |
| 4E | 0.7442970390749418 | | | | |
| 4E | 0.7227347967552878 | | | | |
| 4E | 0.784645835526449 | | | | |
| 4E | 0.8033491803246924 | | | | |
| 4E | 0.8033491803246924 | | | | |
| 4E | 0.7227347967552878 | | | | |
| 4E | 0.784645835526449 | | | | |
| 4E | 0.8033491803246924 | | | | |
| 4E | 0.8678542381330093 | | | | |
| 4E | 0.8033491803246924 | | | | |
| 4E | 0.8678542381330093 | | | | |
| 4E | 0.7003196517854821 | | | | |
| 4E | 0.8033491803246924 | | | | |
| 4E | 0.7649496200629358 | | | | |
| 4E | 0.7442970390749418 | | | | |
| 4E | 0.6286670485401589 | | | | |
| 4E | 0.8210333150533 | | | | |
| 4E | 0.653205812754646 | | | | |
| 4E | 0.6035936916839635 | | | | |
| 4E | 0.653205812754646 | | | | |
| 4E | 0.5780842873766148 | | | | |
| 4E | 0.653205812754646 | | | | |
| 4E | 0.5522428781506246 | | | | |
| 4E | 0.6771180898475139 | | | | |
| 4E | 0.653205812754646 | | | | |
| 4E | 0.7003196517854821 | | | | |
| 4E | 0.7227347967552878 | | | | |
| 4E | 0.8033491803246924 | | | | |
| 4E | 0.8813912361231095 | | | | |
| 4E | 0.7227347967552878 | | | | |
| 4E | 0.784645835526449 | | | | |
| 4E | 0.653205812754646 | | | | |
| 4E | 0.6035936916839635 | | | | |
| 4E | 0.784645835526449 | | | | |
| 4E | 0.5 | | | | |
| 4E | 0.6771180898475139 | | | | |
| 4E | 0.8532880583628948 | | | | |
| 4E | 0.7649496200629358 | | | | |
| 4E | 0.6035936916839635 | | | | |
| 4E | 0.7227347967552878 | | | | |
| 4E | 0.784645835526449 | | | | |
| 4E | 0.7003196517854821 | | | | |
| 4E | 0.7003196517854821 | | | | |

| Figure | Measure | Values | N | Statistical test | Significance |
|---|---|---|---|---|---|
| 4E | 0.7442970390749418 | | | | |
| 4E | 0.7227347967552878 | | | | |
| 4E | 0.7442970390749418 | | | | |
| 4E | 0.6771180898475139 | | | | |
| 4E | 0.6771180898475139 | | | | |
| 4E | 0.6286670485401589 | | | | |
| 4E | 0.5261777351887028 | | | | |
| 4E | 0.34679418724535405 | | | | |
| 4E | 0.27726520324471216 | | | | |
| 4E | 0.29968034821451794 | | | | |
| 4E | 0.19665081967530768 | | | | |
| 4E | 0.37133295145984113 | | | | |
| 4E | 0.215354164473551 | | | | |
| 4E | 0.5522428781506246 | | | | |
| 4E | 0.5522428781506246 | | | | |
| 4E | 0.6286670485401589 | | | | |
| 4E | 0.7003196517854821 | | | | |
| 4E | 0.7227347967552878 | | | | |
| 4E | 0.6286670485401589 | | | | |
| 4E | 0.44775712184937544 | | | | |
| 4E | 0.5522428781506246 | | | | |
| 4E | 0.47382226481129724 | | | | |
| 4E | 0.32288191015248613 | | | | |
| 4E | 0.47382226481129724 | | | | |
| 4E | 0.3964063083160365 | | | | |
| 4E | 0.47382226481129724 | | | | |
| 4E | 0.3964063083160365 | | | | |
| 4E | 0.6035936916839635 | | | | |
| 4E | 0.5522428781506246 | | | | |
| 4E | 0.47382226481129724 | | | | |
| 4E | 0.47382226481129724 | | | | |
| 4E | 0.5 | | | | |
| 4E | 0.653205812754646 | | | | |
| 4E | 0.5780842873766148 | | | | |
| 4E | 0.47382226481129724 | | | | |
| 4E | 0.5780842873766148 | | | | |
| 4E | 0.44775712184937544 | | | | |
| 4E | 0.7227347967552878 | | | | |
| 4E | 0.6035936916839635 | | | | |
| 4E | 0.6286670485401589 | | | | |
| 4E | 0.6035936916839635 | | | | |
| 4E | 0.6286670485401589 | | | | |
| 4E | 0.7227347967552878 | | | | |
| 4E | 0.6771180898475139 | | | | |
| 4E | 0.5780842873766148 | | | | |
| 4E | 0.7649496200629358 | | | | |
| 4E | 0.44775712184937544 | | | | |
| 4E | 0.6035936916839635 | | | | |
| 4E | 0.5 | | | | |
| 4E | 0.6286670485401589 | | | | |
| 4E | 0.6035936916839635 | | | | |
| 4E | 0.6286670485401589 | | | | |
| 4E | 0.6771180898475139 | | | | |
| 4E | 0.7227347967552878 | | | | |
| 4E | 0.5780842873766148 | | | | |
| 4E | 0.5261777351887028 | | | | |
| 4E | 0.7227347967552878 | | | | |
| 4E | 0.653205812754646 | | | | |
| 4E | 0.5780842873766148 | | | | |
| 4E | 0.7442970390749418 | | | | |
| 4E | 0.47382226481129724 | | | | |
| 4E | 0.6771180898475139 | | | | |
| 4E | 0.6771180898475139 | | | | |
| 4E | 0.6035936916839635 | | | | |

| Figure | Measure | Values | N | Statistical test | Significance |
|---|---|---|---|---|---|
| 4E | 0.653205812754646 | | | | |
| 4E | 0.5261777351887028 | | | | |
| 4E | 0.653205812754646 | | | | |
| 4E | 0.5522428781506246 | | | | |
| 4E | 0.6035936916839635 | | | | |
| 4E | 0.6035936916839635 | | | | |
| 4E | 0.6035936916839635 | | | | |
| 4E | 0.6286670485401589 | | | | |
| 4E | 0.5261777351887028 | | | | |
| 4E | 0.7003196517854821 | | | | |
| 4E | 0.44775712184937544 | | | | |
| 4E | 0.5522428781506246 | | | | |
| 4E | 0.5780842873766148 | | | | |
| 4E | 0.653205812754646 | | | | |
| 4E | 0.47382226481129724 | | | | |
| 4E | 0.653205812754646 | | | | |
| 4E | 0.4219157126233852 | | | | |
| 4E | 0.6286670485401589 | | | | |
| 4E | 0.7227347967552878 | | | | |
| 4E | 0.7227347967552878 | | | | |
| 4E | 0.6286670485401589 | | | | |
| 4E | 0.5780842873766148 | | | | |
| 4E | 0.6286670485401589 | | | | |
| 4E | 0.5522428781506246 | | | | |
| 4E | 0.653205812754646 | | | | |
| 4E | 0.7227347967552878 | | | | |
| 4E | 0.6286670485401589 | | | | |
| 4E | 0.5780842873766148 | | | | |
| 4E | 0.6035936916839635 | | | | |
| 4E | 0.5780842873766148 | | | | |
| 4E | 0.5522428781506246 | | | | |
| 4E | 0.4219157126233852 | | | | |
| 4E | 0.653205812754646 | | | | |
| 4E | 0.653205812754646 | | | | |
| 4E | 0.5522428781506246 | | | | |
| 4E | 0.5261777351887028 | | | | |
| 4E | 0.5 | | | | |
| 4E | 0.5 | | | | |
| 4E | 0.5 | | | | |
| 4E | 0.5261777351887028 | | | | |
| 4E | 0.44775712184937544 | | | | |
| 4E | 0.5 | | | | |
| 4F | Average reach modulated firing rates across probe depth. Each unit is assigned a probe depth (see methods). Reach modulated firing rates for each regular spiking reach modulated unit were averaged within 400 um bins with a 50 micron sliding window. Periods of significant modulation during the reach were established using a cluster based permutation method (see methods) | see below for all values across depth bins | n=11 control mice<br><br>n=11 demyelinated mice,<br><br>Data drawn from four batches of replicated experiments. | cluster based permutation test (see methods) | see below |
| 4F | p_val | | | | |
| 4F | 0.811378 | | | | |
| 4F | 0.110043 | | | | |
| 4F | 0.068859 | | | | |
| 4F | 0.14 | | | | |
| 4F | 0.14 | | | | |
| 4F | 0.14 | | | | |
| 4F | 0.204514 | | | | |
| 4F | 0.128995 | | | | |
| 4F | 0.018 | | | | |
| 4F | 0.018 | | | | |
| 4F | 0.018 | | | | |
| 4F | 0.018 | | | | |
| 4F | 0.018 | | | | |
| 4F | 0.018 | | | | |
| 4F | 0.018 | | | | |
| 4F | 0.018 | | | | |

| Figure | Measure | Values | N | Statistical test | Significance |
| --- | --- | --- | --- | --- | --- |
| 4F | 0.129855 | | | | |
| 4F | 0.187883 | | | | |
| 4F | 0.391565 | | | | |
| 4F | 0.328648 | | | | |
| 4F | 0.134107 | | | | |
| 4F | 0.200287 | | | | |
| 4F | 0.227259 | | | | |
| 4F | 0.38781 | | | | |
| 4F | 0.409187 | | | | |
| 4F | 0.191034 | | | | |
| 4F | 0.186648 | | | | |
| 4G | Average reach modulated firing rates across probe depth. Each unit is assigned a probe depth (see methods). Reach modulated firing rates for each fast spiking reach modulated unit were averaged within 400 um bins with a 50 micron sliding window. Periods of significant modulation during the reach were established using a cluster based permutation method (see methods) | see below for all values across depth bins | n=11 control mice<br><br>n=11 demyelinated mice,<br><br>Data drawn from four batches of replicated experiments. | cluster based permutation test (see methods) | see below for corrected p values for each time bin |
| 4G | p_val | | | | |
| 4G | 0.498812 | | | | |
| 4G | 0.553981 | | | | |
| 4G | 0.92368 | | | | |
| 4G | 0.702974 | | | | |
| 4G | 0.390022 | | | | |
| 4G | 0.735114 | | | | |
| 4G | 0.863088 | | | | |
| 4G | 0.840174 | | | | |
| 4G | 0.814776 | | | | |
| 4G | 0.871929 | | | | |
| 4G | 0.380546 | | | | |
| 4G | 0.369225 | | | | |
| 4G | 0.262441 | | | | |
| 4G | 0.613686 | | | | |
| 4G | 0.414287 | | | | |
| 4G | 0.89705 | | | | |
| 4G | 0.631492 | | | | |
| 4G | 0.321299 | | | | |
| 4G | 0.249835 | | | | |
| 4G | 0.33132 | | | | |
| 4G | 0.209388 | | | | |
| 4G | 0.296176 | | | | |
| 4G | 0.177764 | | | | |
| 4G | 0.749043 | | | | |
| 4G | 0.987267 | | | | |
| 4G | 0.776793 | | | | |
| 4H | Net demyelination (est. %). Defined as the estimated deficit in myelin (as a percentage of the original quantity) considering both myelin loss and myelin gain until the indicated day.<br>Deep RS firing rate. Geometric mean of all regular-spiking reach-modulated firing rates below a 500 um probe depth per animal. | demyelin: 95% CI = [-0.898, 0.282] | n=8 demyelinated mice<br><br>Data drawn from two batches of replicated experiments. | pearson correlation<br>r = -0.527 | p = 0.179 |
| 4I | Net demyelination (est. %). Defined as the estimated deficit in myelin (as a percentage of the original quantity) considering both myelin loss and myelin gain until the indicated day.<br>Deep RS firing rate. Geometric mean of all regular-spiking reach-modulated firing rates below a 500 um probe depth per animal. | 95% CI =[-0.964, -0.245] | n=8 demyelinated mice<br><br>Data drawn from two batches of replicated experiments. | pearson correlation, Pearsonr = -0.810 | p=0.015 |
| 4J | acceleration changes. Average number of the measured number of acceleration changes over the duration of each reach. Describes the smooth movement measurement from the post cuprizone kinematics.<br>Deep RS firing rate. Average movement-modulated deep layer regular spiking firing rate (geometric mean) per mouse against rate of acceleration changes in acceleration | healthy: 95% CI =[-0.345, 0.808]<br>demyelin: 95% CI = [-0.803, 0.569] | n=10 healthy mice<br>n=8 demyelinated mice<br><br>Data drawn from four batches of replicated experiments. | pearson correlation<br>healthy: r=0.363<br>demyel: r = -0.227 | healthy: p=0.302<br>demyel: p=0.589 |
| 4K | acceleration changes. Average number of the measured number of acceleration changes over the duration of each reach. Describes the smooth movement measurement from the post cuprizone kinematics.<br>Deep FS firing rate. Average movement-modulated deep layer fast spiking firing rate (geometric mean) per mouse against rate of acceleration changes in acceleration | healthy: 95% CI = [-0.577, 0.677]<br>demyelin: 95% CI = [-0.969, -0.312] | n=10 healthy mice<br>n=8 demyelinated mice<br><br>Data drawn from four batches of replicated experiments. | pearson correlation<br>healthy: r = 0.082<br>demyel: r = -0.833 | healthy: p=0.821<br>demyel: p=0.010 |

| Figure | Measure | Values | N | Statistical test | Significance |
|---|---|---|---|---|---|
| 4L&M | Healthy initial reach firing rates. Initial reach z-scored activity. Average z-scored regular spiking and fast-spiking reach-related neural activity during the initial reach period (1 second before reach max). Inset includes the average activity for each mouse divided by the 1 second duration to create an average firing rate for both RS and FS over the initial reach duration. | Control: mean = 0.47 ± 0.07 (SEM), median = 0.56, IQR = 0.24<br>Treatment: mean = 0.73 ± 0.09 (SEM), median = 0.73, IQR = 0.34 | n=11 control mice<br><br>n=11 demyelinated mice<br><br>Data drawn from four batches of replicated experiments. | t-test (equal var): t=-2.387<br>Cohen's d=-1.018<br>Shapiro-Wilk p (control) = 0.734<br>Shapiro-Wilk p (treatment) = 0.719<br>Levene's test p (equal variances) = 0.557 | p=0.027 |
| 4L&M | Healthy post reach firing rates. Post reach z-scored activity. Average z-scored regular spiking and fast-spiking reach-related neural activity during the post reach period (1 second duration after reach max). Inset includes the average activity for each mouse divided by the 1 second duration to create an average firing rate for both RS and FS over the post reach duration. | Control: mean = 0.40 ± 0.07 (SEM), median = 0.38, IQR = 0.29<br>Treatment: mean = 0.50 ± 0.05 (SEM), median = 0.51, IQR = 0.15 | n=11 control mice<br><br>n=11 demyelinated mice<br><br>Data drawn from four batches of replicated experiments. | t-test (equal var): t=-1.081<br>Cohen's d=-0.472<br>Shapiro-Wilk p (control) = 0.926<br>Shapiro-Wilk p (treatment) = 0.761<br>Levene's test p (equal variances) = 0.326 | p=0.293 |
| 4L&M | Demyelinated initial reach firing rates. Initial reach z-scored activity. Average z-scored regular spiking and fast-spiking reach-related neural activity during the initial reach period (1 second before reach max). Inset includes the average activity for each mouse divided by the 1 second duration to create an average firing rate for both RS and FS over the initial reach duration. | Control: mean = 0.64 ± 0.06 (SEM), median = 0.65, IQR = 0.19<br>Treatment: mean = 0.64 ± 0.06 (SEM), median = 0.66, IQR = 0.16 | n=11 control mice<br><br>n=11 demyelinated mice<br><br>Data drawn from four batches of replicated experiments. | t-test (equal var): t=-0.039<br>Cohen's d=-0.017<br>Shapiro-Wilk p (control) = 0.830<br>Shapiro-Wilk p (treatment) = 0.621<br>Levene's test p (equal variances) = 0.721 | p=0.970 |
| 4L&M | Demyelinated post reach firing rates. Post reach z-scored activity. Average z-scored regular spiking and fast-spiking reach-related neural activity during the post reach period (1 second duration after reach max). Inset includes the average activity for each mouse divided by the 1 second duration to create an average firing rate for both RS and FS over the post reach duration. | Control: mean = 0.48 ± 0.06 (SEM), median = 0.49, IQR = 0.19<br>Treatment: mean = 0.76 ± 0.05 (SEM), median = 0.68, IQR = 0.22 | n=11 control mice<br><br>n=11 demyelinated mice<br><br>Data drawn from four batches of replicated experiments. | t-test (equal var): t=-3.566<br>Cohen's d=-1.521<br>Shapiro-Wilk p (control) = 0.901<br>Shapiro-Wilk p (treatment) = 0.316<br>Levene's test p (equal variances) = 0.825 | p=0.002 |
| 4N | RS peak to baseline. 'Peak' is the average z-scored regular spiking reach related neural activity during the 220 ms reach duration for each mouse. Baseline is the average z-scored regular-spiking reach-related neural activity during the pre-reach period (-2:-1.5 ms period aligned to reach max). The peak to baseline measurement is the average peak activity divided by the average baseline activity. | healthy: mean = 1.40 ± 0.14 (SEM), median = 1.32, IQR = 0.73<br>demyelin: mean = 1.93 ± 0.09 (SEM), median = 1.90, IQR = 0.25 | n=11 control mice<br><br>n=11 demyelinated mice<br><br>Data drawn from four batches of replicated experiments. | t-test (equal var): t=-3.007<br>Cohen's d=-1.351<br>Shapiro-Wilk p (control) = 0.832<br>Shapiro-Wilk p (treatment) = 0.870<br>Levene's test p (equal variances) = 0.069 | p=0.008 |
| 4O | FS peak to baseline. 'Peak' is the average average z-scored fast spiking reach related neural activity during the 220 ms reach duration for each mouse. Baseline is the average z-scored fast-spiking reach-related neural activity during the pre-reach period (-2:-1.5 ms period aligned to reach max). The peak to baseline measurement is the average peak activity divided by the average baseline activity. | Control:mean = 1.98 ± 0.12 (SEM), median = 1.94, IQR = 0.60<br>Treatment: mean = 2.50 ± 0.20 (SEM), median = 2.58, IQR = 0.86 | n=11 control mice<br><br>n=11 demyelinated mice<br><br>Data drawn from four batches of replicated experiments. | t-test (equal var): t=-2.237<br>Cohen's d=-0.954<br>Shapiro-Wilk p (control) = 0.141<br>Shapiro-Wilk p (treatment) = 0.608<br>Levene's test p (equal variances) = 0.136 | p=0.037 |
| 5B | Cell pairs (%). The computed proportion of cell pairs were determined as the number of total positive cell pairs divided by the total number of possible cell pairs (all combinations of recorded units) per animal. Data is presented as the percentage of this proportion (multiplied by 100). Normality of group distributions was assessed with the Shapiro–Wilk test. All groups were normally distributed, so a Welch's t-test was used. Since multiple comparisons are present, the p-values were corrected for family wise comparisons using bonferroni-holm | Control median [IQR]: 0.952 [0.191]<br>Demyelinated median [IQR]: 1.458 [0.623] | n=11 control mice<br><br>n=11 demyelinated mice<br><br>Data drawn from four batches of replicated experiments. | t-test with bonferroni-holm correction,<br>t = -3.161, raw p = 0.0069 | corrected p = 0.0208 |
| 5B | | Control median [IQR]: 0.425 [0.162]<br>Demyelinated median [IQR]: 0.598 [0.304] | | t-test with bonferroni-holm correction,<br>t = -2.190, raw p = 0.0481 | corrected p = 0.0481 |
| 5B | | Control median [IQR]: 0.527 [0.071]<br>Demyelinated median [IQR]: 0.778 [0.188] | | t-test with bonferroni-holm correction,<br>t = -2.741, raw p = 0.0184 | corrected p = 0.0367 |
| 5C | Net demyelination (est. %). Defined as the estimated deficit in myelin (as a percentage of the original quantity) considering both myelin loss and myelin gain until the indicated day.<br>Cell pairs (%). The computed proportion of cell pairs were determined as the number of total positive cell pairs divided by the total number of possible cell pairs (all combinations of recorded units) per animal. Data is presented as the percentage of this proportion (multiplied by 100). | 95% CI = [-0.702, 0.707] | n=8 demyelinated mice<br><br>Data drawn from two batches of replicated experiments. | pearson correlation, Pearson r = 0.005 | p-value = 0.991 |
| 5D | Cell pairs (%). The computed proportion of cell pairs were determined as the number of total positive cell pairs divided by the total number of possible cell pairs (all combinations of recorded units) per animal. Data is presented as the percentage of this proportion (multiplied by 100).<br>Firing rate. Defined as the geometric mean of measured overall firing rates for all collected units for each mouse. Overall firing rate for each unit was measured as the number of spikes collected over the full reacording session divided by the length of the recording session (60 minutes) | healthy: 95% CI = [-0.599, 0.601]<br>demyelinated: 95% CI = [-0.490, 0.691] | n=11 healthy mice<br>n=11 demyelinated mice<br>Data drawn from four batches of replicated experiments. | healthy: pearson r = 0.002<br>demyelinated pearson r = 0.156 | healthy: p = 0.996, demyelinated: p=0.156 |
| 5E | Peak offset times. The absolute value of the peak in latency from 0 in the cross correlation of each positive cell pair | | n=11 control mice, 578 pairs<br><br>n=11 demyelinated mice, 943 pairs<br><br>Data drawn from four batches of replicated experiments. | kolmogorov-smirnov (KS) test<br>k=0.0288 | p=0.927 |
| 5F | AUC. Area under the cross correlogram between the -10:10 ms time lags for all positive cell pairs. Measures the quality of the spike-timing precision between cell pairs. | See below Mixed Linear Model Regression Results | n=11 control mice, 578 pairs<br>n=11 demyelinated mice, 943 pairs<br><br>Data drawn from four batches of replicated experiments. | REML,<br>cohort = fixed effect<br>mouse = random effect | p=0.471 |

| Figure | Measure | Values | N | Statistical test | Significance |
|---|---|---|---|---|---|
| 5G | RSRS cell pairs (%). The computed proportion of RS & RS cell pairs were determined as the number of total positive RSRS cell pairs divided by the total number of possible RSRS cell pairs (all combinations of recorded RS and RS units) per animal. Data is presented as the percentage of this proportion (multiplied by 100).<br>Reach consistency calculated as the mean pairwise pearson correlation coefficient of all time-warped 3-dimensional reaches that passed kinematic tracking quality metrics as outlined in methods. Each data point is the within session (recording session at 4D post-cuprizone) consistency metric for each mouse. | control: 95% CI = [-0.515, 0.722]<br>demyelinated: 95% CI = [0.344, 0.961] | n=10 control mice<br><br>n=9 demyelinated mice<br><br>Data drawn from four batches of replicated experiments. | control: pearson r = 0.170<br>demyelinated pearson r = 0.820 | control: p = 0.638<br>demyelinate: p = 0.007 |
| 5J | Peak JSPTH coincidence. Joint peristimulus time histograms were calculated for all combinations of reach related units. Diagonals representing coincident spikes were measurd for each jPSTH and averaged per mouse. The peak coincidence value indicates the peak of animal's jPSTH | Control: mean = 0.08 ± 0.01 (SEM), median = 0.08, IQR = 0.04<br>Treatment: mean = 0.11 ± 0.01 (SEM), median = 0.10, IQR = 0.02 | n=11 control mice<br><br>n=10 demyelinated mice<br><br>Data drawn from four batches of replicated experiments. | Shapiro-Wilk p (treatment) = 0.101<br>Shapiro-Wilk p (control) = 0.045<br>Mann–Whitney U: U=21.000, p=0.018, rank-biserial r=0.618 | p=0.018 |
| Fig. 6C | Change in mean firing rate (Log10 Hz). Defined as the change in mean firing rate of cells in each population (E and I) when comparing healthy and demyelinated case increasing propagation delays. | E healthy: Median = 0.112800, IQR = 0.912749<br><br>I healthy: Median = 0.107085, IQR = 0.913814<br><br>E demyelinated: Median = 0.373719, IQR = 0.733459<br><br>I demyelinated: Median = 0.370595, IQR = 0.745967 | n=100 excitatory neurons, 100 inhibitory neurons (firing rates averaged over T=60s for 10 trials) | Kolmogorov-Smirnov test<br><br>p < 0.01 (*)<br>p < 0.05 (**)<br>p <0.001 (***) | |
| Fig. 6D | Change in mean firing rate by cell type (%) at varying conduction velocities. Defined as the percentage difference in firing rate from the mean of the healthy case (highlighted in gray). | NA | n=100 excitatory neurons, 100 inhibitory neurons (firing rates averaged over 10 trials of T=60s) | NA | |
| Fig. 6E | Change in mean pairwise correlation by cell type (%) at varying conduction velocities. Defined as the percentage difference in the mean Pearson correlation coefficient from the healthy case (highlighted in gray). | NA | n=100 excitatory neurons, 100 inhibitory neurons (averaged across all neurons pairs over 10 trials of T=60s) | NA | |
| Fig. 6F | Change in mean firing rate (Log10 Hz). Defined as the change in mean firing rate of cells in each population (E and I) when comparing healthy and demyelinated case increasing propagation failure. | E healthy: Median = 0.007299, IQR = 1.036723<br><br>I healthy: Median = 0.129507, IQR = 0.442664<br><br>E demyelinated: Median = 0.322570, IQR = 0.895176<br><br>I demyelinated: Median = 0.442664, IQR = 0.860172 | n=100 excitatory neurons, 100 inhibitory neurons (firing rates averaged over T=60s for 10 trials) | Kolmogorov-Smirnov test<br><br>p < 0.01 (*)<br>p < 0.05 (**)<br>p <0.001 (***) | |
| Fig. 7C | New myelin (est. %). Defined as the percentage of new myelin (as a percentage of the original population of oligodendrocytes) estimated to be gained between baseline and the indicated day<br>------------------------------<br><br>Lost myelin (est. %). Defined as the percentage of the original quantity of myelin that was estimated to be lost between baseline and the indicated day | New myelin (est. %) 24d: 39.2±5.336 (mean±SEM)<br><br>------------------------------<br><br>Lost myelin (est. %) 24d: 82.3±5.224 (mean±SEM) | n=8 mice |  |  |
| Fig. 7D | Net demyelination (est. %). Defined as the estimated deficit in myelin (as a percentage of the original quantity) considering both myelin loss and myelin gain until the indicated day | | n=8 mice | | |
| Fig. 7E | Net myelin (est. %). Defined as the estimated myelin (as a percentage of the original quantity) considering both myelin loss and myelin gain until the indicated day.<br><br>Mathematically equal to 100 - Net demyelination (ext. %) | Net myelin (est. %) remyel. (3d): 49.31.±3.622 (mean±SEM)<br><br>------------------------------<br><br>Net myelin (est. %) remyel. (24d): 56.9.±4.639 (mean±SEM) | Remyelinated: 8 mice | Paired t-test:<br>t(7)=2.592, p=0.0358 |  |
| Fig. 7F | Net myelin (est. %). Defined as the estimated myelin (as a percentage of the original quantity) considering both myelin loss and myelin gain until the indicated day.<br><br>Mathematically equal to 100 - Net demyelination (ext. %) | Net myelin (est. %) demyel. (3d): 42.01.±3.489 (mean±SEM)<br><br>------------------------------<br><br>Net myelin (est. %) remyel. (24d): 56.9.±4.639 (mean±SEM) | Demyelinated: 9 mice<br><br>------------------------------<br><br>Remyelinated: 8 mice | Unpaired t-test:<br>t(15)=2.6, p=0.02 |  |
| 7G | Reach consistency calculated as the mean pairwise pearson correlation coefficient of all time-warped 3-dimensional reaches that passed kinematic tracking quality metrics as outlined in methods. Each data point is the within session (recording session at 25D post-cuprizone) consistency metric for each mouse. | healthy: mean = 0.94 ± 0.01 (SEM), median = 0.94, IQR = 0.02<br>demyelinated: mean = 0.92 ± 0.02 (SEM), median = 0.93, IQR = 0.07 | n=5 control mice<br><br>n=7 demyelinated mice<br><br>Data drawn from two batches of replicated experiments. | Shapiro-Wilk p (control) = 0.964<br>Shapiro-Wilk p (treatment) = 0.0.331<br>Levene's test p (equal variances) = 0.224<br><br>t-test (equal var): t=-1.013, Cohen's d=0.588 | p=0.333 |
| 7H | acceleration changes. Average number of the measured number of acceleration changes over the duration of each reach. Describes the smooth movement measurement from the post cuprizone kinematics.<br>Pre-cuprizone kinematics represent the expert reaching status on the final day of reach training, 21 days before removal of cuprizone. Post-cuprizone kinematics were collected at the time of neuropixels recording 25 days following cuprizone removal. Only a subset of mice had the pre-cuprizone kinematics collected. Mice without pre-cuprizone kinematics are excluded from this analysis. | Control PRE: 25.06 ± 1.70 (SEM)<br>Control POST: 25.65 ± 2.14 (SEM)<br>Mean Δ (post-pre): 0.59<br><br>Treatment PRE: 22.73 ± 1.15 (SEM)<br>Treatment POST: 27.59 ± 1.08 (SEM)<br>Mean Δ (post-pre): 4.86 | n=5 control mice<br><br>n=5 demyelinated mice<br><br>Data drawn from two batches of replicated experiments. | healthy difference normality p = 0.704<br>Control: Paired t-test t=-0.271, Cohen's dz=0.121<br><br>demyelinated difference normality p = 0.956<br>Treatment: Paired t-test t=-3.518, Cohen's dz=1.573 | control: p=0.800, demyelinated: p=0.024 |
| 8B | The reach modulated firing rate for each unit was measured as the average number of spikes for all reach attempts over the four second reach duration. Reaches in which the unit did not spike were excluded. Only reach modulated units were included for this comparison. The firing rates for each reach modulated unit were logged (base 10) to transform the poisson rate distributions into a normal distribution for better visualization of distributions and to perform linear statistical modeling. | healthy control: mean = 0.70 ± 0.02 (SEM), median = 0.75, IQR = 0.58<br>remyelinated: mean = 0.62 ± 0.02 (SEM), median = 0.67, IQR = 0.68 | n= 349 units, 7 control mice<br><br>n= 421 units, 7 demyelinated mice<br><br>Data drawn from three batches of replicated experiments. | Restricted Maximum Likelihood (REML) with cohort as fixed effect and mouse ID as the random effect | p=0.279 |
| 8C | Net demyelination (est. %). Defined as the estimated deficit in myelin (as a percentage of the original quantity) considering both myelin loss and myelin gain until the indicated day.<br>Reach-modulated firing rate. Defined as the average (geometric mean) of all reach-modulated firing rates for each mouse | 95% CI = [-0.763, 0.943] | n=5 demyelinated mice<br><br>Data drawn from two batches of replicated experiments. | pearson correlation, Pearson r = 0.33 | p-value = 0.58 |

| Figure | Measure | Values | N | Statistical test | Significance |
|---|---|---|---|---|---|
| 8D | Cell pairs (%). The computed proportion of cell pairs were determined as the number of total positive cell pairs divided by the total number of possible cell pairs (all combinations of recorded units) per animal. Data is presented as the percentage of this proportion (multiplied by 100). Normality of group distributions was assessed with the Shapiro–Wilk test. All groups were normally distributed, so a Welch's t-test was used. Since multiple comparisons are present, the p-values were corrected for family wise comparisons using bonferroni-holm | Control median [IQR]: 1.026 [1.422]<br>Demyelinated median [IQR]: 1.836 [0.799] | n= 7 control mice<br><br>n= 7 demyelinated mice<br><br>Data drawn from three batches of replicated experiments. | Mann–Whitneywith bonferroni-holm correction, U = 15.000, raw p = 0.2593 | corrected p = 0.7780 |
| 8D | | Control median [IQR]: 0.697 [0.741]<br>Demyelinated median [IQR]: 0.814 [0.136] | | Mann–Whitney with bonferroni-holm correction,<br>U = 17.000, raw p = 0.3829 | corrected p = 0.7780 |
| 8D | | Control median [IQR]: 0.616 [0.876]<br>Demyelinated median [IQR]: 1.256 [0.758] | | Mann–Whitney with bonferroni-holm correction,<br>U = 15.000, raw p = 0.2593 | corrected p = 0.7780 |
| 8E | RSRS cell pairs (%). The computed proportion of RS & RS cell pairs were determined as the number of total positive RSRS cell pairs divided by the total number of possible RSRS cell pairs (all combinations of recorded RS and RS units) per animal. Data is presented as the percentage of this proportion (multiplied by 100).<br>Reach consistency calculated as the mean pairwise pearson correlation coefficient of all time-warped 3-dimensional reaches that passed kinematic tracking quality metrics as outlined in methods. Each data point is the within session (recording session at 25D post-cuprizone) consistency metric for each mouse. | healthy: 95% CI = [-0.811, 0.928]<br>demyel.: 95% CI = [-0.906, 0.639] | n=n=5 healthy mice<br>n=6 demyelinated mice<br>Data drawn from two batches of replicated experiments. | pearson correlation<br>healthy: r = 0.252<br>demyelinated r = -0.358 | healthy: p-value = 0.683<br>demyelin. p-value = 0.486 |
| 8F | Net demyelination (est. %). Defined as the estimated deficit in myelin (as a percentage of the original quantity) considering both myelin loss and myelin gain until the indicated day.<br>Cell pairs (%). The computed proportion of cell pairs were determined as the number of total positive cell pairs divided by the total number of possible cell pairs (all combinations of recorded units) per animal. Data is presented as the percentage of this proportion (multiplied by 100). | 95% CI = [0.355, 0.990] | n=6 demyelinated mice<br>Data drawn from two batches of replicated experiments. | pearson correlation<br>demyel: r = -0.91 | demyelin. p-value = 0.013 |
| 8H | acceleration changes. Average number of the measured number of acceleration changes over the duration of each reach. Describes the smooth movement measurement from the post cuprizone kinematics.<br>Deep RS firing rate. Average movement-modulated deep layer regular spiking firing rate (geometric mean) per mouse against rate of acceleration changes in acceleration | healthy control: 95% CI = [-0.960, 0.676]<br>demyelin: 95% CI = [-0.959, 0.686] | n=6 healthy mice<br>n=5 demyelinated mice<br>Data drawn from two batches of replicated experiments. | pearson correlation<br>healthy: r =-0.511<br>demyelinated: r =-0.497 | healthy control p = 0.379<br>demyelinated: p-value = 0.39 |
| 8I | acceleration changes. Average number of the measured number of acceleration changes over the duration of each reach. Describes the smooth movement measurement from the post cuprizone kinematics.<br>Deep FS firing rate. Average movement-modulated deep layer fast spiking firing rate (geometric mean) per mouse against rate of acceleration changes in acceleration | healthy control: 95% CI = [-0.883, 0.881]<br>demyelin: 95% CI = [0.046, 0.993] | n=6 healthy mice<br>n=5 demyelinated mice<br>Data drawn from two batches of replicated experiments. | pearson correlation<br>healthy:Pearson r = -0.004<br>demyelinated:r = 0.892 | healthy: p-value = 0.995<br>demyelinated: p-value =0.042 |
| S1B | Density of excitatory synapse colocalizations | Control: mean = 0.05 ± 0.01 (SEM), median = 0.05, IQR = 0.03<br>Treatment: mean = 0.06 ± 0.02 (SEM), median = 0.03, IQR = 0.05 | n=7 healthy control mice<br>n=5 demyelinated mice<br><br>Data drawn from four batches of replicated experiments. | t-test (equal var): t=-0.262<br>Cohen's d=-0.153<br>Shapiro-Wilk p (control) = 0.352<br>Shapiro-Wilk p (treatment) = 0.093<br>Levene's test p (equal variances) = 0.368 | p=0.799 |
| S1C | Density of inhibitory synapse colocalizations | Control: mean = 0.07 ± 0.01 (SEM), median = 0.07, IQR = 0.02<br>Treatment: mean = 0.08 ± 0.02 (SEM), median = 0.07, IQR = 0.05 | n=7 healthy control mice<br>n=7 demyelinated mice<br><br>Data drawn from four batches of replicated experiments. | t-test (equal var): t=-0.941<br>Cohen's d=-0.503<br>Shapiro-Wilk p (control) = 0.946<br>Shapiro-Wilk p (treatment) = 0.258<br>Levene's test p (equal variances) = 0.086 | p=0.365 |
| Fig. S1E | OLs/0.1 mm2: Defined as the number of MOBP-EGFP+ oligodendrocytes in an area of 0.1 mm2 of superfical M1 quantified via histology | Healthy: 279.9±2.559 (mean±SEM)<br><br>------------------------------<br><br>Demyelinated: 126.6±18.18 (mean±SEM) | Healthy: 3 mice<br><br><br>------------------------------<br><br>Demyelinated: 5 mice | Unpaired t-test:<br>t(6)=6.308, p=0.0007 |  |
| Fig. S1G | MBP (% coverage): Defined as the % of the area that is covered by MBP signal identified using automatic thresholding in Image J | Healthy: 25.61±1.23 (mean±SEM)<br><br>------------------------------<br><br>Demyelinated: 13.14±2.28 (mean±SEM) | Healthy: 4 mice<br><br><br>------------------------------<br><br>Demyelinated: 5 mice | Unpaired t-test:<br>t(7)=4.45, p=0.003 |  |
| Fig. S1H | MBP (% coverage): Defined as the % of the area that is covered by MBP signal identified using automatic thresholding in Image J<br>------------------------------<br>OLs/0.1 mm2: Defined as the number of MOBP-EGFP+ oligodendrocytes in an area of 0.1 mm2 of deep M1 quantified via histology |  | Healthy: 3 mice<br><br><br>------------------------------<br><br>Demyelinated: 5 mice | Standard least squares regression.<br>Run in JMP as a Fit Y by X analysis<br><br>R2 = 0.934, F(1, 6)= 85.06, p<0.0001 |  |
| Fig. S1J | OLs/0.1 mm2: Defined as the number of MOBP-EGFP+ oligodendrocytes in an area of 0.1 mm2 of deep M1 quantified via histology | Healthy:777.6±23.55 (mean±SEM)<br><br>------------------------------<br><br>Demyelinated: 281.6±47.36 (mean±SEM) | Healthy: 3 mice<br><br><br>------------------------------<br><br>Demyelinated: 5 mice | Unpaired t-test:<br>t(6)=7.578, p=0.0003 |  |
| Fig. S1K | OLs/0.1 mm2: Defined as the number of MOBP-EGFP+ oligodendrocytes in an area of 0.1 mm2 of M1 quantified via histology |  | Healthy: 3 mice<br><br><br>------------------------------<br><br>Demyelinated: 5 mice | Standard least squares regression.<br>Run in JMP as a Fit Y by X analysis<br><br>R2 = 0.905, F(1, 6)= 57.27, p=0.0003 |  |
| Fig. S2B | # of new sheaths. Defined as the # of sheaths produced by randomly selected new oligodendrocytes | # of new sheaths:<br>41.555±2.286% (mean±SEM) | n=9 oligodendrocytes from 3 mice | | |
| Fig. S2D | Frequency distribution of lost sheaths in relation to time of oligodendrocyte loss | | n=12 oligodendrocytes from 5 mice | | |
| Fig. S2E | Cummulative lost sheaths (%).<br>Calculated from data in Fig S1D. | | n=12 oligodendrocytes from 5 mice | 3P logistic regression:<br>Asymptote=104.49<br>Growth rate=0.264<br>Inflection point=6.49 | |

| Figure | Measure | Values | N | Statistical test | Significance |
|---|---|---|---|---|---|
| Fig. S2F | Cumul. Lost OLs 24d (%). Defined as the percentage of the original population of oligodendrocytes that was lost between baseline and day 24.<br><br>Cumul. Lost OLs 4d (%). Defined as the percentage of the original population of oligodendrocytes that was lost between baseline and day 4. | | n=8 mice | Standard least squares regression.<br>Run in JMP as a Fit Y by X analysis<br><br>R2 = 0.958, F(1, 6)= 138.46, p<0.0001 | |
| Fig. S2G | Cumul. Lost OLs (%). Defined as the percentage of the original population of oligodendrocytes that was lost between baseline and the indicated day | 3P logistic regression:<br>Predicted asymptote calculated as 11.76+1.34*Cumul. Lost OLs 4d (%), given relationship present in Fig. S1F<br><br>Growth rate and inflection point defined from average of animals in the remyelination groups as below:<br>Growth rate=0.264<br>Inflection point=6.49 | n=8 mice | | |
| RAW DATA | | | | | |
| Fig. S2I | Lost sheaths (%). Defined as the percentage of myelin sheaths that were lost by each surviving oligodendrocyte | # of new sheaths:<br>18.47±3.094% (mean±SEM) | n=10 oligodendrocytes from 3 mice | | |
| Fig. S2J | Frequency distribution of lost sheaths in surviving oligodendrocytes relation to time of end of curprizone | | n=10 oligodendrocytes from 3 mice | | |
| Fig. S2K | Cummulative lost sheaths (%).<br>Calculated from data in Fig S1J. | | n=10 oligodendrocytes from 3 mice | Probit 2P logistic regression:<br>Growth rate=0.0773<br>Inflection point=10.405 | |
| S3A | Reaches. The number of total reaches recorded for each animal during the 60 minute neuropixels recording session | healthy: mean = 73.00 ± 9.06 (SEM), median = 73.00, IQR = 30.50<br>demyelin: mean = 64.73 ± 7.85 (SEM), median = 66.00, IQR = 31.50 | n=11 healthy mice<br>n=11 demyelinated mice<br>Data drawn from four batches of replicated experiments. | t-test (equal var): t=0.690, Cohen's d=0.294<br>Shapiro-Wilk p (control) = 0.605<br>Shapiro-Wilk p (treatment) = 0.833<br>Levene's test p (equal variances) = 0.647 | p=0.498 |
| S3B | Reaches. The number of total reaches recorded for each animal during the 60 minute neuropixels recording session.<br>Net demyelination (est. %). Defined as the estimated deficit in myelin (as a percentage of the original quantity) considering both myelin loss and myelin gain until the indicated day. | 95% CI = [-0.836, 0.637] | n=7 demyelinated mice<br>Data drawn from four batches of replicated experiments. | Pearson r = -0.223 | p-value = 0.631 |
| S3C | Pathlength. 3 dimensional (x,y,z) pathlength (reach initiation to reach max) for each reach calculated (see methods) and averaged across all reaches during the 60 minute recording session for each animal | healthy: mean = 15.79 ± 0.82 (SEM), median = 15.10, IQR = 4.07<br>demyelin: mean = 14.72 ± 0.78 (SEM), median = 13.78, IQR = 2.77 | n=11 healthy mice<br>n=9 demyelinated mice<br>Data drawn from four batches of replicated experiments. | t-test (equal var): t=0.930<br>Cohen's d=0.418<br>Shapiro-Wilk p (control) = 0.311<br>Shapiro-Wilk p (treatment) = 0.508<br>Levene's test p (equal variances) = 0.577 | p=0.365 |
| S3D | Mean duration. The duration for each reach (initiation to reach max) averageed across all reaches over the 60 minute recording session per mouse | healthy: mean = 0.11 ± 0.01 (SEM), median = 0.11, IQR = 0.05<br>demyelin: mean = 0.11 ± 0.01 (SEM), median = 0.11, IQR = 0.04 | n=11 healthy mice<br>n=9 demyelinated mice<br>Data drawn from four batches of replicated experiments. | t-test (equal var): t=-0.108<br>Cohen's d=-0.049<br>Shapiro-Wilk p (control) = 0.281<br>Shapiro-Wilk p (treatment) = 0.171<br>Levene's test p (equal variances) = 0.338 | p=0.915 |
| S3E | Mean absolute velocity. Defined as the pathlength of an individual reach divided by that reach's duration. Averaged across reaches per mouse. | healthy: mean = 153.19 ± 7.29 (SEM), median = 157.06, IQR = 30.26<br>demyelin: mean = 140.73 ± 6.92 (SEM), median = 139.55, IQR = 27.19 | n=11 healthy mice<br>n=9 demyelinated mice<br>Data drawn from four batches of replicated experiments. | t-test (equal var): t=1.231<br>Cohen's d=0.598<br>Shapiro-Wilk p (control) = 0.919<br>Shapiro-Wilk p (treatment) = 0.958<br>Levene's test p (equal variances) = 0.831 | p=0.237 |
| S3F | Pathlength. 3 dimensional (x,y,z) pathlength (reach initiation to reach max) for each reach calculated (see methods) and averaged across all reaches during the 60 minute recording session for each animal<br>Net demyelination (est. %). Defined as the estimated deficit in myelin (as a percentage of the original quantity) considering both myelin loss and myelin gain until the indicated day. | 95% CI = [-0.809, 0.683] | n=8 demyelinated mice<br>Data drawn from four batches of replicated experiments. | Pearson r = -0.144 | p-value = 0.758 |
| S3G | Mean duration. The duration for each reach (initiation to reach max) averageed across all reaches over the 60 minute recording session per mouse<br>Net demyelination (est. %). Defined as the estimated deficit in myelin (as a percentage of the original quantity) considering both myelin loss and myelin gain until the indicated day. | 95% CI = [-0.674, 0.815] | n=8 demyelinated mice<br>Data drawn from four batches of replicated experiments. | Pearson r = 0.161 | p-value = 0.730 |
| S3H | Mean absolute velocity. Defined as the pathlength of an individual reach divided by that reach's duration. Averaged across reaches per mouse.<br>Net demyelination (est. %). Defined as the estimated deficit in myelin (as a percentage of the original quantity) considering both myelin loss and myelin gain until the indicated day. | 95% CI = [-0.903, 0.441] | n=8 demyelinated mice<br>Data drawn from four batches of replicated experiments. | Pearson r = -0.467 | p-value = 0.290 |
| S4A | Units. Number of units recorded across cortex that were classified as well isolated single units at the 4D timepoint | healthy: mean = 111.50 ± 12.82 (SEM), median = 114.00, IQR = 36.75<br>demyelin: mean = 99.82 ± 11.45 (SEM), median = 91.00, IQR = 44.50 | n=11 healthy mice<br>n=11 demyelin. mice<br>Data drawn from four batches of replicated experiments. | t-test (equal var): t=0.682, Cohen's d=0.298<br>Shapiro-Wilk p (control) = 0.851<br>Shapiro-Wilk p (treatment) = 0.592<br>Levene's test p (equal variances) = 0.969 | p=0.503 |
| S4B | Overall firing rate for each unit was measured as the average number of spikes for the length of the recording session. The firing rates for each unit was logged (base 10) to transform the poisson rate distributions into a normal distribution for better visualization of distributions and to perform linear statistical modeling. Outliers removed if outside the interquartile range (IQR). | healthy: mean = 0.15 ± 0.02 (SEM), median = 0.22, IQR = 0.82<br>demyelin: mean = 0.24 ± 0.02 (SEM), median = 0.32, IQR = 0.67<br>values in log10 | n=11 control mice, 1380 units<br><br>n=11 demyelinated mice, 1098 units<br><br>Data drawn from four batches of replicated experiments. | Restricted Maximum Likelihood (REML) with cohort as fixed effect and mouse ID as the random effect | p = 0.086 |
| S4C | Overall firing rate defined as the number of spikes recorded for each unit divided by the recording duration. Averaged across all recorded units per mouse. | Control: mean = 1.85 ± 0.11 (SEM), median = 1.85, IQR = 0.37<br>Treatment: mean = 2.18 ± 0.17 (SEM), median = 2.17, IQR = 0.76 | n=11 healthy mice<br>n=11 demyelin. mice<br>Data drawn from four batches of replicated experiments. | t-test (equal var): t=-1.614, Cohen's d=-0.705,<br>Shapiro-Wilk p (control) = 0.962<br>Shapiro-Wilk p (treatment) = 0.552<br>Levene's test p (equal variances) = 0.114 | p=0.123 |
| S4D | Overall firing rate defined as the number of spikes recorded for each unit divided by the recording duration. Averaged across all recorded units per mouse<br>Net demyelination (est. %). Defined as the estimated deficit in myelin (as a percentage of the original quantity) considering both myelin loss and myelin gain until the indicated day. | 95% CI = [-0.731, 0.676] | n=8 demyelinated mice<br>Data drawn from four batches of replicated experiments. | Pearson r = -0.055 | p-value = 0.898 |
| S4E | Overall firing rate defined as the number of spikes recorded for each unit divided by the recording duration. Averaged across all recorded units per mouse.<br>acceleration changes. Average number of the measured number of acceleration changes over the duration of each reach. Describes the smooth movement measurement from the post cuprizone kinematics. | healthy: 95% CI = [-0.656, 0.601]<br>demyelin: 95% CI = [-0.974, -0.392] | n=10 healthy mice<br>n=8 demyelinated mice | Pearson<br>healthy: r = -0.045<br>demyelin: r = -0.859 | healthy: p-value = 0.901<br>demyelin: p-value = 0.006 |

| Figure | Measure | Values | N | Statistical test | Significance |
|---|---|---|---|---|---|
| S4G | reach modulated firing rate. Defined as the geometric mean of the reach modulated firing rates for all reach modulated units per mouse. | Control: mean = 3.30 ± 0.14 (SEM), median = 3.26, IQR = 0.34<br>Treatment: mean = 4.08 ± 0.21 (SEM), median = 4.18, IQR = 0.90 | n=11 demyelinated mice<br>n=11 demyelinated mice<br>Data drawn from four batches of replicated experiments. | t-test (equal var): t=-2.968,<br>Cohen's d=-1.334<br>Shapiro-Wilk p (control) = 0.389<br>Shapiro-Wilk p (treatment) = 0.817<br>Levene's test p (equal variances) = 0.186 | p=0.008 |
| S4H | Reach modulated firing rate. Defined as the geometric mean of the reach modulated firing rates for all reach modulated units per mouse.<br>Pathlength. 3 dimensional (x,y,z) pathlength (reach initiation to reach max) for each reach calculated (see methods) and averaged across all reaches during the 60 minute recording session for each animal | healthy: 95% CI = [-0.568, 0.685]<br>demyelin: 95% CI = [-0.767, 0.629] | n=10 healthy mice<br>n=8 demyelin. Mice | Pearson<br>healthy r=0.1<br>demyelin r=-0.14 | healthy p=0.79<br>demyelin. p=0.75 |
| S4I | Reach modulated firing rate. Defined as the geometric mean of the reach modulated firing rates for all reach modulated units per mouse.<br>Pathlength. 3 dimensional (x,y,z) pathlength (reach initiation to reach max) for each reach calculated (see methods) and averaged across all reaches during the 60 minute recording session for each animal<br>Mean absolute velocity. Defined as the pathlength of an individual reach divided by that reach's duration. Averaged across reaches per mouse. | healthy: 95% CI = [-0.827, 0.294]<br>demyelin: 95% CI = [-0.155, 0.921] | n=10 healthy mice<br>n=8 demyelin. Mice | Pearson<br>healthy r=-0.41<br>demyelin. r=0.62 | healthy p=0.24<br>demyelin. p=0.10 |
| S4J | reach modulated firing rate. Defined as the geometric mean of the reach modulated firing rates for all reach modulated units per mouse.<br>Reach consistency calculated as the mean pairwise pearson correlation coefficient of all time-warped 3-dimensional reaches that passed kinematic tracking quality metrics as outlined in methods. Each data point is the within session (recording session at 25D post-cuprizone) consistency metric for each mouse. | healthy: 95% CI = [-0.770, 0.430]<br>demyelin: 95% CI = [-0.727, 0.681] | n=9healthy control mice<br>n=10 demyelinated mice<br><br>Data drawn from four batches of replicated experiments. | Pearson<br>healthy: r = -0.273<br>demyelin: r = -0.045 | healthy: p-value = 0.445<br>demyelin: p-value = 0.916 |
| S5A | Depth profile for one example mouse (ID: 276). Average power from 10 one-second chunks of LFP data across physiological frequency bands( delta, theta, alpha, beta, and gamma) measured for each channel. "brain surface" was determined as the channel with the greatest drop off in frequency power with agreement across bands. | Brain surface: channel 197 | n=1 example mouse | | |
| S5B | waveform duration. Time between the spike trough and subsequent peak (or full width of the spike waveform), reflecting the overall temporal width of the action potential.<br>Waveform repolarization slope. Slope of the voltage decay following the spike peak, reflecting the speed of action potential repolarization for each unit. | | n=11 control mice, 976 units<br><br>n=11 demyelinated mice, 775 units<br><br>Data drawn from four batches of replicated experiments. | | |
| S6A | Inhibitory cell pairs (%). The computed proportion of cell pairs were determined as the number of total positive inhibitory cell pairs divided by the total number of possible cell pairs (all combinations of recorded units) per animal. Data is presented as the percentage of this proportion (multiplied by 100). Inhibitory cell pairs were algorithmically determined if a trough was found 7 standard deviations below the mean of the CCG flank. Very few observations were observed in a subset of animals. | | n=5demyelinated mice<br>n=5 demyelinated mice<br>Data drawn from two batches of replicated experiments. | | |
| S6B | Deep RS firing rate. Deep RS firing rate. Average movement-modulated deep layer regular spiking firing rate (geometric mean) per mouse against rate of acceleration changes in acceleration<br>Cell pairs (%). The computed proportion of cell pairs were determined as the number of total positive cell pairs divided by the total number of possible cell pairs (all combinations of recorded units) per animal. Data is presented as the percentage of this proportion (multiplied by 100). | healthy: 95% CI = [-0.716, 0.451]<br>demyelin: 95% CI = [-0.558, 0.638] | n=11 demyelinated mice<br>n=11 demyelinated mice<br>Data drawn from four batches of replicated experiments. | Pearson<br>healthy: r = -0.204<br>demyelin: r = 0.063 | healthy: p-value = 0.548<br>demyelin: p-value = 0.855 |
| S6C | Deep FS firing rate. Deep FS firing rate. Average movement-modulated deep layer fast spiking firing rate (geometric mean) per mouse against rate of acceleration changes in acceleration<br>Cell pairs (%). The computed proportion of cell pairs were determined as the number of total positive cell pairs divided by the total number of possible cell pairs (all combinations of recorded units) per animal. Data is presented as the percentage of this proportion (multiplied by 100). | healthy: 95% CI = [-0.716, 0.451]<br>demyelin: 95% CI = [-0.558, 0.638] | n=11 demyelinated mice<br>n=11 demyelinated mice<br>Data drawn from four batches of replicated experiments. | Pearson<br>r = -0.204<br>r = 0.063 | healthy: p-value = 0.548<br>demyelin: p-value = 0.855 |
| S6D | distance between paired units. The distance in micrometers between the assigned depths within unit cell pairs. | Control: mean = 68.30 ± 1.82 (SEM), median = 44.98, IQR = 85.34<br>Treatment: mean = 69.67 ± 1.93 (SEM), median = 52.79, IQR = 75.58 | n=11 demyelinated mice<br>n=11 demyelinated mice<br>Data drawn from four batches of replicated experiments. | Restricted Maximum Likelihood Model (REML) | p=0.496 |
| S6E | Cell pairs (%). The computed proportion of cell pairs were determined as the number of total positive cell pairs divided by the total number of possible cell pairs (all combinations of recorded units) per animal. Data is presented as the percentage of this proportion (multiplied by 100).<br>acceleration changes. Average number of the measured number of acceleration changes over the duration of each reach. Describes the smooth movement measurement from the post cuprizone kinematics. | healthy: 95% CI = [-0.749, 0.472]<br>demyelin: 95% CI = [-0.531, 0.822] | n = 10 healthy mice<br>n = 8 demyelinated mice<br>Data drawn from four batches of replicated experiments. | Pearson<br>healthy: r = -0.225<br>demyelin: r = 0.278 | healthy: p-value = 0.532<br>demyelin: p-value = 0.505 |
| S6F | acceleration changes. Average number of the measured number of acceleration changes over the duration of each reach. Describes the smooth movement measurement from the post cuprizone kinematics.<br>RSRS ell pairs (%). The computed proportion of RSRS cell pairs were determined as the number of total positive RSRS cell pairs divided by the total number of possible RSRS cell pairs (all combinations of recorded units) per animal. Data is presented as the percentage of this proportion (multiplied by 100). | healthy: 95% CI = [-0.749, 0.472]<br>demyelin: 95% CI = [-0.531, 0.822] | n = 10 healthy mice<br>n = 8 demyelinated mice<br>Data drawn from four batches of replicated experiments. | Pearson<br>healthy: r = -0.225<br>demyelin: r = 0.278 | healthy: p-value = 0.532<br>demyelin: p-value = 0.505 |
| S6G | Net demyelination (est. %). Defined as the estimated deficit in myelin (as a percentage of the original quantity) considering both myelin loss and myelin gain until the indicated day<br>Peak JSPTH coincidence. Joint peristimulus time histograms were calculated for all combinations of reach related units. Diagonals representing coincident spikes were measurd for each jPSTH and averaged per mouse. The maximum value of each animal's averaged jPSTH was taken. | 95% CI = [-0.426, 0.861] | n = 8 demyelinated mice<br>Data drawn from four batches of replicated experiments. | Pearson<br>r = 0.399 | p-value = 0.328 |

| Figure | Measure | Values | N | Statistical test | Significance |
|---|---|---|---|---|---|
| S6H | Peak JSPTH coincidence. Joint peristimulus time histograms were calculated for all combinations of reach related units. Diagonals representing coincident spikes were measurd for each jPSTH and averaged per mouse. The maximum value of each animal's averaged jPSTH was taken.<br>reach modulated firing rate. Defined as the geometric mean of the reach modulated firing rates for all reach modulated units per mouse.<br>reach modulated firing rate. Defined as the geometric mean of the reach modulated firing rates for all reach modulated units per mouse. | healthy: 95% CI = [-0.751, 0.555]<br>demyelinated: 95% CI =[-0.759, 0.373] | n=9 healthy mice<br>n=11 demyelinated mice | Pearson<br>healthy: r = -0.173<br>demyelinated: r = -0.292 | healthy: p-value = 0.655<br>demyelinated: p-value = 0.384 |
| Fig. S7A | Cumul. New OLs (%). Defined as the percentage of new oligodendrocytes (as a percentage of the original population of oligodendrocytes) gained between baseline and the indicated day<br><br>Cumul. Lost OLs (%). Defined as the percentage of the original population of oligodendrocytes that was lost between baseline and the indicated day | | n=9 mice | | |
| S7B | success rates defined as the percentage of successful reaches out of all reach attempts for all healthy controls and cuprizone-treated demyelinated animals that reached expert-level status and underwent electrophysiology experiments at 4 days following cuprizone removal. Data points are individual mice, box and whisker plots represet median and IQR | healthy: mean = 41.94 ± 4.68 (SEM), median = 46.77, IQR = 14.46<br>remyelin: mean = 40.15 ± 3.22 (SEM), median = 40.19, IQR = 13.44 | n=7 healthy mice<br>n=8 remyelinated mice<br>Data drawn from two batches of replicated experiments. | t-test (equal var): t=0.323, Cohen's d=0.167<br>Shapiro-Wilk p (control) = 0.243<br>Shapiro-Wilk p (treatment) = 0.310<br>Levene's test p (equal variances) = 0.796 | p=0.752 |
| S7C | Number of reaches | Control: mean = 48.71 ± 7.49 (SEM), median = 50.00, IQR = 19.50<br>Treatment: mean = 53.75 ± 6.30 (SEM), median = 60.00, IQR = 25.75 | n=7 healthy mice<br>n=8 demyelinated mice<br>Data drawn from two batches of replicated experiments. | t-test (equal var): t=-0.519<br>Cohen's d=-0.268<br>Shapiro-Wilk p (control) = 0.985<br>Shapiro-Wilk p (treatment) = 0.469<br>Levene's test p (equal variances) = 0.858 | p=0.613 |
| S7D | Net demyelination (est. %). Defined as the estimated deficit in myelin (as a percentage of the original quantity) considering both myelin loss and myelin gain until the indicated day.<br>Rate of acceleration changes describes the smooth movement measurement from the post cuprizone kinematics. Only included mice that had both post-cuprizone kinematics and longitudinal imaging | 95% CI = [-0.935, 0.511] | n=6 demyelinated mice<br>Data drawn from two batches of replicated experiments. | Pearson<br>r=-0.513 | p=0.298 |
| S7E | Pathlength. 3 dimensional (x,y,z) pathlength (reach initiation to reach max) for each reach calculated (see methods) and averaged across all reaches during the 60 minute recording session for each animal | Control: mean = 15.13 ± 0.23 (SEM), median = 15.13, IQR = 0.92<br>Treatment: mean = 13.41 ± 0.76 (SEM), median = 14.25, IQR = 3.20 | n=5 healthy mice<br>n=7 demyelinated mice<br>Data drawn from two batches of replicated experiments. | t-test (equal var): t=1.842, Cohen's d=1.078<br>Shapiro-Wilk p (control) = 0.337<br>Shapiro-Wilk p (treatment) = 0.291<br>Levene's test p (equal variances) = 0.067 | p=0.095 |
| S7F | Mean duration. The duration for each reach (initiation to reach max) averageed across all reaches over the 60 minute recording session per mouse | healthy: mean = 0.10 ± 0.01 (SEM), median = 0.10, IQR = 0.02<br>remyelin: mean = 0.09 ± 0.01 (SEM), median = 0.09, IQR = 0.01 | n=5 healthy mice<br>n=7 demyelinated mice<br>Data drawn from two batches of replicated experiments. | t-test (equal var): t=0.222, Cohen's d=0.130<br>Shapiro-Wilk p (control) = 0.753<br>Shapiro-Wilk p (treatment) = 0.845<br>Levene's test p (equal variances) = 0.305 | p=0.829 |
| S7G | Mean absolute velocity. Defined as the pathlength of an individual reach divided by that reach's duration. Averaged across reaches per mouse. | healthy: mean = 164.46 ± 15.78 (SEM), median = 161.00, IQR = 45.78<br>remyelin: mean = 147.47 ± 3.16 (SEM), median = 145.49, IQR = 6.09 | n=5 healthy mice<br>n=7 remyelinated mice<br>Data drawn from two batches of replicated experiments. | Welch's t-test: t=1.056, Cohen's d=0.702<br>Shapiro-Wilk p (control) = 0.580<br>Shapiro-Wilk p (treatment) = 0.389<br>Levene's test p (equal variances) = 0.035 | p=0.346 |
| S8A | Units. Number of units recorded across cortex that were classified as well isolated single units at the 4D timepoint | healthy: mean = 64.43 ± 12.17 (SEM), median = 48.00, IQR = 42.50<br>remyelin: mean = 81.62 ± 9.93 (SEM), median = 76.50, IQR = 53.00 | n=7 healthy mice<br>n=8 demyelinated mice<br>Data drawn from two batches of replicated experiments. | t-test (equal var): t=-1.106, Cohen's d=-0.572<br>Shapiro-Wilk p (control) = 0.171<br>Shapiro-Wilk p (treatment) = 0.130<br>Levene's test p (equal variances) = 0.982 | p=0.289 |
| S8B | The reach modulated firing rate for each unit was measured as the average number of spikes for all reach attempts over the four second reach duration. Reaches in which the unit did not spike were excluded. Only reach modulated units were included for this comparison. The firing rates for each reach modulated unit were logged (base 10) to transform the poisson rate distributions into a normal distribution for better visualization of distributions and to perform linear statistical modeling. | healthy: mean = 0.78 ± 0.06 (SEM), median = 0.87, IQR = 0.20<br>remyelin: mean = 0.79 ± 0.03 (SEM), median = 0.79, IQR = 0.11 | n=7 control mice<br><br>n=7 remyelinated mice<br><br>Data drawn from four batches of replicated experiments. | Mann–Whitney U: U=30.000, rank-biserial r=-0.224<br>Shapiro-Wilk p (control) = 0.028<br>Shapiro-Wilk p (treatment) = 0.386<br>Levene's test p (equal variances) = 0.326 | p=0.535 |
| S8E | Average reach modulated neural activity for regular spiking units. Spiking activity for reach modulated units was binned into 20 ms bins for the 4 second reach period and averaged for each mouse and then averaged across healthy and demyelinated mice. Periods of significant modulation during the reach were established using a cluster based permutation method (see methods) | see below for values | n=11 control mice<br><br>n=11 demyelinated mice,<br><br>Data drawn from four batches of replicated experiments. | cluster based permutation test (see methods) | see below for corrected p values for each time bin |
| S8E | Average reach modulated neural activity for fast spiking units. Spiking activity for reach modulated units was binned into 20 ms bins for the 4 second reach period and averaged for each mouse and then averaged across healthy and demyelinated mice. Periods of significant modulation during the reach were established using a cluster based permutation method (see methods) | see below for values | n=11 control mice<br><br>n=11 demyelinated mice,<br><br>Data drawn from four batches of replicated experiments. | cluster based permutation test (see methods) | see below for corrected p values for each time bin |
| S8F | RS peak to baseline. 'Peak' is the average z-scored regular spiking reach related neural activity during the 220 ms reach duration for each mouse. Baseline is the average z-scored regular-spiking reach-related neural activity during the pre-reach period (-2:-1.5 ms period aligned to reach max). The peak to baseline measurement is the average peak activity divided by the average baseline activity. | Control: mean = 1.61 ± 0.18 (SEM), median = 1.50, IQR = 0.78<br>Treatment: mean = 1.88 ± 0.19 (SEM), median = 1.73, IQR = 0.49 | n=7 healthy mice<br>n=7 demyelinated mice<br>Data drawn from three batches of replicated experiments. | t-test (equal var): t=-1.047, Cohen's d=-0.559<br>Shapiro-Wilk p (control) = 0.270<br>Shapiro-Wilk p (treatment) = 0.618<br>Levene's test p (equal variances) = 0.822 | p=0.316 |
| S8F | FS peak to baseline. 'Peak' is the average z-scored fast spiking reach related neural activity during the 220 ms reach duration for each mouse. Baseline is the average z-scored fast-spiking reach-related neural activity during the pre-reach period (-2:-1.5 ms period aligned to reach max). The peak to baseline measurement is the average peak activity divided by the average baseline activity. | Control: mean = 2.97 ± 0.33 (SEM), median = 3.31, IQR = 1.30<br>Treatment: mean = 2.65 ± 0.18 (SEM), median = 2.84, IQR = 0.53 | n=7 healthy mice<br>n=7 demyelinated mice<br>Data drawn from three batches of replicated experiments. | t-test (equal var): t=0.867, Cohen's d=0.463<br>Shapiro-Wilk p (control) = 0.306<br>Shapiro-Wilk p (treatment) = 0.754<br>Levene's test p (equal variances) = 0.204 | p=0.403 |
| S8G | Average reach modulated firing rates across probe depth. Each unit is assigned a probe depth (see methods). Reach modulated firing rates for each regular spiking reach modulated unit were averaged within 400 um bins with a 50 micron sliding window. Periods of significant modulation during the reach were established using a cluster based permutation method (see methods) | see below for values across time bins | n=7 healthy mice<br>n=7 demyelinated mice<br>Data drawn from three batches of replicated experiments. | cluster-based permutation test (see methods) | |
| S8G | Average reach modulated firing rates across probe depth. Each unit is assigned a probe depth (see methods). Reach modulated firing rates for each regular spiking reach modulated unit were averaged within 400 um bins with a 50 micron sliding window. Periods of significant modulation during the reach were established using a cluster based permutation method (see methods) | see below for values across time bins | n=7 healthy mice<br>n=7 demyelinated mice<br>Data drawn from three batches of replicated experiments. | cluster-based permutation test (see methods) | |

| Figure | Measure | Values | N | Statistical test | Significance |
|---|---|---|---|---|---|
| S8H | Net demyelination (est. %). Defined as the estimated deficit in myelin (as a percentage of the original quantity) considering both myelin loss and myelin gain until the indicated day.<br>Cell type specific deep firing rate. Geometric mean of all regular-spiking or fast-spiking reach-modulated firing rates below a 500 um probe depth per animal. | RS : 95% CI = [-0.345, 0.984]<br>FS: 95% CI = [-0.966, 0.631] | n=5 demyelinated mice<br>Data drawn from two batches of replicated experiments. | pearson correlation<br>RS: r = 0.772<br>FS: r = -0.567 | RS: p-value = 0.126<br>FS: p-value = 0.319 |
